# Phenolic acid-degrading *Paraburkholderia* prime decomposition in forest soil

**DOI:** 10.1101/2020.09.28.317347

**Authors:** Roland C. Wilhelm, Christopher M. DeRito, James P. Shapleigh, Eugene L. Madsen, Daniel H. Buckley

**Affiliations:** School of Integrative Plant Science, Bradfield Hall, Cornell University, Ithaca, NY, 14853, USA; Department of Microbiology, Wing Hall, Cornell University, Ithaca, NY, 14853, USA

**Keywords:** soil ecology, carbon cycling, priming effect, *Paraburkholderia*, forest soil, stable isotope probing, *p*-hydroxybenzoic acid, *pobA*

## Abstract

Plant-derived phenolic acids are metabolized by soil microorganisms whose activity may enhance the decomposition of soil organic carbon (SOC). We characterized whether phenolic acid-degrading bacteria would enhance SOC mineralization in forest soils when primed with ^13^C-labeled *p*-hydroxybenzoic acid (PHB). We further investigated whether PHB-induced priming could explain differences in SOC content among mono-specific tree plantations in a 70-year-old common garden experiment. The activity of *Paraburkholderia* and *Caballeronia* dominated PHB degradation in all soils regardless of tree species or soil type. We isolated the principal PHB-degrading phylotype (*Paraburkholderia madseniana* RP11^T^), which encoded numerous oxidative enzymes, including secretion signal-bearing laccase, aryl-alcohol oxidase and DyP-type peroxidase, and confirmed its ability to degrade phenolics. The addition of PHB to soil led to significant enrichment (23-fold) of the RP11^T^ phylotype (RP11^ASV^), as well as enrichment of other phylotypes of *Paraburkholderia* and *Caballeronia*. Metabolism of PHB primed significant loss of SOC (3 to 13 µmols C g^-1^ dry wt soil over 7 days). In contrast, glucose addition reduced SOC mineralization (−3 to -8 µmols C g^-1^ dry wt soil over 7 days). RP11^ASV^ abundance and the expression of PHB monooxygenase (*pobA*) correlated with PHB respiration and were inversely proportional to SOC content in the field. We propose that plant-derived phenolics stimulate the activity of phenolic acid-degrading bacteria thereby causing soil priming and SOC loss. We show that *Burkholderiaceae* dominate soil priming in diverse forest soils and this observation counters the prevailing view that priming phenomena are a generalized non-specific response of community metabolism.

## Introduction

Forest soils are rich in plant-derived phenolic acids which represent a sizeable proportion of fast-cycling, low-molecular weight soil organic carbon (SOC). Phenolic acid concentrations fluctuate widely in soils (3- to 4-fold) in response to seasonal plant inputs and phenolic acid-degrading activity [1–6]. Several studies have linked the activity of phenolic acid-degrading populations to the soil priming effect [7–12], a key component of terrestrial C-cycling [13, 14]. Phenolic acids produce greater priming per unit biomass than glucose or cellobiose [10], and phenol oxidase activity positively correlates with priming [8]. The potential influence of phenolic acid-degrading activity on the persistence of SOC is suggested by the low phenolic acid content of aged SOC [15, 16]. Accordingly, we would expect to find a link between the ecology and functional capacity of phenolic acid-degrading populations, soil priming, and SOC accrual in nature. However, the identity of the microbes that mediate soil priming, and the mechanisms behind such activity, remain poorly described.

The major sources of phenolic acids to soil are litter leachate, root exudates, and the decomposition products of lignin-rich plant residues. Total phenolic concentrations in forest soils are estimated to be ∼ 0.7 % of total SOC, or between 30 – 600 ppm [5, 17–20]. The chemical composition of the phenolic pool is dependent on plant species, environmental conditions, and soil properties [21–24]. Soil phenolic composition differs broadly between coniferous and deciduous forests, with higher proportions of *p*-hydroxybenzoic (PHB) and vanillic acids in coniferous forest soils [17, 25–27], where PHB concentrations range from 0.1 and 1.4 µM [2, 28, 29]. PHB content in plant biomass positively correlates with priming of SOC [9], suggesting a link between plant matter, PHB-degrading bacteria, and the priming of SOC. Here, we examined the activity of PHB-degrading bacteria in a common garden experiment containing plantations of a coniferous (red pine) and two deciduous tree species: one leguminous (black locust) and one non-leguminous (sugar maple).

Most phenolic acid-degrading microorganisms isolated from soil are fast-growing members of the *Beta-* and *Gammaproteobacteria* (Table S1) [30–33]. Phenolics rapidly adsorb to mineral surfaces and particulate organic matter, reducing bioavailability and promoting SOC accumulation [34–37]. Hence, rapid growth may be an important life-history trait for phenolic acid-degrading microbes in soil. Though a culture-independent survey of phenolic acid-degrading microbes has yet been made in soils, metagenomic analyses of lignin degradation implicate *Alphaproteobacteria* (*Rhizobiales, Sphingomonadales* and *Caulobacterales*), *Betaproteobacteriales* (*Burkholderiaceae*) and *Actinobacteria* (*Nocardiaceae, Frankiaceae, Streptomycetaceae* and *Microbacteriaceae*) [38–40]. Populations of *Alpha*- and *Betaproteobacteriales* are also abundant during white-rot decay of wood, where phenolic acid concentrations are high [41, 42]; and in the rhizosphere, where roots exude phenolic acids to facilitate plant-microbe interactions [43, 44]. Tree roots can exude high concentrations of PHB and benzoic acid [24, 36], and benzoic acid can stimulate the growth of *Paraburkholderia* and *Caballeronia* (*Burkholderiaceae*) and prime decomposition [11]. Accordingly, we expect PHB-degraders to be relatively specialized and phylogenetically clustered relative to microbes that mediate priming from the addition of glucose [45].

The capacity to synthesize a broad range of oxidative enzymes (phenol oxidases and peroxidases) is considered one potential mechanism in soil priming [7]. PHB degradation in soil is primarily aerobic, though several classes of sulfate- or nitrate-reducing *Proteobacteria* degrade PHB anaerobically in aquatic environments [46–49]. PHB oxidation is catalyzed by *p*-hydroxybenzoate 3-monooxygenase (*pobA;* EC 1.14.13.2) as part of the peripheral pathways of aromatic degradation [50]. PobA catalyzes an essential step in the degradation of soil phenolics, and this step is rate limiting in the degradation of certain polyphenolics [51]. One of the best characterized bacteria to catabolize phenolics and polyaromatic hydrocarbons in soil is *Paraburkholderia xenovorans* LB400^T^, which encodes *pobA* along with an expansive set of peripheral pathways for degrading aromatics [52]. Notably, the expression of *pobA* was upregulated in *Burkholderia multivorans* when grown in soil versus laboratory media [53], indicating a fitness benefit in soil. The activity and functional attributes of *pobA*-encoding soil populations have yet to be studied in relation to soil priming.

We tested whether PHB primes microbial activity that enhances SOC decomposition, and whether this phenomenon could explain differences in SOC content in tree plantations in a 70-year-old common garden experiment [54]. We measured PHB degradation in the field and the ability of PHB to prime SOC mineralization in a microcosm experiment. PHB-degrading populations were characterized using 16S rRNA amplicon sequencing, stable isotope probing (SIP) with ^13^C-PHB, transcriptional analysis of *pobA*, as well as physiological and genomic characterization of PHB-degrading isolates. We hypothesized that specific populations of bacteria mediate phenolic induced priming of SOC and that their activity would explain variation in SOC content in forest soils. Specifically, we predicted (i) a direct relationship between the activity of PHB-degrading bacteria and SOC priming; (ii) PHB-degraders are phylogenetically distinct, reflecting metabolic specialization; and (iii) SOC content in forest soils is inversely proportional to PHB-degrading activity. Our microbe-centric approach supposes that the ecological and functional traits of phenolic acid-degrading bacteria are mechanistically important to soil priming and in environmental processes that govern the fate of SOC.

## Material and Methods

### Description of field site

We studied PHB-degradation and priming in soils from a 70-year old experimental forest consisting of uniform 0.4 ha plots planted with monocultures of 13 tree species spanning types typical of northeastern USA (Turkey Hill Forest Plantation, Dryden, NY) [54, 55]. The site was reforested following a period of over 100 years of agricultural use. We sampled plantations of sugar maple (“SM,” *Acer saccharum* Marsh; 42.451137, -76.420519), red pine (“RP,” *Pinus resinosa* Ait.; 42.450945, -76.420638), and black locust (“BL,” *Robinia pseudoacacia* L.; 42.451818, -76.420614) grown in typic Fragiochrept soils (inceptisols) belonging to either the Mardin B (“MaB”; Typic Fragiudepts) or Lordstown channery silt loam C soil series (“LnC”; Typic Dystrudepts). Details on the site and three separate field sampling campaigns are provided in the Supplementary Methods. Our study examined a total of five combinations of tree species + soil type, which we have termed ‘ecoplots,’ that encompassed a gradient in total soil organic matter: BL_MaB_ (21.9% w/w) > SM_MaB_ (14.5%) > BL_LnC_ (14.0%) > RP_LnC_ (12.75%) > SM_LnC_ (10.2%). There were no plantations of RP in MaB soil. The concentration of bioavailable elements in soils was determined using the modified Morgan extraction procedure [56] by the Cornell Nutrient Analysis Laboratory. Estimates of soil organic matter were made according to the loss-on-ignition method.

*PHB degradation in field soil (in situ)*

PHB mineralization rates were measured in field soils for every ecoplot. Three sampling plots (20 × 20 cm) were randomly selected per ecoplot. The litter layer was cleared to expose the O_a_ / A-horizon soil and stainless-steel chambers (radius 1.25 cm) were driven into the soil at an even depth (2 cm). Photographs of field sampling are provided (Figure S1). Soils were amended with 150 µl of either ^13^C_7_-labeled *p*-hydroxybenzoic acid (99% atom ^13^C; Sigma-Aldrich) or unlabeled control (‘^12^C’; Sigma-Aldrich) at a concentration of 1,000 ppm (6.9 and 7.3 mM, respectively). Chambers were immediately enclosed with rubber septa and ^13^CO_2_ was sampled five times over a period of 27 hrs. At each time point, 2.5 mL of headspace was sampled and stored in evacuated 2-mL vials prior to analysis with a Hewlett Packard 5890 gas chromatograph (Wilmington, DE) equipped with a 5971A mass selective detector (details in Supplementary Methods). Net ^13^CO_2_ was determined by subtracting average natural abundance ^13^CO_2_ in ^12^C controls from the total ^13^CO_2_ respired in ^13^C-PHB-amended soils. Following the incubation period, approximately 0.5 g of soil was collected from the upper 1 cm of the dosed soil, transported on ice and stored at -80°C for use in DNA-SIP.

### PHB-induced soil priming

Soil microcosms were prepared with soil sampled from the field to link the activity of phenolic acid-degrading populations with soil priming. Soils were subjected to seven treatments: (i) water-only (control), (ii) ^13^C-glucose, (iii) ^13^C-PHB, (iv) ^13^C-PHB + PobA inhibitor, (v) PobA inhibitor-only (control), (vi) ^13^C-labeled RP11^T^ cells + dilute ^12^C-PHB, and (vii) unlabeled RP11^T^ cells + dilute ^13^C-PHB (schematic overview in Figure S2, details provided below). Fresh soil was collected from the entire upper 5 cm of a 20 × 20 cm sampling plot for all LnC soils, sieved to remove root and litter (2-mm), homogenized, and pre-incubated for 10 days. Sieving, homogenization, and pre-incubation were necessary to reduce soil SOC heterogeneity and limit variability in the concentration of labile SOC at the time of sampling. After pre-incubation, soils were air dried for 48 hrs in a sterile biosafety hood. Ten grams of dry soil were added to 120-mL serum vials and wetted during treatment application to 60% water holding capacity. Glucose and PHB were added at a concentration of 0.5 mg C per g dry wt. soil. Each treatment was run in quadruplicate for each ecoplot. Isotopically labeled substrates (17.5 atom % ^13^C) or RP11^T^ cells (99 atom % ^13^C) were used to distinguish between the CO_2_ derived from amendment versus SOC. The isotopic composition of amendments was confirmed by EA-IRMS by the Cornell Stable Isotope Laboratory. RP11^T^ cells were grown on either ^13^C-glucose (99% atom ^13^C; Sigma-Aldrich) or unlabeled glucose and re-suspended in 0.85% NaCl buffer (1.5 × 10^9^ cells per microcosm) with dilute PHB (25 µg C per g dry wt. soil) to stimulate PHB-degrading activity prior to amendment. Headspace sampling was performed every 24 hrs for a week with the headspace exchanged with filtered air after each sampling. The quantity of ^12^CO_2_ (m/z 44) and ^13^CO_2_ (m/z 45) was analyzed using GC/MS (Shimadzu GCMS-QP2010S) with a set of standards ranging from 1000 – 40 000 ppm CO_2_. PobA activity was inhibited with methyl 4-hydroxy-3-iodobenzoate (Fisher Scientific) at a saturating concentration (0.48 mM), which fully inhibited growth of RP11^T^ for 72 hrs in mineral salts media containing 25 mM PHB as the sole carbon source (see Supplementary Results). After seven days, sub-samples of 0.5 g of wet wt. soil were stored at -80°C for use in RNA-based quantification of *pobA* gene abundance and determination of bacterial community composition. Headspace CO_2_ measurements continued at reduced frequency for a total of 26 days. Additional details are available in the Supplementary Methods.

### Identifying PHB-degrading populations

DNA and RNA were extracted from 0.5 g of soil using the modified phenol-chloroform method from [57] following bead-beating at 5.5 m·s^-1^ for 2.5 min (Bio Spec Products, Santa Clara, CA, USA) in 2-mL Lysing Matrix E Tubes (MP Biomedicals, OH). For the DNA-SIP experiment, DNA was extracted from ^13^C- and ^12^C-PHB amended field soils and ^13^C-enriched DNA was separated from unlabeled DNA by CsCl density gradient ultracentrifugation as previously described [58, 59]. For additional details on nucleic acid extractions and DNA-SIP methods see Supplementary Methods. Twenty 200-µL fractions were collected from each CsCl gradient. 16S rRNA gene amplicon sequencing libraries were prepared for fractions F_1-2_ (pooled), F_3_, F_4_, F_5_, F_6_, F_7_, F_8_, F_9-10_ and F_11-12_ (spanning CsCl buoyant density of 1.77 - 1.70 g·mL^-1^). For *in situ pobA* mRNA profiling and soil priming experiments, DNA and RNA were extracted concurrently from PHB-amended and unamended field soil. Complementary DNA (cDNA) was synthesized using SunScript reverse transcriptase (Expedeon, San Diego USA) to reverse transcribe DNAse-treated RNA extracts for use in qPCR (*pobA*) and 16S rRNA amplicon libraries (details in Supplementary Methods). Bacterial community composition was assessed by amplification of the V4 region using polymerase chain reaction (PCR) with dual-indexed barcoded 515f/806r primers as previously described [60]. PCR was performed in duplicate and pooled, and PCR products were purified and normalized to a standard concentration prior to sequencing on three lanes of Illumina MiSeq (2×250 paired end) at the Cornell Biotechnology Resource Centre. Amplicon libraries were deposited in the European Sequence Archive under the BioProject accession PRJEB23740. Following sequencing processing (described below), amplicon sequence variants (phylotypes) were designated as PHB-degraders when their relative abundance was at least 8-fold greater in heavy CsCl fractions (F_3_-F_8_) in ^13^C-enriched versus ^12^C-control amplicon libraries, after variance stabilization by DESeq2 (v. 1.24.0) [61]. Phylotypes increasing in response to substrate additions in the priming experiment were identified by indicator species analyses using the R package ‘indicspecies’ [62]. Full details on methods and controls for DNA extractions, cDNA synthesis, PCR conditions and isopycnic gradient centrifugation are available in the Supplementary Methods.

### Isolation, genome sequencing, and functional description of PHB degraders

PHB degrading isolates were obtained from red pine and sugar maple LnC soil by serial dilution. Diluted cells were plated onto mineral salts media containing 3 mM PHB (MSM-PHB) as the sole carbon source and incubated at room temperature as previously described [63]. Colonies appeared after 3 days and were isolated by streaking on MSM-PHB and stored in 20% glycerol (v/v) at -80°C. Genomic DNA was extracted from isolates as previously described and isolates were identified by Sanger sequencing of the 16S rRNA gene (27f/1492r) as previously described [64]. Isolates were screened for PHB degradation based on protocatechuate production with GC/MS (details in Supplementary Methods). An isolate, RP11, matched one of the predominant PHB-degrading phylotypes in amplicon libraries and its genome was sequenced using a quarter lane of Illumina MiSeq (2 × 250bp). The sequencing reads and genome assembly are available from NCBI BioProject: PRJNA558488. The strain was subsequently described as *Paraburkholderia madseniana* RP11^T^ (DSM 110123) and exhibited growth on several phenolics including *p*-coumaric and phthalic acids [63]. Details on the assembly, calculation of average nucleotide identity (ANI) and functional gene annotation are available in the Supplementary Methods.

### Quantifying pobA expression

Custom PCR primers were designed to target a 75-bp fragment of the ‘*pobA1*’ gene identified in the genome of isolate RP11^T^: pobA1_f (5’-AGA TCG AAT CCA CCA TCC GC-3’) and pobA1_r (5’-TTC AAA GCC GTG ATG CAA CG -3’) using NCBI’s Primer Blast software. Quantification of *pobA* and 16S rRNA transcripts (primers 341f/534r) was performed by RT-qPCR using an AB7300 Real-Time PCR system (Applied Biosystems) as described previously [65] (details in Supplementary Methods). Triplicate soil cores were taken with 3-mL plastic syringe barrels from each ecoplot and were immediately extruded into 10-mL serum bottles and dosed with 75 µL of ^13^C-labeled PHB (1000 ppm). The expression of *pobA* was determined from ∼1 g following a 24-hr incubation with PHB.

### Analysis of pobA in public databases

The environmental abundance of *pobA* was surveyed in publicly available metagenomic (*n* = 14 139) and genomic datasets (*n* = 11 643) through the IMG/ER portal [66]. The number of *pobA* copies per genome and the relative abundance in metagenomes were determined using the KEGG function search tool (March 19^th^, 2019). Taxonomic classifications were based on those provided by IMG/ER and the environmental source of metagenomes was attributed to the ‘habitat’ descriptor. Custom *pobA* Hidden Markov Models (HMMs) were generated from an alignment of homologs present in all *Burkholderiaceae* genomes in the NCBI refseq_genomic database (*n* = 3 991; as of June 20th, 2019). HMMs were developed for each of the seven *pobA* clades. A detailed description of all *pobA* analyses are provided in the Supplementary Methods and Supplementary Results, including information on enzyme structure and function and the phylogenetic trait depth of *pobA* paralogs [67].

### Data analysis and statistics

Amplicon libraries were quality processed using QIIME2 (v. 2017.9) [68] with dependency on DADA2 (v. 1.2.0) [69] to assign sequences to amplicon sequence variants (ASVs), referred to throughout as phylotypes. Taxonomic classification was performed using the QIIME2 ‘q2-feature-classifier’ trained on the Silva database (nr_v132) [70]. Libraries were filtered to remove ASVs present in reagent blanks or in low relative abundance (< 0.05% of all sequences) or few samples (minimum three samples). Estimates of species richness and Pielou’s evenness were calculated using rarified data (*n*_min_ = 9 140 per sample). All other analyses were performed on counts normalized to library totals (i.e. counts per thousand). All statistical analyses were performed in R (v. 3.4.0; R Core Team, 2018) with general dependency on the following packages: ggplot2 (v. 3.2.1) [71] and phyloseq (v. 1.22) [72]. Non-metric multidimensional scaling (NMDS) and perMANOVA were performed using the R package ‘vegan’ (v. 2.5-6) [73] on whole amplicon libraries using Unifrac distances based on a rooted tree produced using the QIIME2 wrapper ‘q2-phylogeny’ for MAFFT (v. 7.407) [74] and FastTree (v. 2.1.10) [75]. Variable selection for the major soil physicochemical parameters was performed using a stepwise regression with the ‘step’ function in R. These parameters were then fit to the NMDS with the ‘envfit’ function in vegan. 16S rRNA gene data from several studies of phenolic acid-induced priming, plant roots and lignin decomposition were re-analyzed using the aforementioned workflow to identify broader trends in bacterial populations of interest (see Supplementary Results). All analyses are reproducible using R scripts in the Supplementary Data.

## Results

Phenolic acid-induced priming of SOC was examined in soil microcosms using ^13^C-labeled substrates to differentiate the respiration of substrate from native SOC, while monitoring changes in microbial community composition and *pobA* expression. In addition, the composition and activity of PHB-degrading bacteria were determined in field soils using ^13^C-PHB DNA-SIP and by monitoring respiration and *pobA* expression. This combined approach enabled us to identify PHB-degrading taxa and link their priming activity to variation in SOC content of ecoplots, which differ in tree species and soil type (BL_LnC_, RP_LnC_, SM_LnC_, BL_MaB_, and SM_MaB_), in a 70-year old common garden experiment.

### PHB-induced soil priming in microcosms

The addition of PHB primed the decomposition of significant amounts of SOC during the first four days of incubation, when rates of PHB respiration were highest (Figure 1A). The majority of PHB had been mineralized by day 7 (Figure 1), corresponding with the cumulative priming of 3.3 (SM_LnC_), 5.5 (BL_Lnc_), and 13 (RP_LnC_) µmols C g^-1^ dry wt soil, representing 3.6%, 4.5%, and 8.7% more SOC mineralization than soils that received water only. In contrast, glucose reduced SOC mineralization between -3 to -8 µmols C g^-1^ dry wt soil in the same period. PHB degradation and priming were significantly higher in RP_LnC_ than BL_LnC_ and SM_LnC_ (TukeyHSD; p < 0.05), while differences between BL_LnC_ and SM_LnC_ were not significant. PHB was respired more rapidly and to a greater extent (93-100% of amendment) than glucose (62-71%) in all ecoplots (Figure 1B). Inhibition of *pobA*, while effective in pure culture (see Supplementary Results), was ineffective in soil, having no effect on PHB respiration or PHB-induced priming in soil microcosms (Figure 1).

**Figure 1.**
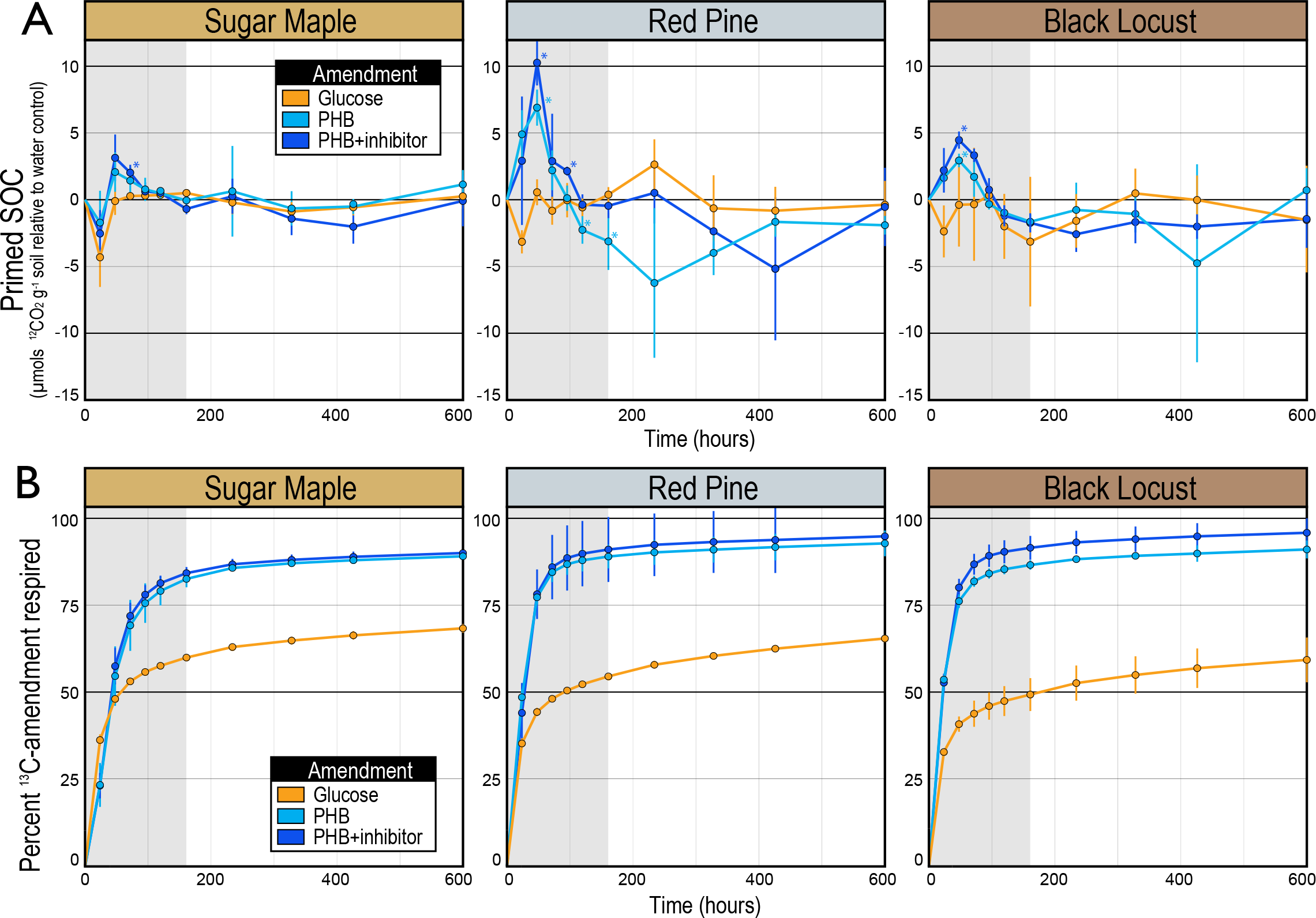
In soil microcosms, the enhanced mineralization of SOC is primed by the addition of PHB but not glucose. The addition of PHB caused positive priming (A), which corresponded with maximal respiration of ^13^C-PHB (B). Primed SOC is a measure of the native SOC respired relative to water-only controls. Error bars correspond to standard deviations among replicates (n = 4). Asterisks denote time points where respiration of native SOC significantly differed from water-only control. Priming was assessed in LnC soils which varied in soil organic matter content (BL_LnC_: 14.0% > RP_LnC_: 12.7% C > SM_LnC_: 10.2%).

In soil microcosms, 70 bacterial ASVs (‘phylotypes’) increased in relative abundance in response to glucose (n = 8), PHB (n = 10), water & glucose (n = 36) or glucose & PHB (n = 16; Table S2). *Paraburkholderia* and *Rhodanobacter* responded consistently to PHB and glucose, respectively (Table S2). Four of the seven phylotypes with the greatest response to PHB (relative abundance in PHB versus water-only) were *Paraburkholderia*, while the others were *Alicyclobacillus* (SM_LnC_), *Streptacidiphilus* (RP_LnC_) and an unclassified member of the phylum WPS-2 (RP_LnC_). The relative abundance of *Paraburkholderia* increased in response to PHB and glucose in all soils (Figure 2A), through this effect was not significant in BL_LnC_. In addition, *Paraburkholderia* responded more strongly to PHB than to glucose in SM_LnC_ soil (TukeyHDS; *p* < 0.01; Figure 2A). Transcripts of *pobA* were undetectable at day 7 (detection limit ∼ 300 copies per µL; data not shown) consistent with PHB mineralization dynamics (Figure 1A).

**Figure 2.**
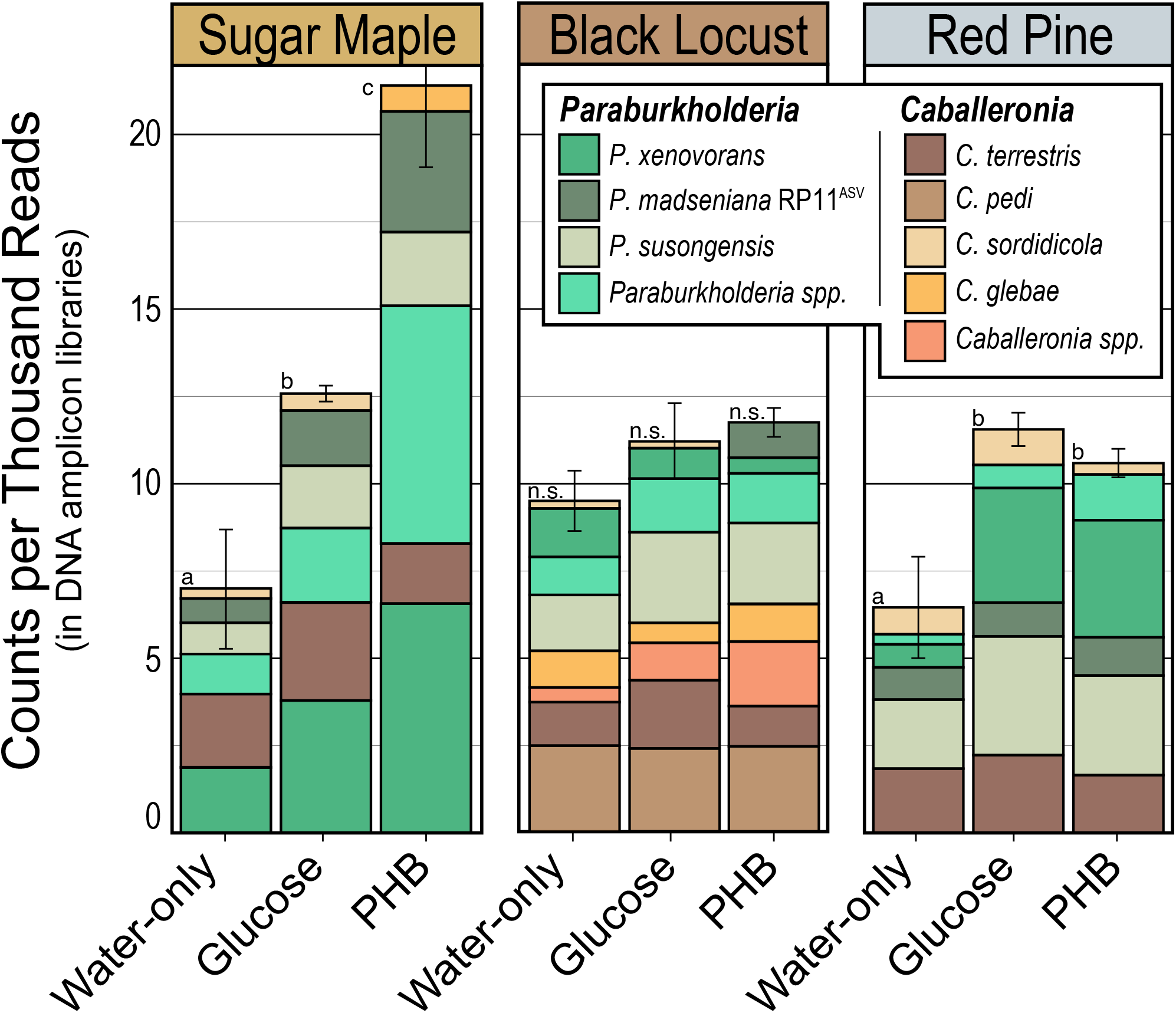
In soil microcosms, the relative abundance of *Paraburkholderia* and *Caballeronia* phylotypes increased when either PHB or glucose was added. Phylotypes were designated to a species group based on the top representative BLAST hit to type strains (100% identity).

Soils amended with glucose, or with RP11^T^ cells, mineralized less SOC than those that received only water (i.e., ‘negative priming’), though the effect was statistically significant only in the latter treatment (Figure S3). Amendment with RP11^T^ cells produced 5 - 14% less SOC mineralization when compared to controls. In all cases, negative priming was associated with steep increases in the relative abundance of *Streptomyces* and *Streptacidiphilus* phylotypes (Figure S3; details in Supplementary Results).

### PHB-degrading activity of field soils

The amount of PHB respired *in situ* differed considerably among ecoplots (Figure 3). PHB was respired rapidly in SM_LnC_ and RP_LnC_ soils with maximum instantaneous rates observed after about 10 hours (Figure 3AB). In contrast, PHB-degrading activity was relatively modest in SM_MaB_, BL_MaB_ and BL_LnC_, with substrate respiration plateauing at ∼ 1.5 – 2% of the total C added to soil (Figure 3B). SOC content was lowest in SM_LnC_ and RP_LnC_ and these plots had the highest rates of PHB respiration, *pobA* transcription, and the highest relative abundances of RP11^ASV^ (the phylotype of isolate RP11^T^; Figure 3CDE). These trends were not driven by total microbial biomass since RNA yield and 16S rRNA abundance were directly proportional to soil organic matter content (Figure S4AB).

**Figure 3.**
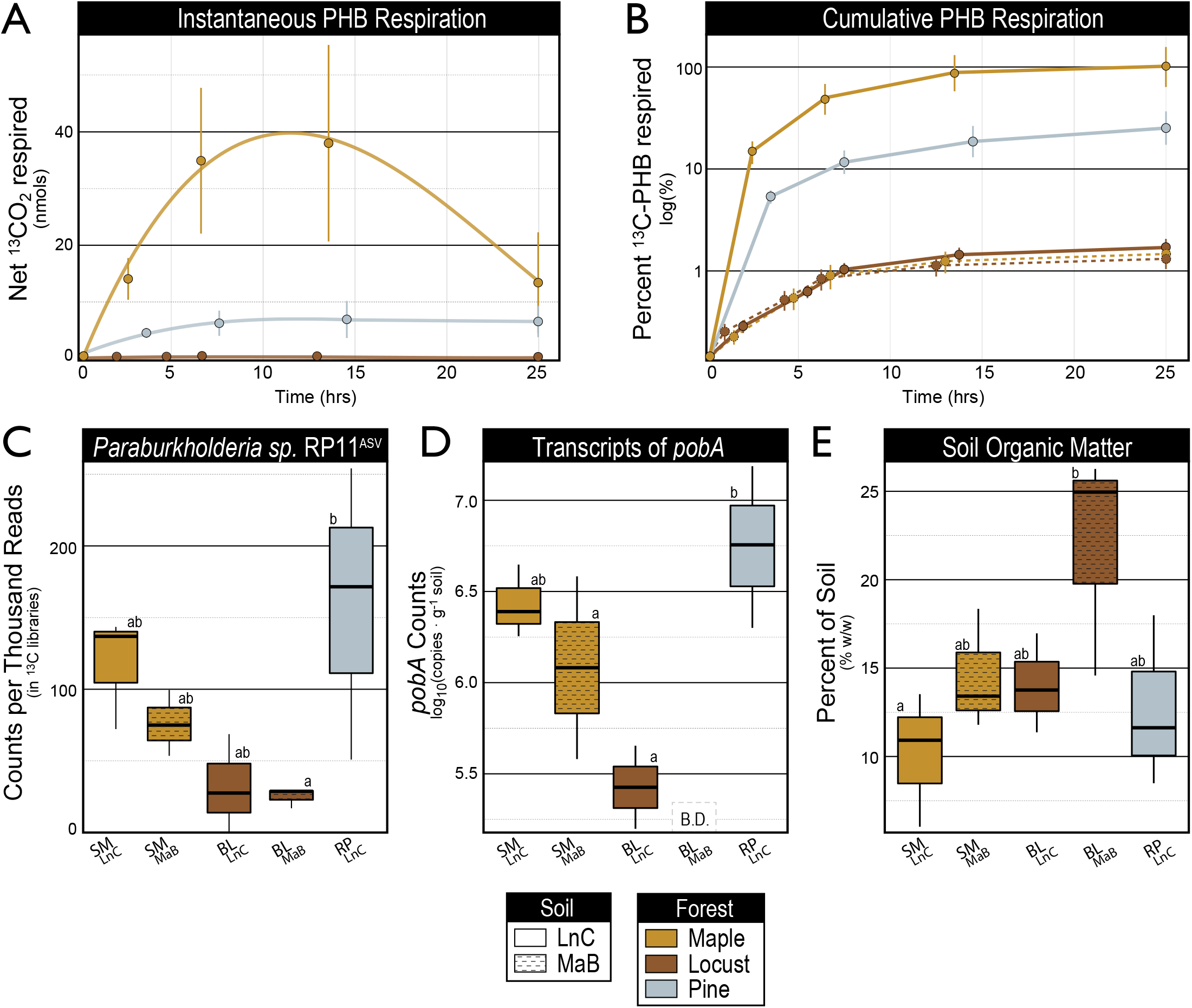
*In situ* measurements show that maximal rates of PHB respiration (A, B) are associated with high relative abundance of the RP11 phylotype (C), high levels of *pobA* transcripts (D), and low levels of soil organic matter (E). Error bars correspond to standard deviation (n = 3) and letters denote significant differences (p < 0.05) based on Tukey’s Honest Significant Difference tests (n = 3).

### PHB-degrading populations identified in field soils with DNA-SIP

Overall bacterial community composition in the field varied most by tree species (R^2^_perMANOVA_ = 25%; p < 0.001) followed by soil type (R^2^ = 14%; p < 0.001), illustrated by the NMDS ordination (Figure 4A). Of all the soil properties, pH and aluminum content explained the most variation in community structure (R2 = 14% and 9.6%, respectively; Table S3). BL forest soils had significantly lower pH, higher iron and sulfur, and lower cadmium and manganese than other forest types, while LnC and MaB soil types differed by aluminum and calcium content (Figure 4B; Table S4). These data show that major differences in soil properties and microbial communities had developed among ecoplots during the 70-year period after the common garden was planted.

**Figure 4.**
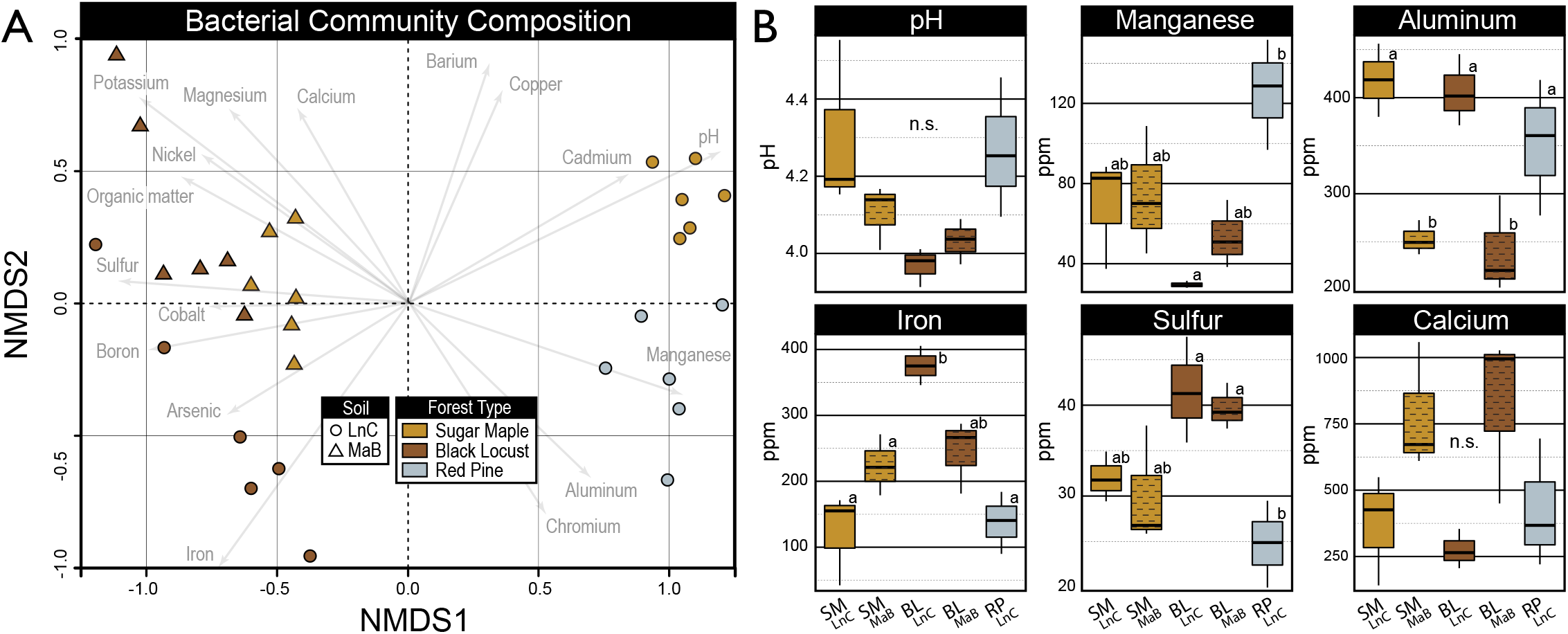
Variation in microbial community composition was driven by tree species composition and soil properties. NMDS ordination based on Bray-Curtis dissimilarity of 16S rRNA sequences from PHB-amended field soils indicate that soil series, tree species, and soil properties influence community composition (A). Soil physicochemical properties differed among ecoplots (B). In (A), community composition is represented by the average of all gradient fractions for ^13^C-labeled and ^12^C-control libraries for each ecoplot (n = 6). In (B), letters denote significant differences in soil properties (p < 0.05) based on Tukey’s Honest Significant Difference tests (n= 3). Measured concentrations correspond to the bioavailable elemental content of soils according to the modified Morgan extraction procedure. Comprehensive soil physicochemical properties are available in Table S4.

Despite these differences, we observed that the same six *Burkholderiaceae* phylotypes dominated *in situ* PHB degradation in all ecoplots (Figure 5A). To identify the bacteria responsible for PHB degradation *in situ*, we performed a DNA-SIP experiment with unlabeled PHB and ^13^C-labeled PHB applied directly to soil in all five ecoplots in the field. Bacterial communities profiled from the ^13^C-labeled DNA recovered from the heavy fractions of CsCl gradients exhibited greatly reduced richness and evenness relative to unlabeled DNA, indicating strong ^13^C-enrichment of PHB-degrading subpopulations (Figure S5). Between 10-40% of all amplicon sequences in ^13^C-DNA pools corresponded to phylotypes classified as *Paraburkholderia* (8 ASVs) and *Caballeronia* (4 ASVs), of which six predominated in all ecoplots (Figure 5AB). Most of these phylotypes were also enriched by PHB addition in the microcosm priming experiment, including four of the six phylotypes (Table S5). Two *Paraburkholderia* phylotypes, those matching *P. madseniana* (RP11^ASV^) and *P. xenovorans* LB400, were the dominant PHB-responders in both soil microcosm priming experiments and DNA-SIP experiments performed in field soils (Figure 2 and Figure 5).

**Figure 5.**
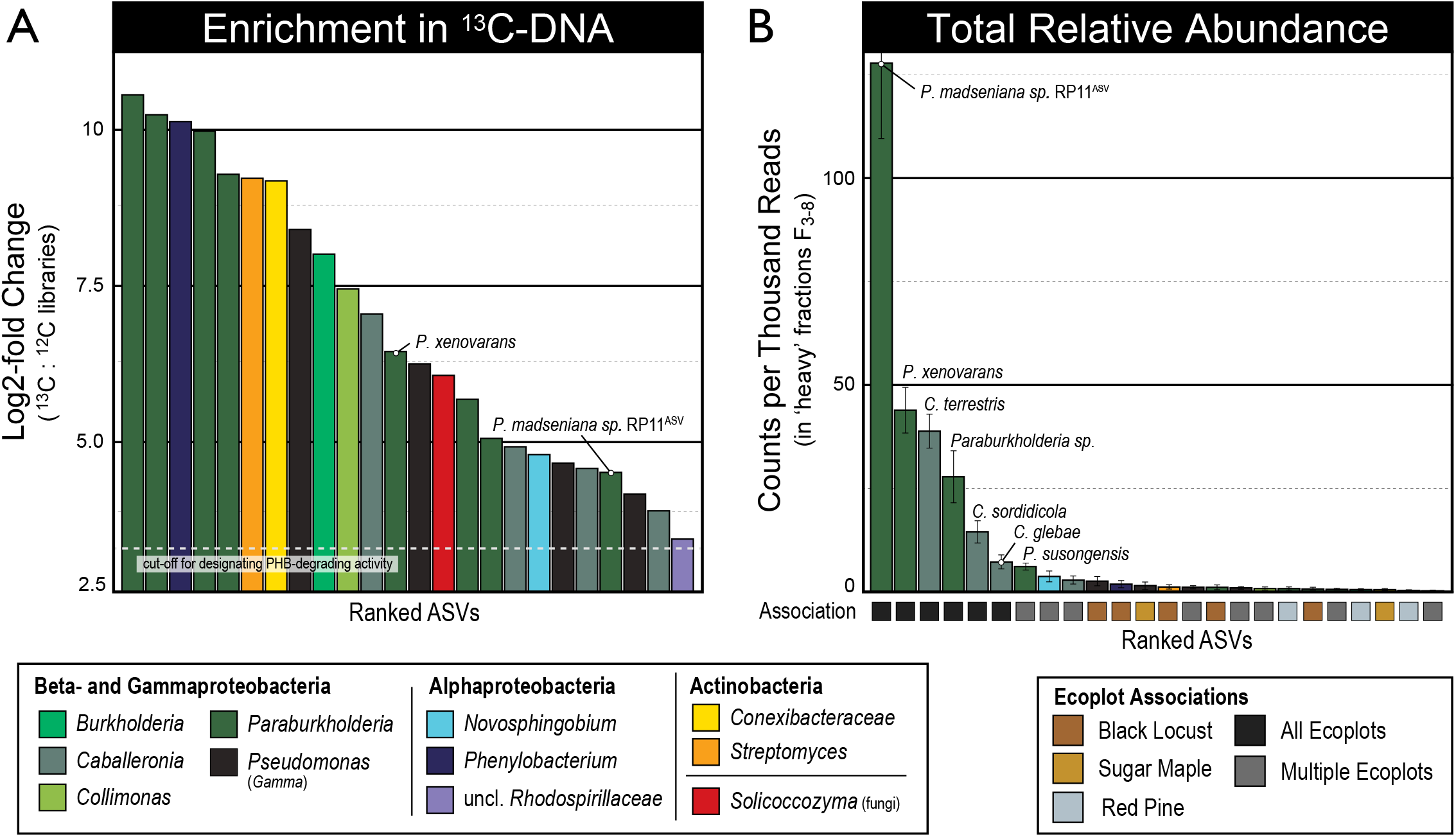
*Paraburkholderia* phylotypes (ASVs) are highly abundant in ^13^C-DNA recovered from DNA-SIP experiments in which ^13^C-PHB was added *in situ* to forest soils. *Paraburkholderia* represented one third of all phylotypes enriched significantly in ^13^C-DNA as determined by their log_2_-fold differential abundance relative to corresponding ^12^C-DNA controls shown in (A). *Paraburkholderia* and *Caballeronia* phylotypes identified in (A) comprised the seven most relatively abundant phylotypes in ^13^C-DNA, and these phylotypes were found in all ecoplots, as shown in (B). The two most abundant ^13^C-labeled phylotypes matched *P. madseniana* RP11^T^ and *P. xenovorans*, respectively (B). Bars are colored by taxonomic classification. Error bars correspond to standard deviation (n = 3). Phylotypes were designated to a species group based on the top representative BLAST hit to a type strain (100% identity). Details on the rank and relative abundance of ASVs are available in Table S5.

Less abundant ^13^C-PHB-labeled phylotypes detected in the DNA-SIP experiment included *Burkholderia, Collimonas, Pseudomonas* (*Gammaproteobacteria*), *Phenylobacterium* (*Alphaproteobacteria*), *Streptomyces* and *Conexibacteraceae* (*Actinobacteria*), and *Solicoccozyma* (a basidiomycotal yeast; Figure S6). These phylotypes were observed sporadically among forest and soil types (Figure 5B; Table S5), unlike phylotypes of *Paraburkholderia* and *Caballeronia* which predominated following PHB addition in all ecoplots.

### Isolation of PHB-degrading bacteria

Nine bacteria were isolated from RP_LnC_ soil using minimal media with PHB as the sole carbon source. All of these isolates produced protocatechuate (the reaction product of PobA) during growth on PHB (data not shown). Isolates were classified as *Paraburkholderia* (n = 5), *Pseudomonas* (n = 3) and *Cupriavidus*. Four of the *Paraburkholderia* isolates had identical full-length 16S rRNA genes that matched one of the dominant PHB-degrading phylotypes (RP11^ASV^) detected in DNA-SIP and microcosm experiments. One of these strains, isolate RP11^T^, was chosen for genome sequencing. The RP11^T^ genome was large (10.1 Mb), contained two chromosomes and encoded 6 copies of the *rrn* operon [63]. The genome encoded the complete pathway to mineralize PHB and included two paralogs of *pobA* which shared 48% amino acid sequence identity. RP11^T^ encoded an array of oxidative enzymes, including a DyP-type peroxidase, laccases, aryl-alcohol oxidases and an array of ring-cleaving and ring-hydroxylating dioxygenases (Table S6). Several of these enzymes contained signal peptide sequences, indicating potential for extracellular activity, including the DyP-type peroxidase, five laccases, and an aryl-alcohol oxidase [76].

### Phylogenetic diversity and environmental distribution of pobA-encoding bacteria

A search of publicly available genome sequences revealed that *pobA* homologs were most commonly encoded in genomes of *Betaproteobacteriales* and *Alphaproteobacteria*, occurring in more than 45% of genomes classified to these groups (Figure 6A, and Table S7). Genomes encoding paralogs (two copies of *pobA*) were more common in *Paraburkholderia* (25% of genomes) than any other genus of *pobA*-encoding *Burkholderiaceae* (Fisher’s exact, O.R. = 3.1, *p* < 0.001). In addition, *pobA* paralogs were more common in soil isolates compared to other environmental sources (O.R. = 2.7, *p* < 0.001; Figure S7A). Furthermore, *pobA* homologs were more abundant in forest soil metagenomes than in other environmental sources, and this difference was significant (Figure 6B). Homologs of *pobA* comprised seven major phylogenetic clades, and the two largest of these clades were represented by *pobA1* and *pobA2* of RP11^T^ (clades 6 and 5, respectively, Figure S7B). Additional information on the phylogenetic diversity and inferred activity of *pobA* homologs can be found in the Supplementary Results.

**Figure 6.**
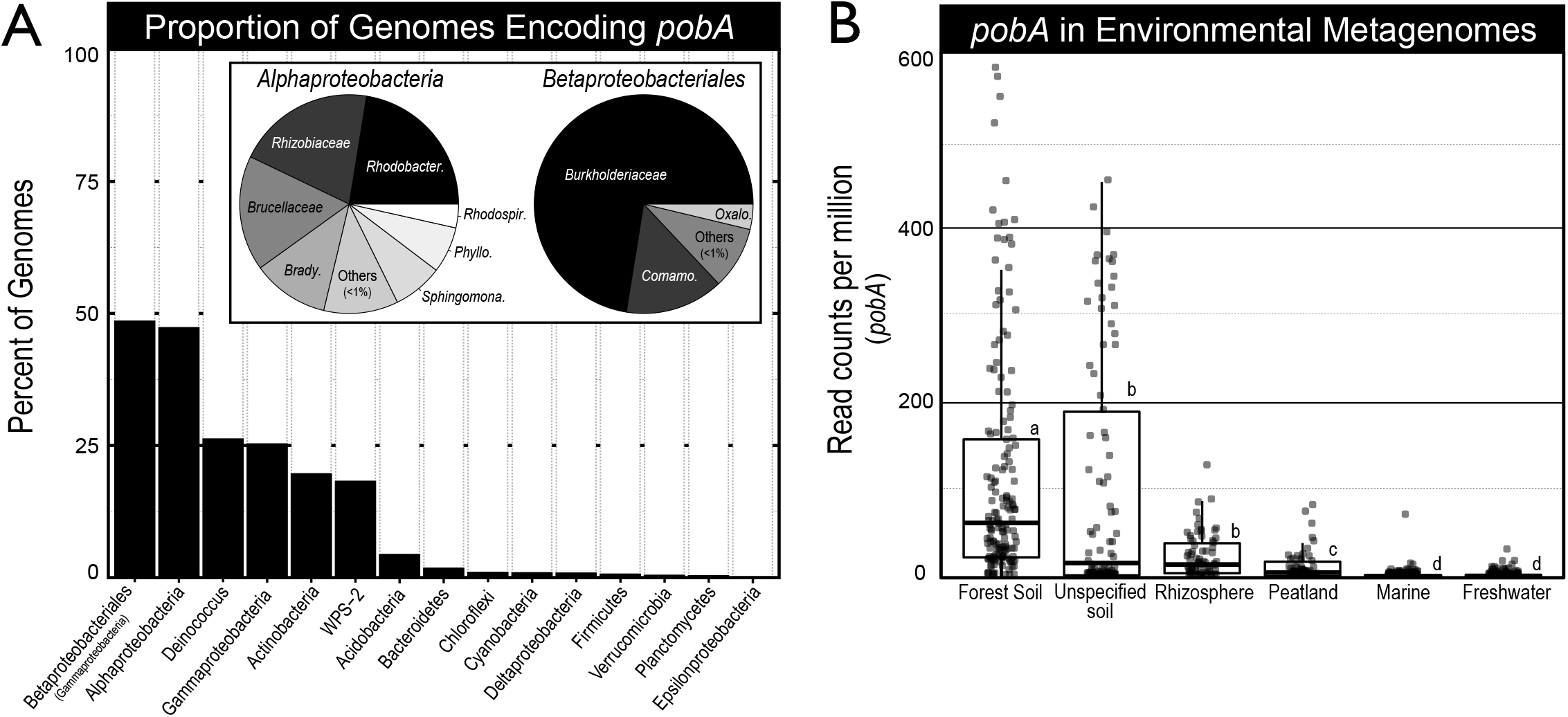
The phylogenomic distribution and environmental associations of *pobA* in (A) bacterial genomes (n = 11 643) and (B) environmental metagenomes (n = 14 139) from the IMG/ER database. In (A), phyla were ranked by the proportion of genomes encoding *pobA*. Only phyla represented by > 10 genomes were surveyed, and only those with at least one genome encoding *pobA* are shown. The taxonomic classification of *pobA*-encoding genomes from *Alpha*- and *Betaproteobacteria* are indicated by the inset pie charts in (A). Each taxonomic group was normalized to the total number of genomes present in the database. Homologs of *pobA* were far more common in soils in general, and in forest soils in particular, relative to other sample types as shown in (B). Letters denote significant differences (p < 0.05) based on pairwise Wilcoxon Rank Sum tests.

## Discussion

Our results demonstrate that the degradation of *p*-hydroxybenzoic acid (PHB) reliably primes the decomposition of SOC in forest soils. This phenomenon was driven by the activity of *Paraburkholderia* and, to a lesser extent, *Caballeronia*, from the family *Burkholderiaceae*. The predominant *Paraburkholderia* phylotypes were close relatives of *P. madseniana* RP11^T^ and *P. xenovorans* LB400^T^, species notable for having more genes and pathways associated with phenolic acid degradation than close relatives, and among the highest of any bacterial species [63, 77]. These findings are inconsistent with the hypothesis that all priming phenomena result from the non-specific, aggregate activity of functionally and phylogenetically diverse microbial populations [45]. While glucose addition stimulated growth of *Paraburkholderia*, it also stimulates a wide diversity of microorganisms whose competitive interactions likely govern the magnitude and direction of the priming response. In contrast, additions of PHB (Figure 1), benzoate [11] or vanillin [10, 12] cause consistent positive priming in forest soils and enrichment for *Paraburkholderia*. The identification of the specific taxa and functional genes responsible for SOC priming in forest soils is an important step toward understanding the mechanisms of soil priming. Furthermore, several lines of evidence suggest a more general role for phenolic acid-degrading bacteria in SOC cycling, such as the correspondence between SOC content in our common garden ecoplots with *in situ* rates of phenolic acid-degrading activity.

### Characteristics of PHB-induced soil priming

The catabolism of PHB and glucose differed in ways that revealed aspects of the nature of the priming effect. The addition of PHB to soil yielded significant positive priming with nearly twice the amount of PHB mineralized compared to glucose, which yielded slight, though insignificant, negative priming (Figure 1). This difference in use efficiency has been documented in forest soils, where added PHB was mineralized at twice the rate of glucose, and produced only half the microbial biomass [78]. The same effect occurred in grassland soil amended with vanillic acid, which yielded higher mineralization rates and lower microbial biomass than glucose or cellobiose, and also a greater positive priming effect [10]. We hypothesize that PHB-induced priming is caused, in part, by a metabolic imbalance whereby C catabolism greatly exceeds the needs of anabolism, evidenced by the high rates of PHB mineralization. Stoichiometric limitation-induced degradation of soil organic matter is a well-known mechanism in priming [45, 79–82], which could also explain why the turnover of RP11^T^ cells (a relatively rich nutrient source) was associated with strong negative priming (Figure S3). We propose that variation in the priming effect can be governed by substrate-specific metabolic use efficiency, creating differences in stoichiometric limitation. These dynamics are governed by life-history traits of decomposer populations or competitive and facilitative interactions with other soil microbes which are not yet fully understood.

Growth rate is a life-history trait that is correlated with substrate use efficiency and may contribute to the high mineralization rates of PHB. PHB-degrading populations were dominated by fast-growing bacteria which tend to be less efficient in converting substrate C into biomass [83]. Members of the family *Burkholderiaceae* are major zymogenous populations in soil [32] and the rapid *in situ* growth of RP11^ASV^ was evident in its increase from ∼ 0.7% to 15% of total sequences in 24 hrs, during the period of maximal PHB respiration. Consistent with these observations, isolate RP11^T^ exhibited rapid growth in pure culture with PHB as the sole carbon source (µ = 0.22 hr^-1^), and encodes six copies of the *rrn* operon, characteristic of high growth rate capacity [63, 84]. These results indicate that the microbial traits relevant to priming are partly defined by rapid growth, highlighting the importance of pulsed sources of phenolic C in SOC cycling.

Positive priming was also contingent on the metabolic activity of degrader populations. In RP_LnC_ and BL_LnC_, identical *Paraburkholderia* phylotypes responded to both glucose and PHB additions (Figure 2), yet only PHB induced positive priming (Figure 1). This phenomenon was apparent in a related study on soil priming, where the same *Paraburkholderia* phylotypes responded to benzoic acid and glucose amendment (also identical to our two major phylotypes), yet only benzoic acid elicited positive priming [11]. Notably, the co-addition of glucose and benzoic acid enhanced priming and coincided with an even greater increase in *Paraburkholderia*. We conclude that the metabolic activity induced by phenolics and aromatics is essential for the positive priming we observed, which may be enhanced by glucose when sufficient concentrations of aromatics are present to induce aromatic metabolism (Figure 7). Aromatic catabolism in *Paraburkholderia* is known to be regulated by concentrations of aromatic and phenolic compounds and insensitive to glucose [85, 86]. These results suggest that priming in forest soils is mediated by specific metabolic pathways and not by generic responses of the soil community to carbon addition.

The activity of oxidative enzymes has long been considered a potential mechanism in priming as a result of ‘co-metabolism’ [7, 80]. *Paraburkholderia* are renowned for their oxidative catabolism of aromatics, phenolics, and polyaromatic hydrocarbons [87, 88]. The two dominant PHB-degrading phylotypes in our study matched to *P. madseniana* and *P. xenovorans*, species which encode an extensive array of aromatic degrading pathways [52, 63]. *P. madseniana* RP11^T^ encoded among the greatest number of aromatic degrading genes compared to close relatives, including several putatively secreted oxidases [76] and aryl-alcohol oxidases important for bacterial lignin degradation [40]. *P. madseniana* RP11^T^ also grows on phthalic acid [63], a major by-product of lignin degradation [89–91]. Indeed, our analysis demonstrated that the two major PHB-degrading phylotypes of *Paraburkholderia* predominated in studies of lignin degradation [40], and white rot decay [41], and were the principal degraders in all soil priming experiments where phenolic/aromatic compounds were utilized [10–12] (see Supplementary Results). All evidence indicates that *Paraburkholderia*, and close relatives, monopolize phenolic acid degradation in soil and co-metabolize SOC, resulting in priming of SOC stocks.

**Figure 7.**
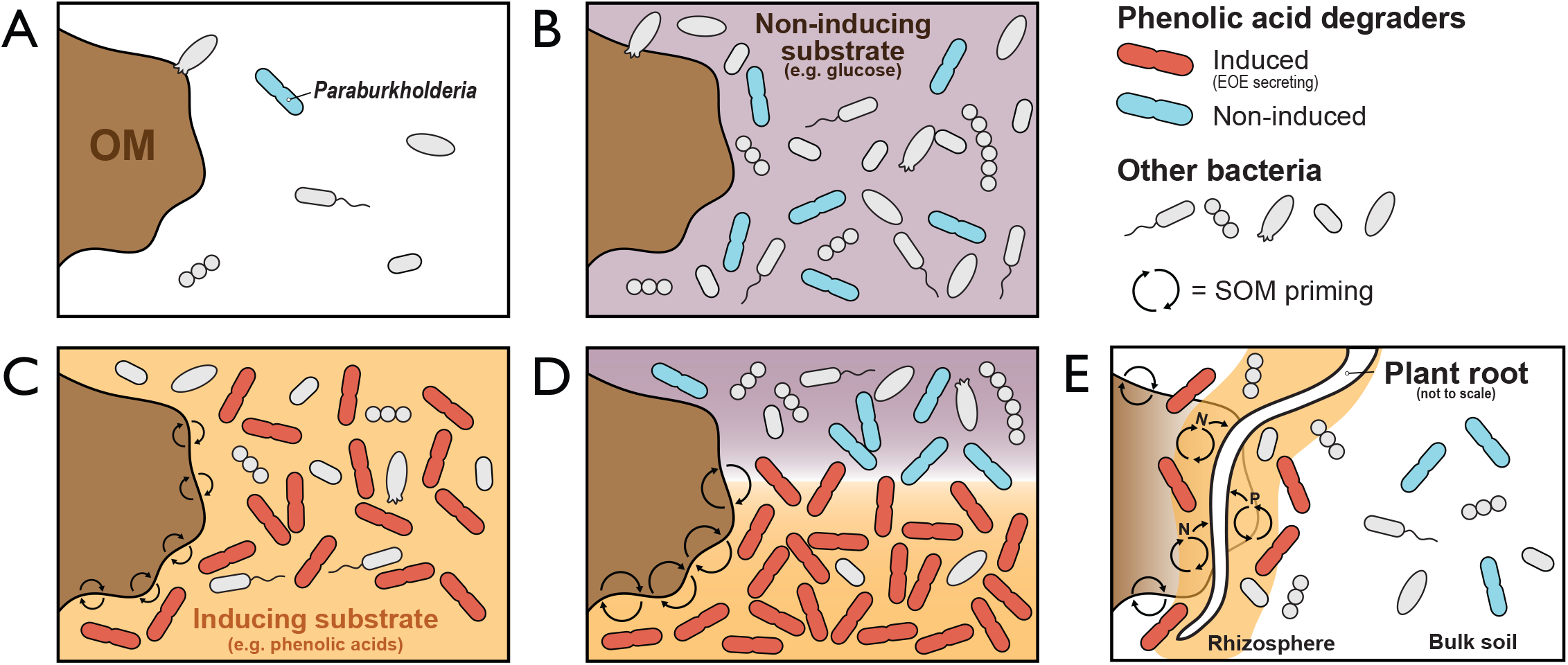
A conceptual framework illustrating the dependency of SOM priming on the abundance of phenolic acid-degrading bacteria and the induction of genes encoding extracellular oxidative enzymes (EOE) that degrade aromatic and polyaromatic compounds. In (A), limiting SOM priming occurs when phenolic acid-degrading bacteria are rare and aromatic metabolism is not induced. In (B), the addition of non-inducing substrates, such as glucose, stimulate growth of aromatic acid-degrading bacteria, but priming does not occur if the concentration of aromatic acids is insufficient to induce EOE. In (C), the addition of inducing substrates, such as phenolic or other aromatic acids, stimulate SOM priming by inducing EOE and by promoting the growth of specialized degrader populations. In (D), if the concentration of endogenous aromatic acids is sufficient to induce EOE then the co-addition of non-inducing substrates can enhance SOM priming by promoting the growth of phenolic acid-degrading bacteria. Hence, while glucose can stimulate the growth of *Paraburkholderia*, priming only occurs when aromatic metabolism is induced. In (E), plant roots gain access to inaccessible forms of N and P by stimulating the growth and induce priming activity of phenolic acid-degrading bacteria by exuding phenolic / aromatic acids.

### Impacts of phenolic acid-degrading populations on soil C-cycling

*Paraburkholderia* and *Caballeronia* routinely occupy sites where lignin- and phenolic-rich plant matter is decomposed [41, 42, 92–94], and are also common in root microbiomes [95], where phenolic-rich exudates recruit *Paraburkholderia* during root development [96]. The capacity for phenolic acids to prime decomposition, and the widespread association of these compounds with both living and dead plant biomass, raises questions about the broader impact of this phenomenon on soil C-cycling. Phenol oxidase activity and SOC content are inversely correlated at ecosystem scales, but this relationship is weak and highly interrelated with other soil properties [97]. In our common garden experiment, PHB-degrading activity and SOC content were inversely proportional, supporting the potential for phenolic acid-based priming to drive SOC dynamics. PHB-degrading activity was greatest where the relative abundance of *Paraburkholderia* and *pobA* expression were highest, and where SOC was lowest (SM_LnC_ and RP_LnC_). However, SOC content also followed trends in soil pH, iron, nickel and arsenic content which could independently affect SOC cycling. Earthworms can broadly affect SOC cycling in forest soils [98, 99], but, in our case, did not correspond with trends in SOC, as populations were historically highest in BL_LnC_, where SOC was greatest, and negligible in SM_LnC_ and RP_LnC_ [100].

We hypothesize that an association between plant roots and phenolic acid-induced SOC priming may explain difference in SOC accumulation among ecoplots. Both *Paraburkholderia* and *Caballeronia* are capable of nodulating legumes, including black locust [101], and exist endophytically in a broad range of plants [102–107]. All three of the closest relatives of *P. madseniana* RP11^T^ were isolated from plant roots: *P. sycomorum* ST111 (94.3% ANI; from *Acer* roots) [108], *P. aspalathi* LMG 27731^T^ (94.2%; *Aspalathus* root nodule) [109] and *P. sp*. OK233 (94%; *Populus* roots) [110]. The two dominant *Paraburkholderia* phylotypes we observed are common in maple and hemlock forest soils where shallow roots colonize upper, organic matter rich soil [11, 76, 111, 112]. At our field site, root density was by far greatest in sugar maple (SM_LnC_) than any other ecoplots according to field observations (Figure S1) and prior research [54]. The high *in situ* PHB-degrading activity in SM_LnC_ suggests a possible role for rhizosphere stimulation. The greater effect of ecoplot on PHB-degrading activity *in situ*, than in microcosms, supports the possibility that phenolic-rich root exudates may influence priming activity.

Phenolic root exudates facilitate plant-microbe interactions that can be essential for plant nutrition [43, 113]. The PHB content of root exudates double in poplar when these trees grow under conditions of nitrogen and phosphorus limitation [36]. *Paraburkholderia* are commonly associated with poplar roots [114, 115] where phenolic acids may stimulate their priming, nitrogen fixing [105, 111, 116] or mineral phosphorus-solubilizing activities [117, 118]. We hypothesize that certain tree species, and likely other plants, modulate the phenolic acid content of root exudates to recruit and stimulate phenolic acid-degrading bacteria to prime SOC degradation and access soil nutrients (Figure 7E). Differences in plant-microbe associations could, therefore, affect the dynamics of SOC cycling. In our field experiment, black locust plantations exhibited the lowest phenolic-degrading activity and greatest SOC content among ecoplots. Black locust trees can obtain nitrogen internally from symbiotic root nodules and would, therefore, have less need to promote SOC priming. Therefore, lower levels of phenolic acid degradation would be expected in soil colonized by black locust roots.

### Tracking priming activity with pobA

The expression of *pobA* (PHB monooxygenase) was predictably correlated with PHB-degrading activity and SOC priming. The use of *pobA* as a functional gene marker for priming activity has clear advantages over the 16S rRNA gene since we observed priming only during PHB metabolism, while the relative abundance of *Paraburkholderia* increased when both PHB and glucose were added. The fleeting nature of PHB-induced priming also attests to the importance of directly tracking functional activity, which may not necessarily produce major demographic shifts in community structure. The relative abundance of *pobA*, among other aromatic degradation genes, is also associated with vanillin-induced priming, corresponding with the predominance of the *P. xenovorans* phylotype [12]. Together, these findings illustrate the potential utility for tracking priming dynamics with PobA, a key enzyme in the peripheral degradation of phenolic acids [50].

## Conclusions

We conclude that specialized bacteria from the family *Burkholderiaceae* act as principal agents of phenolic acid-induced soil priming in forest soils, and likely more broadly. These populations consisted primarily of fast-growing *Paraburkholderia* characterized by their capacity to produce strongly oxidative catabolic enzymes, and their frequent associations with plant roots. These findings run counter to the hypothesis that soil priming is due to the broad stimulation of general microbial populations when soils are repeatedly dosed with glucose [45] or where a single bolus of glucose induces negligible or negative priming [11, 80, 119, 120].

The transient nature of PHB-induced priming underscores the importance of pulsed resource availability, or ‘hot moments’, in soil C-cycling [121]. Moving forward, it is imperative to characterize the role of phenolic acid-degrading activity and priming in natural settings where seasonal and root-mediated dynamics are considered. In one case, even slight warming over the winter (2.3 °C) led to a significantly greater priming of recalcitrant SOC by phenolic acid-degrading *Paraburkholderia* in Arctic tundra soil [12]. Future studies of phenolic acid-induced priming might help shed light on the nature of SOC persistence, which has previously been linked to low phenolic acid content [15, 16]. The findings from our 70-year old common garden experiment illustrate the potential role of phenolic acid-degrading *Paraburkholderia* in long-term C-cycling dynamics.

## Acknowledgements

We acknowledge the intellectual contributions of Drs. Marie Zwetsloot and Taryn Bauerle for their foundational research on root exudates and soil priming, as well as Dr. Yiu-Kwok Chan for enabling our access to a strain of N-fixing *P. xenovorans* 4B (ATCC 43038). We thank Drs. Murray McBride, Dominic Woolf and Timothy Fahey for their discussion of soil chemical, biogeochemistry and forest soil ecology, and Patricia Brenchley and Derrick Barret for their effective management of project resources. This work was supported by McIntire Stennis under NYC-189502 from the USDA National Institute of Food and Agriculture, and by the U.S. Department of Energy, Office of Biological & Environmental Research Genomic Science Program under award number DE-SC0016364. We would like to highlight the contributions of the late Dr. Eugene Madsen who initiated this research project prior to his untimely passing. May his contributions be an inspiration to future environmental microbiologists.

## Author Contributions

RW performed all data analysis, research and writing, as well as designing the priming experiment and assisting field work. CD performed all field experiments, methods development, including *pobA* primer design, and assisted with the priming experiment. JS identified the *pobA* paralogs and provided valuable feedback in preparation of the manuscript. EM conceived of the central aims of the study, selected the field site and designed and performed all field experiments. DB guided all research efforts and contributed significantly to designing the priming experiment, interpreting data and writing.

## Supplementary Tables

**Table S1**. A curated list of all isolates with characterized PHB degrading activity sourced from the eLignin database (http://www.elignindatabase.com/) [122].

**Table S2**. A complete list of all ASVs deemed to be responding to substrate amendments in the soil priming experiment according to indicator species analysis. The majority of trends were weak and inconsistent in all indicator groups. Therefore, manual validation was performed by assessing whether indicator assignments were driven by a single outlier sample and whether trends were consistent within a genus or family. Taxonomic groups present at high overall abundance were most likely to have individual ASVs, occurring in low abundance (< 0.5) identified as indicators that lacked robust support.

**Table S3**. perMANOVA tables showing the amount of variation in microbial community composition explained by major (A) treatment factors and (B) soil physicochemical parameters.

**Table S4**. Soil physicochemical properties according to forest and soil type. All measurements correspond to soil content (i.e. ppm of soil) calculated from bioavailable nutrients recovered using the modified Morgan extraction. Significant differences are denoted by lettering according to the Tukey HSD test, *p*-value <= 0.05.

**Table S5**. A complete list of all ASVs deemed to be ^13^C-enriched by PHB. The relative abundance and log2-fold enrichment of each ASV is given for each forest and soil type. The false-discovery rate corrected *p*-value calculated by DESeq is given. The majority of taxonomic classifications were based on SILVA. In several cases, SILVA returned ‘putative’ or ‘unclassified’ assignments at lower taxonomic ranks or unresolved genus groups, such as *Burkholderia*-*Paraburkholderia*-*Caballeronia*. In these cases, the genus corresponding to the top BLASTn hit was provided in place.

**Table S6**. A complete list of all dioxygenases and auxiliary activity enzymes (putatively lignin-degrading) encoded in the genome of *Paraburkholderia madseniana* RP11^T^. Cell have been colored (in an alternating pattern) according the genes present on the same contig.

**Table S7**. A complete list of all publicly available genomes encoding pobA from genera identified as ^13^C-labeled in our DNA-SIP dataset. These genomes were identified using IMG-ER on March 19th, 2019.

**Table S8**. (*cited in Supplementary Results*) Sample information for shotgun metagenomes obtained from a previously published SIP-lignin experiment [40].

**Table S9**. (*cited in Supplementary Results*) Summary of the phylogenetic conservatism and dispersion of several genes involved in catabolism of aromatics in the family *Burkholderiaceae*. In (A), estimates of phylogenetic conservatism were based on τ_D_ (trait depth) calculated with consenTRAIT. The lower the τ_D_ the less conserved the traits is among close relatives. In (B), phylogenetic dispersion was based on Purvis and Fritz’s D. A value of D < 0 suggests a highly clustered trait; D ∼ 0 reflects a Brownian motion (random walk) mode of evolution, D = 1 suggests a complete random mode of evolution and D > 1 suggests phylogenetic overdispersion (a lack of phylogenetic signal).

## Supplementary Figures

**Figure S1**. Photographs of field work measuring the respiration of ^13^C-labeled PHB (A-D) and images of soils from each forest type (E, F, and G). Soils dosed with PHB were eventually collected for DNA extraction and density gradient ultracentrifugation to recover ^13^C-enriched ‘heavy’ DNA corresponding to the PHB-degrading populations. In (A), a close-up of metal chambers used for collecting soil CO_2_ efflux deployed in the red pine ecoplot. In (B), the application of 150 µL of ^13^C-labeled or unlabeled PHB to soil in chambers embedded in the sugar maple ecoplot. In (C), sampling headspace for GC/MS analysis from triplicates in sugar maple plantation soil. In (D), a view of the landscape during field sampling in November of 2016. Macro images of field soils reveal differences in root density among forest types, where the dense root structures of sugar maple (E) where absent in red pine (F) and black locust (G). All root material was removed by sieving prior to microcosm soil priming experiment. Image scale differs slightly among E, F and G.

**Figure S2**. The design of the soil priming experiment displaying preparation methods, a numerically accurate representation of sample numbers, and a rationale for each treatment.

**Figure S3**. Trends in soil microcosm treatments (glucose and RP11^T^ cell amendments) where negative soil priming was observed (A and B) corresponded with the relative abundance of Streptomycetaceae phylotypes (C, D, and E). In (A), the spontaneous priming of native SOC over time and (B) the cumulative respiration of amended substrate is shown. Phylotypes were named according to the top representative BLAST hit to a type strain (100% identity). Taxonomic classifications correspond to a ‘species group’ since the V4 region of the 16S rRNA gene cannot resolve closely related species. In (C), stacked barplots display the relative abundance of all phylotypes in DNA extracts. In (D), the total relative abundance of *Streptomycetaceae* in DNA and cDNA pools are shown for soil microcosms amended with ^13^C-labeled RP11^T^ cells.

**Figure S4**. The size of active bacterial populations differed among ecoplots and was directly proportional to soil organic matter, as revealed in (A) RNA yields from DNAse treated soil extractions, (B) the abundance of copies of 16S rRNA according to RT-qPCR, and (C) the percent soil organic matter of field soil. In (A), RNA yields were measured from unamended and PHB-amended field soil in parallel for all ecoplots. In (B), RT-qPCR was performed on the RNA extracts measured in (A). Letters denote significant differences (*p* < 0.05) based on Tukey’s Honest Significant Difference tests (n = 3).

**Figure S5**. Species richness (A) and Pielou’s evenness (B) of ^13^C-enriched versus control 16S rRNA gene libraries in heavy gradient fractions (gradient fractions F_3_-F_8_) in all ecoplots. Asterisks denote significant differences (p < 0.05) based on Mann–Whitney tests (n=3 per). The average read depth per 16S rRNA gene library was 80,000 sequences, with a minimum of 2 200.

**Figure S6**. The ^13^C-enrichment of a *Solicoccozyma* phylotype evidenced by their increased relative abundance in heavy fractions of the CsCl gradients. Blue colored bars correspond to soil DNA extracts from soils incubated with ^13^C-labeled PHB, and pink bars to corresponding soils incubated with unlabeled PHB. The 18S rRNA gene was incidentally amplified using our universal bacterial primers (515f/805r). *Solicoccozyma* are common forest soil decomposers [123] and have been reported to degrade PHB [124, 125].

**Figure S7**. The gene for *p*-hydroxybenzoate 3-monooxygenase (*pobA;* EC 1.14.13.2) is encoded by numerous *Burkholderiaceae* genomes (n = 3 991), and duplicated *pobA* paralogs occur in many of these genomes (n = 410), as revealed by the maximum-likelihood phylogeny constructed from 24 single-copy genes shown in (A). *Burkholderia* were subdivided into sub-genera based on phylogenetic relatedness and presence (clades A, B and C) or absence (clade D) of *pobA* paralogs. Pie charts representing each clade indicate the isolation source of genomes that have one (indicated by a single black dot) or two (indicated by two black dots) copies of *pobA*, shown inset in (A). The genera *Caballeronia* and *Paraburkholderia* are paraphyletic suggesting that phylogenetic relationships within the *Burkholderiaceae* require further refinement, as previously reported [126]. The maximum likelihood phylogeny of *pobA* encoded by *Burkholderiaceae* is comprised of seven major clades, shown in (B). The leaves of the tree are colored with respect to genus, and the proportion of genera that comprise each *pobA* clade is inset. In clade 5, the category ‘others’ includes unclassified *Burkholderiaceae, Lautropia, Polynucleobacter, Robbsia*, and *Paucimonas* (in order of relative abundance) and, in clade 7 the category ‘others’ includes *Trinickia*. Phylogenetic trees in Newick format and HMM models for all *pobA* clades are available in the Supplementary Data package.

