## Supplementary material for "Phenolic acid-degrading *Paraburkholderia* prime decomposition in forest soil": 4a. pobA - structural analysis - tcoffee - espresso.html

T-COFFEE, Version\_11.00.d625267 (2016-01-11 15:25:41 - Revision d625267 - Build 507)  
Cedric Notredame   
CPU TIME:0 sec.  
SCORE=99  
\*  
 BAD AVG GOOD  
\*  
RP11\_pobA1   :  99  
RP11\_pobA2   :  99  
P\_fluor      :  99  
P\_CBS3       :  99  
cons         :   9  
  
RP11\_pobA1   MR----TQVGIIGAGPAGLLLSHLLHLRGIDSVVL-ESRTREQIESTIRAGVLEQGTMDLLTEAGVGAR  
RP11\_pobA2   MK----TKVVIIGAGPSGLLLGQLLQKAAIDNVIL-ERQSGEYVLGRIRAGVLEQGMVDLMREAGVSER  
P\_fluor      MK----TQVAIIGAGPSGLLLGQLLHKAGIDNVIL-ERQTPDYVLGRIRAGVLEQGMVDLLREAGVDRR  
P\_CBS3       MKSVTRTQVGIIGAGPAGLLLSHLLCIAGIDSVVVVESRSRAEMESTIRAGVLEQGTMDLLTSAGVGAR  
  
cons         \*:    \*:\* \*\*\*\*\*\*:\*\*\*\*.:\*\*   .\*\*.\*:: \* ::   : . \*\*\*\*\*\*\*\*\* :\*\*: .\*\*\*. \*  
  
  
RP11\_pobA1   MKAEGALHHGFELAFEGKRRRIDLTGLTG-RSITVYAQHEVIKDLVAARLAADGALR-FGVTETSLHGI  
RP11\_pobA2   MDAEGLVHDGIELVFGGRRDRIDLKKLTGGSSVLVYGQTEVTRDLMAAR-EASGATTIYEAANVQLHDV  
P\_fluor      MARDGLVHEGVEIAFAGQRRRIDLKRLSGGKTVTVYGQTEVTRDLMEAR-EACGATTVYQAAEVRLHDL  
P\_CBS3       MHAEGAVHHGIALAFEGERRRIDLTGLTG-RAITVYAQHEVIKDLVAAR-EAAGVLPVFEVTDTRIEDM  
  
cons         \*  :\* :\*.\*. :.\* \*.\* \*\*\*\*. \*:\*  :: \*\*.\* \*\* :\*\*: \*\*  \* \*.   : .::. :..:  
  
  
RP11\_pobA1   DTDQPSIRYRHEGEACELQCDFVIGCDGSQGVSRASIPQALR-KDFERVYPFGWFGILCEGPPSSEELI  
RP11\_pobA2   KGEAPYVTFEKSGETYRLDCDYVAGCDGFHGVSRKTIP-AEVLTHYERIYPFGWLGLLSDTPPVNHELI  
P\_fluor      QGERPYVTFERDGERLRLDCDYIAGCDGFHGISRQSIP-AERLKVFERVYPFGWLGLLADTPPVSHELI  
P\_CBS3       DTEKPVVRYVRDGVDGALVCDYVVGCDGFHGPSRQTIP-VQAREEFERVYPFGWFGILVEAPPSSEELM  
  
cons         . : \* : : :.\*    \* \*\*:: \*\*\*\* :\* \*\* :\*\* .     :\*\*:\*\*\*\*\*:\*:\* : \*\* ..\*\*:  
  
  
RP11\_pobA1   YARHDRGFALVSTRSANVQRMYFQCDPKDSVDNWSDDRIWAELHARVDSHDGQHIVDGKIFQKNIVGMR  
RP11\_pobA2   YAHHERGFVLCSQRSTTRSRYYLQVPLTEKAEDWSDERFWEELKTRLPQDVAEKVVTGPSLEKSIAPLR  
P\_fluor      YANHPRGFALCSQRSATRSRYYVQVPLTEKVEDWSDERFWTELKARLPAEVAEKLVTGPSLEKSIAPLR  
P\_CBS3       YARHDRGFALVSTRSPGIQRMYFQCGPSESVESGRTRKIWEELHTRLDSIDGWKIIKGKIFQKNIVGMR  
  
cons         \*\*.\* \*\*\*.\* \* \*\*.  .\* \*.\*   .:..:.    ::\* \*\*::\*:    . ::: \*  ::\*.\*. :\*  
  
  
RP11\_pobA1   SFVSATMQHGRLFLAGDAAHIVPPTGAKGMNLAVADVRVLTQALSAFYVE----YRTD-LLDSYSATAL  
RP11\_pobA2   SYVVEPMQYGKLFLVGDAAHIVPPTGAKGLNLAASDVSTLYRILNRVYRD----GRTD-LLEKYSQIAL  
P\_fluor      SFVVEPMQHGRLFLAGDAAHIVPPTGAKGLNLAASDVSTLYRLLLKAYRE----GRGE-LLERYSAICL  
P\_CBS3       SFVCATMRYGRLLLAGDAAHIVPPTGAKGLNLA---VNNL-RLLAKAFAELYTTGSQERLLS-YSHNAL  
  
cons         \*:\*  .\*::\*:\*:\*.\*\*\*\*\*\*\*\*\*\*\*\*\*\*:\*\*\*   \*  \* : \*   : :       : \*\*. \*\*  .\*  
  
  
RP11\_pobA1   KRIWRAEHFSYWMTSMMHRIEGASPFEQQLQVAELEYVTTSRAA-ATVMAENYVGIAVV---  
RP11\_pobA2   RRVWKAERFSWFMTNLLHEFSEHDTFDKRMQRTDYDYYTISEAGLKTI-AENYVGLPYESIE  
P\_fluor      RRIWKAERFSWWMTSVLHRFPDTDAFSQRIQQTELEYYLGSEAGLATI-AENYVGLPYEEIE  
P\_CBS3       RRVWRPEHFSWWITTMLHNFDNATPFHQRLLVAQLDYLTTSEAGARVL-SENYLG-PLTH--  
  
cons         :\*:\*:.\*:\*\*:::\*.::\*.:    .\* :::  :: :\*   \*.\*.  .: :\*\*\*:\* .       
  
  
  
  
  
