## Supplementary material for "Phenolic acid-degrading *Paraburkholderia* prime decomposition in forest soil": BLOCK01a - in situ - Respiration and qPCR.R

```
library(plyr)
library(ggplot2)

### Calculate Standard Error
summarySE <- function(data=NULL, measurevar, groupvars=NULL, na.rm=FALSE,
                      conf.interval=.95, .drop=TRUE) {
  library(plyr)
  
  # New version of length which can handle NA's: if na.rm==T, don't count them
  length2 <- function (x, na.rm=FALSE) {
    if (na.rm) sum(!is.na(x))
    else       length(x)
  }
  
  # This does the summary. For each group's data frame, return a vector with
  # N, mean, and sd
  datac <- ddply(data, groupvars, .drop=.drop,
                 .fun = function(xx, col) {
                   c(N    = length2(xx[[col]], na.rm=na.rm),
                     mean = mean   (xx[[col]], na.rm=na.rm),
                     sd   = sd     (xx[[col]], na.rm=na.rm)
                   )
                 },
                 measurevar
  )
  
  # Rename the "mean" column    
  datac <- rename(datac, c("mean" = measurevar))
  
  datac$se <- datac$sd / sqrt(datac$N)  # Calculate standard error of the mean
  
  # Confidence interval multiplier for standard error
  # Calculate t-statistic for confidence interval: 
  # e.g., if conf.interval is .95, use .975 (above/below), and use df=N-1
  ciMult <- qt(conf.interval/2 + .5, datac$N-1)
  datac$ci <- datac$se * ciMult
  
  return(datac)
}

##########################
## Figure 3D and Figure S6
##########################

# import data
x <- read.csv(file = "raw_data/in.situ.qPCR.cDNA.data.csv", header=T, stringsAsFactors = F)

# make combos
x$tree <- gsub(" ","_",x$tree)
x$combo <- paste(x$tree, x$soil.type, sep="_")
x$combo <- factor(x$combo, levels = c("Sugar_Maple_LnC","Sugar_Maple_MaB", "Black_Locust_LnC", "Black_Locust_MaB","Red_Pine_LnC"))
x$copies.per.g <- x$copies/0.5

x <- subset(x, !(treatment %in% c("H2O.amended","autoclaved")))

# Figure S6B - qPCR Counts of 16S rRNA Gene
plot <- ggplot(subset(x, measure == "rRNA.16S"), aes(x=combo, y=log(copies.per.g, 10), fill = soil.type)) + geom_boxplot(outlier.shape = NA) + ylab("copies 16S rRNA gene per g soil")
plot

# stats
foobie <- data.frame(values = log(subset(x, measure == "rRNA.16S")$copies.per.g, 10), combo = subset(x, measure == "rRNA.16S")$combo, stringsAsFactors = F)
fit <- lm(values ~ combo, data = foobie)
TukeyHSD(aov(fit))

# pobA
plot <- ggplot(subset(x, measure == "pobA.2"), aes(x=combo, y=log(copies.per.g, 10), fill = soil.type)) + geom_boxplot(outlier.shape = NA) + ylab("copies pobA.2 gene per g soil")
plot

# stats
fit <- lm(copies.per.g ~ combo, data = subset(x, measure == "pobA.2"))
fit <- TukeyHSD(aov(fit))


###################
## Figure 3A and 3B
###################

# import
x <- read.csv(file = "raw_data/in.situ.co2.respiration.data.csv", header = T, stringsAsFactors = F)
x$h.chamber.cm <- as.numeric(x$h.chamber.cm)

# remove unused columns
x$experiment <- NULL

# convert peak height to concentration 

## Calculate mols C
# calculate headspace volume (note: 1mL == 1 cm3)
x$volume <- (3.141593*((1.25)^2)*x$h.chamber.cm)/1000  # V=πr2h (radius and height in cm) & convert to L (1000 mL / 1L)
x$DeRito.volume.uL <- x$volume*1000  # from DeRio spreadsheet
#dilution.factor <- standard_injection_volume/100  # we inject 100 uL of sample

# Get standard curve  (from DeRito file '161019 Turkey Hill CO2.xls')
standards <- data.frame(concentration.nM = c(3.73,	3.73,	7.46,	7.46,	11.2, 11.2, 18.6, 37.3, 55.9, 74.6), mz.44 = c(61430, 62369, 118025, 121185, 179882, 182744, 361810, 752300, 1094153, 1425203), mz.45 = c(94, 221, 901, 1133, 1709, 1838, 3898, 8478, 12736, 16558))

# adjust mz.45 on low end of concentration
standards$mz.45[1:6] <- standards$mz.44[1:6]*0.011

# Visualize
ggplot(standards, aes(concentration.nM, mz.45)) + geom_point() + geom_smooth(method=lm, se=F)

m <- coef(lm(concentration.nM ~ mz.45, data = standards))[2]
b <- coef(lm(concentration.nM ~ mz.45, data = standards))[1]

# Calculate nM from standard curve
#x$newnM.C <- ((x$mz.45*m + b)*dilution.factor # Only if different volume of standard was used
x$nM.C <- x$mz.45*m + b

# Calculate nmols from standard curve
x$nmols.C <- x$nM.C * x$volume

# calculate nM C13 respired
x$DeRito.nM.C = ((x$mz.45/19282)/100)*x$DeRito.volume.uL  # Not clear how equation was derived or where '19282' comes from

# adjusted nM based on average standard value
x$DeRito.nM.C.adj <- x$DeRito.nM.C*(2120652/mean(subset(x, tree == "std")$mz.44)) # Not clear how equation was derived or where '2120652' cines from

# remove standards
x <- subset(x, tree != "std")

# make comboID  
x$tree <- gsub(" ","_",x$tree)
x$combo <- paste(x$tree, x$soil.type, sep="_")

# Calculate Cumulative Respiration
x$comboID <- paste(x$combo, x$replicate, x$enrichment, sep = "_")

count = 1
for (sample in unique(x$comboID)){
  foo<-subset(x, comboID == sample)
  foo$time <- as.numeric(foo$time)
  foo<-foo[order(foo$time),]
  
  # Need to impute values for C12 samples missing timepoints
  # make assumption that respiration is ~ 70% of the increase from between the two timepoints
  if (any(is.na(foo$nmols.C))){
    for (n in which(is.na(foo$nmols.C))){
      value = (foo$nmols.C[n+1]-foo$nmols.C[n-1])*0.7
      value = value+foo$nmols.C[n-1]
      foo$nmols.C[n] <- value
    }
  }
  
  foo$cum.nM.C <- cumsum(foo$nmols.C)
  
  if (count == 1){
    cumulative <- foo
    count = count + 1
  } else{
    cumulative <- rbind(cumulative, foo)
  }
}

#########
# Plot 1 : Instantaneous Respiration

plot_me <- summarySE(x, measurevar="nmols.C", groupvars=c("time","combo","enrichment"))
plot_me$time <- as.numeric(plot_me$time)
levels(plot_me$enrichment) <- c("C12", "C13")
plot <- ggplot(plot_me, aes(time, nmols.C, color = combo)) + geom_point() + geom_smooth(method = "lm", formula = y ~ splines::bs(x, 3), se = FALSE) + geom_errorbar(aes(ymin=nmols.C-se, ymax=nmols.C+se), width=.1) + facet_grid(~enrichment)
plot


#########
# Plot 2 : Cumulative C13 Respiration

# subtract natural abundance C13
cumulative.C13 <- reshape(cumulative[,c("time", "combo", "enrichment", "replicate", "cum.nM.C")], direction = "wide", idvar = c("combo","replicate","time"), timevar = "enrichment")
cumulative.C13$cum.nmols.C <- cumulative.C13$cum.nM.C.C13 - (cumulative.C13$cum.nM.C.C12*0.011)
cumulative.C13$cum.nmols.C[which(cumulative.C13$cum.nmols.C < 0)] <- 0
cumulative.C13$cum.nmols.C[which(cumulative.C13$time == 0)] <- 0

# Plot
plot_me <- summarySE(cumulative.C13, measurevar="cum.nmols.C", groupvars=c("time","combo"))
plot_me$time <- as.numeric(plot_me$time)
plot <- ggplot(plot_me, aes(time, log(cum.nmols.C, 10), color = combo)) + geom_point() + geom_line() + geom_errorbar(aes(ymin=log(cum.nmols.C-se, 10), ymax=log(cum.nmols.C+se,10)), width=.1)
plot


#########
# Plot 3B : Cumulative % Total Substrate Respiration

# We added 150 µl of 1000 ppm PHB to soil
# molecular weight of PHB = 137.114 g / mol
# 137,114 ppm = 1 M
M.PHB <- 1000/137114          # mols/L
mols.PHB <- M.PHB*0.00015     # mols
nmols.PHB <- mols.PHB/10e-09  # nmols

# convert nmolsC to percent of total
cumulative.C13$percent.PHB <- (cumulative.C13$cum.nmols.C/nmols.PHB)*100

ddply(cumulative.C13, ~ combo, summarise, most = max(percent.PHB))

# Plot
plot_me <- summarySE(cumulative.C13, measurevar="percent.PHB", groupvars=c("time","combo"))
plot_me$time <- as.numeric(plot_me$time)
plot <- ggplot(plot_me, aes(time, percent.PHB, color = combo)) + geom_point() + geom_line() + geom_errorbar(aes(ymin=percent.PHB-se, ymax=percent.PHB+se), width=.1)
plot


#############
## Figure S6A

## RNA Quantification
x <- read.csv(file = "raw_data/in.situ.nucleic.acid.quant.csv", header = T, stringsAsFactors = F)
x$tree <- gsub(" ","_",x$tree)
x$combo <- paste(x$tree, x$soil.type, sep="_")
x$concentration <- as.numeric(x$concentration)
x$treatment <- factor(x$treatment, levels = c("unamended.soil", "PHB.amended"))
x$combo <- factor(x$combo, levels = c("Sugar_Maple_LnC","Sugar_Maple_MaB", "Black_Locust_LnC", "Black_Locust_MaB","Red_Pine_LnC"))

plot <- ggplot(x, aes(x=combo, y=concentration, fill = treatment)) + geom_boxplot(outlier.shape = NA) + ylab("[RNA] ng/uL") 
plot

# stats
for (data in unique(x$treatment)){
  print(paste("-------- Dataset: ",data," --------"))
  fit <- lm(concentration ~ combo, data = subset(x, treatment == data))
  print(TukeyHSD(aov(fit)))
}
```
