## Supplementary material for "Phenolic acid-degrading *Paraburkholderia* prime decomposition in forest soil": BLOCK01b - in situ - Importing and Normalizing Data.R

```
library(phyloseq)

#################################
## Fix Sample Data in SSU Dataset
p <- readRDS(file = "raw_data/p_merged.final.rds")  # This was created by QIIME2 by merging the table.qza and rep.seqs.qza from the three separate runs.
                                                    # A tree was created using the merged rep.seqs in QIIME2, too.

# Fix Sample Information  (mixed-up during QIIME2 processing because there were no column names provided in metadata file)
x.meta <- read.table(file = "raw_data/sequencing.metadata.tsv", stringsAsFactors = F, header = F)
colnames(x.meta) <- c("SampleID","BarcodeSequence","LinkerPrimerSequence","tree","soil.type","enrichment","fraction.type","fraction","latitude","longitude","env_biome","derito.label","run.name","soil.collection","run.data","Original.barcode")
x.meta <- subset(x.meta, SampleID != "M047")  #Failed sample
x.meta$rep <- gsub("SugarMaple|BlackLocust|RedPine|LnC|MaB|\\.|12|13|Fraction1.2|Fraction3.4|Fraction5|Fraction6|Fraction7|Fraction8|Fraction9.10|Fraction11|Rep","", x.meta$derito.label)
x.meta$ComboID <- paste(x.meta$soil.type, x.meta$tree, x.meta$rep, sep="_")

# Remove Huson Samples (separate project with Dr. Ian Huson)
p <- prune_samples(x.meta[!is.na(x.meta$rep),"SampleID"], p)
x.meta <- x.meta[!is.na(x.meta$rep),]

# Add in Elemental Analysis Data
x.EA <- read.csv(file = "raw_data/soil.chem.data.csv", stringsAsFactors = F, header = T)
x.EA$tree <- gsub(" ","_",x.EA$tree)
x.EA$depth <- gsub(" ","",x.EA$depth)
x.EA <- subset(x.EA, depth == "0-2.5")
x.EA$ComboID <- paste(x.EA$soil.type, x.EA$tree, x.EA$rep, sep="_")

# Merge Data
x.meta <- merge(x.meta, x.EA, by = "ComboID")
rownames(x.meta) <- x.meta$SampleID
x.meta$SampleID <- NULL
x.meta$ComboID <- NULL
x.meta$soil.type.y <- NULL
x.meta$tree.y  <- NULL
x.meta$rep.y  <- NULL
x.meta$sampleID  <- NULL
colnames(x.meta) <- gsub(".x","",colnames(x.meta))

# Incorporate New Correct and Complete Sample Data into phyloseq
p_foo <- merge_phyloseq(otu_table(p), tax_table(p))
p_foo <- merge_phyloseq(p_foo, phy_tree(p))
x.meta <- x.meta[match(sample_names(p_foo), rownames(x.meta)),]
x.meta <- sample_data(x.meta)
p <- merge_phyloseq(p_foo, x.meta)

######################################################
## Pre-process to Filter Low Abundance and Sparse Taxa

# Keep information about the total # of sequences and OTUs
total_seqs <- sum(sample_sums(p))
total_otus <- length(taxa_names(p))

## Subtract OTUs Present in Blanks
p_blanks <- subset_samples(p, rep == "Blank")
p_blanks <- subset_taxa(p_blanks, taxa_sums(p_blanks) > 0)
p_filtered <- prune_taxa(setdiff(taxa_names(p), taxa_names(p_blanks)), p)
p_filtered <- subset_samples(p_filtered, rep != "Blank") 

print(paste("The Percent of Sequences Remaining After Removing Contaminants Found in Blanks is",round((sum(sample_sums(p_filtered))/total_seqs)*100, 2),sep=" "))
print(paste("The Number of OTUs was Reduced from", total_otus,"to", length(taxa_names(p_filtered)), "(i.e.", round((length(taxa_names(p_filtered))/total_otus)*100,1),"% of OTUs remain)", sep=" "))

# Save 'fixed' phyloseq object
saveRDS(p_filtered, file = "raw_data/p_merged.final.filtered.rds")

###############################
## Filter Low Abundance Samples

# Add in Temporary Column
sample_data(p)$Combo <- paste(sample_data(p)$tree ,sample_data(p)$soil.type, sep ="_")

# Remove Libries with Low Counts
p  <- prune_samples(names(sample_sums(p)[which(sample_sums(p) > 2000)]), p)  

print(paste("The Percent of Sequences Remaining After Removing Low Abundance Libraries is",round((sum(sample_sums(p))/total_seqs)*100, 2),sep=" "))
print(paste("The Number of OTUs was Reduced from", total_otus,"to", length(taxa_names(p)), "(i.e.", round((length(taxa_names(p))/total_otus)*100,1),"% of OTUs remain)", sep=" "))


#######################################
## Filter Low Abundance and Sparse OTUs

# Set Filters
filter1 <- mean(sample_sums(p)) * 0.0005  #
filter2 <- mean(sample_sums(p)) * 0.01

# Hard Filter OTUs below 0.05% of each sample's Total Abundance (i.e. summed across all samples)
p <- prune_taxa(names(which(taxa_sums(p) > filter1)), p)

# Soft Filter OTUs which occur in less than three samples (with two exceptions)
present_absent<-otu_table(p) 
present_absent[present_absent > 0] <- 1

# 1) Remove all singletons except those with a max relative abundance greater than 1% in any sample
REMOVE_singletons <- rownames(present_absent)[which(rowSums(present_absent) == 1)]
p_assess <- prune_taxa(REMOVE_singletons, p)
KEEP_singletons <- names(which(apply(otu_table(p_assess), 1, function(x) max(x) > filter2)))
REMOVE_singletons <- setdiff(REMOVE_singletons, KEEP_singletons)

# 2) Remove all doubletons except those with a max relative abundance greater than 1% in any sample OR occur in the same forest and soil type
REMOVE_doubletons <- rownames(present_absent)[which(rowSums(present_absent) == 2)]
p_assess <- prune_taxa(REMOVE_doubletons, p)

KEEP_doubletons <- names(which(apply(otu_table(p_assess), 1, function(x) max(x) > filter2)))
CHECK_doubletons <- prune_taxa(names(which(apply(otu_table(p_assess), 1, function(x) max(x) < filter2))), p_assess)

KEEP_paired_doubles <- vector()
for (taxa in taxa_names(CHECK_doubletons)){
  foo <- prune_taxa(taxa, CHECK_doubletons)
  foo <- prune_samples(names(which(sample_sums(foo) > 0)), p)
  
  if (nrow(unique(sample_data(foo)[,"Combo"])) == 1) {
    KEEP_paired_doubles <- c(KEEP_paired_doubles, taxa)
  }
}

KEEP_doubletons <- c(KEEP_doubletons, KEEP_paired_doubles)

# 3) Filter all singletons and doubletons, then add back in those that passed criteria
KEEP <- rownames(present_absent)[which(rowSums(present_absent) >= 3)]
KEEP <- c(KEEP, KEEP_singletons, KEEP_doubletons)
p_filtered <- prune_taxa(KEEP, p)

print(paste("The Percent of Sequences Remaining After Filtering is",round((sum(sample_sums(p_filtered))/total_seqs)*100, 2),sep=" "))
print(paste("The Number of OTUs was Reduced from", total_otus,"to", length(taxa_names(p_filtered)), "(i.e.", round((length(taxa_names(p_filtered))/total_otus)*100,1),"% of OTUs remain)", sep=" "))

saveRDS(p_filtered, file = "raw_data/p_merged.final.filter.rds")  

"The Percent of Sequences Remaining After Removing Contaminants Found in Blanks is 99.55"
"The Number of OTUs was Reduced from 16207 to 16177 (i.e. 99.8 % of OTUs remain)"

"The Percent of Sequences Remaining After Removing Low Abundance Libraries is 99.47"
"The Number of OTUs were unchanged (i.e. 99.8 % of OTUs remain)"

"The Percent of Sequences Remaining After Filtering is 98.5"
"The Number of OTUs was Reduced from 16207 to 6680 (i.e. 41.2 % of OTUs remain)"

##################################################
# Normalize Abundances - Counts Per Thousand Reads

p_filtered <- readRDS(file = "raw_data/p_merged.final.filter.rds")  

x.totals <- colSums(otu_table(p_filtered))/1000

for (i in 1:length(x.totals)){
  otu_table(p_filtered)[,i] <- otu_table(p_filtered)[,i]/x.totals[i]
}

# Filter out C13 Control Samples
p_blanks <- prune_samples(sample_names(p_filtered)[which(sample_data(p_filtered)$rep == "Standard")],p_filtered)
p_filtered <- prune_samples(sample_names(p_filtered)[which(sample_data(p_filtered)$rep != "Standard")],p_filtered)

saveRDS(p_filtered, file = "raw_data/p_merged.final.filter.norm.rds")  
saveRDS(p_blanks, file = "raw_data/p_blanks.final.filter.norm.rds")  


###################
## Sample Breakdown

breakdown <- as.data.frame(sample_data(p))
breakdown$count <- 1

# By Treatments
print(ddply(breakdown, ~ tree + soil.type + enrichment, summarise, total = sum(count)))

# By Fraction
print(ddply(breakdown, ~ enrichment + fraction.type, summarise, total = sum(count)))
```
