## Supplementary material for "Phenolic acid-degrading *Paraburkholderia* prime decomposition in forest soil": BLOCK01c - in situ - Figure 4A - ANOSIM, NMDS and EnvFit.R

```
library(phyloseq)
library(plyr)
library(ggplot2)
library(reshape)
library(vegan)

#######################################################################
# NMDS + Fitted Environmental Variables for Microbiome Data (Figure 4A)
#######################################################################
set.seed(70)

# Import
p <- readRDS(file = "raw_data/p_merged.final.filter.norm.rds") 

# Average all gradient fractions for 13C-PHB and 12C-PHB (controls)
sample_data(p)$fullCombo <- paste(sample_data(p)$Combo, sample_data(p)$enrichment, sample_data(p)$rep, sep = "_")
p <- merge_samples(p, "fullCombo", fun = mean)
sample_data(p)[,c("Combo","fullCombo","depth","env_biome","run.name","derito.label","BarcodeSequence","LinkerPrimerSequence", "tree", "soil.type", "enrichment", "fraction.type", "fraction","soil.collection", "run.data", "Original.barcode")] <- list(NULL)
sample_data(p)$tree <- unlist(lapply(sample_names(p), function(x) paste(unlist(strsplit(x, "_"))[1:2], collapse="_")))
sample_data(p)$soil.type <- unlist(lapply(sample_names(p), function(x) unlist(strsplit(x, "_"))[3]))
sample_data(p)$enrichment <- unlist(lapply(sample_names(p), function(x) unlist(strsplit(x, "_"))[4]))
p <- subset_taxa(p, taxa_sums(p) > 0)

# Ordinate
ord <- ordinate(p, "NMDS", "bray", ~ tree)
NMDS = data.frame(ord$points, stringsAsFactors = F)

# Prep Environmental Parameters
design <- as.data.frame(as(sample_data(p), "matrix"), stringsAsFactors = F)
NMDS <- cbind(NMDS, design)
predictors <- readRDS(file = "raw_data/major.soilchem.predictors.rds") # from BLOCK03
predictors <- append(predictors, c("Nickel","Sulfur","Manganese"))
design <- design[,predictors]
design[] <- lapply(design, function(x) as.numeric(as.character(x)))

# Fit Environmental Parameters
fit <- envfit(ord, design, perm = 999, na.rm = TRUE)
vec<-as.data.frame(scores(fit, display = "vectors"))
vec$species<-rownames(vec)

# Plot NMDS
NMDS$tree <- as.factor(NMDS$tree)
saveRDS(NMDS, file = "raw_data/NMDS.coordinates.rds") # Save for BLOCK01d - perMANOVA

# Plot in ggplot
plot <- ggplot(NMDS, aes(MDS1, MDS2, color = tree, shape = soil.type)) + geom_point()
plot
ggsave(plot, filename="NMDS.ecoplots.pdf", height=6, width=8)

# Plot with basic R graphics
plot(ord)
plot(fit)


###########################################
# NMDS for Soil Chem Data (Inset Figure 4A)
###########################################

soil.chem <- as.data.frame(as(sample_data(readRDS(file = "raw_data/p_merged.final.filter.norm.rds") ), "matrix"), stringsAsFactors = F)
soil.chem.data <- c("tree", "soil.type", "rep", colnames(soil.chem)[18:51])
soil.chem <- ddply(soil.chem, as.formula(paste("~",paste(soil.chem.data,collapse="+"))), summarise, count = 1)
soil.chem$count <- NULL
soil.chem[,3:ncol(soil.chem)] <- lapply(soil.chem[,3:ncol(soil.chem)], as.numeric)

# Calculate distance
dist <- vegdist(soil.chem[,4:ncol(soil.chem)], method = "bray")

# Perform ANOSIM

# soil type
summary(anosim(dist, soil.chem[,c("soil.type")]))

# forest (for LnC)
summary(anosim(vegdist(subset(soil.chem, soil.type == "LnC")[,4:ncol(soil.chem)], method = "bray"), subset(soil.chem, soil.type == "LnC")[,c("tree")]))

# Perform NMDS
ord <- metaMDS(dist)
ord <- as.data.frame(ord$points)
plot_me <- cbind(soil.chem[,1:3], ord)
plot <- ggplot(plot_me, aes(MDS1, MDS2, color = tree, shape = soil.type)) + geom_point()
plot

ggsave(plot, filename="NMDS.tree.plot.pdf", width= 11, height=8.5)
```
