## Supplementary material for "Phenolic acid-degrading *Paraburkholderia* prime decomposition in forest soil": BLOCK01d - in situ - Figure 4B - Analysis of Soil Chem Data.R

```
library(phyloseq)
library(plyr)
library(ggplot2)
library(reshape)

###################################
## Plotting Environmental Variables (Figure 1B)
###################################

# User Input
make_plot <- "Y"    # "Y" | "N" (if you wish to plot boxplots of each soil chem variable)

# Import Data
p <- readRDS(file = "raw_data/p_merged.final.filter.norm.rds") 

# Prep Data
x <- as.data.frame(as(sample_data(p), "matrix"), stringsAsFactors = F)
x$Combo <- paste(x$Combo, x$rep, sep="_")
x <- x[-which(duplicated(x$Combo)),]
x <- x[,c(3:5,16,18:ncol(x)-1)]
x$depth <- NULL
x[,4:ncol(x)] <- lapply(x[,4:ncol(x)], function(x) as.numeric(as.character(x)))
x <- melt(x, id.vars = c("tree","soil.type","enrichment","rep"), value.factor = FALSE)

# Plot Data
variables <- unique(x$variable)

count = 1
for (var in variables){
  
  # Subset
  plot_me <- subset(x, variable == var)
  
  # Set Factor Levels
  plot_me$soil.type <- factor(plot_me$soil.type, levels = c("LnC", "MaB"))
  plot_me$tree <- factor(plot_me$tree, levels = c("Sugar_Maple", "Black_Locust","Red_Pine"))

  # Stats
  plot_me$tree <- paste(plot_me$tree, plot_me$soil.type, sep="_")
  
  plot_me$tree <- factor(plot_me$tree, levels = c("Sugar_Maple_LnC","Sugar_Maple_MaB", "Black_Locust_LnC", "Black_Locust_MaB","Red_Pine_LnC"))
  
  fit <- lm(value ~ tree, data = plot_me)
  fit <- TukeyHSD(aov(fit))
  
  if (!is.nan(fit$tree[,"p adj"])){
    # Do stats
    if (any(fit$tree[,"p adj"] < 0.05)){
      foo <- data.frame(fit$tree)
      foo <- subset(foo, p.adj < 0.05)
      foo$contrast1 <- unlist(lapply(row.names(foo), function(x) unlist(strsplit(x,"-"))[1]))
      foo$contrast2 <- unlist(lapply(row.names(foo), function(x) unlist(strsplit(x,"-"))[2]))
      foo <- foo[,c("contrast1","contrast2","p.adj")]
      foo$variable <- var
      
      if (count == 1){
        results <- foo
        count = count + 1
      } else {
        results <- rbind(results, foo)
      }
    }
  }

  # Plot
  if (make_plot == "Y"){
    plot <- ggplot(plot_me, aes(x=tree, y=value, fill = tree)) + geom_boxplot(outlier.shape = NA) + ylab("Units") + ggtitle(var)
    
    if (first_pass == "Y"){
      plot <- plot + facet_grid(~soil.type)
    }
    
    print(plot)
    
    writeLines("Save plots?")
    switch(menu(c("Discard","Continue","Exit")), step1 <- "Discard", step1 <- "Save to File", step1 <- "Exit")
    
    if (step1 == "Exit"){
      break
    } else if (step1 == "Save to File"){
      ggsave(plot, filename=paste("chem.data",var,'barplot.pdf',sep="."), height=16, width=16)
    }
  }
}

# Output Soil Chem Data as Supplementary Table
x.std <- ddply(x, ~ tree + soil.type + variable, summarise, std_error = std(value))
x.mean <- ddply(x, ~ tree + soil.type + variable, summarise, average = mean(value))
x <- merge(x.mean, x.std, by = c("tree","soil.type","variable"))
x$average <- round(x$average, 2)
x$std_error <- round(x$std_error, 2)
x <- reshape(x, direction = "wide", idvar = c("variable","soil.type"), timevar = "tree")
x$RedPine <- paste(x$average.Red_Pine, x$std_error.Red_Pine, sep=" ± ")
x$BlackLocust <- paste(x$average.Black_Locust, x$std_error.Black_Locust, sep=" ± ")
x$SugarMaple <- paste(x$average.Sugar_Maple, x$std_error.Sugar_Maple, sep=" ± ")

x <- x[,c("soil.type","variable","SugarMaple","BlackLocust","RedPine")]

# Save Output
write.csv(x, file = "Table SX - All Soil Chem Data.csv", row.names = F)
write.csv(results, file = "Table SX - All Soil Chem Data.statistics.csv", row.names = F)
```
