## Supplementary material for "Phenolic acid-degrading *Paraburkholderia* prime decomposition in forest soil": BLOCK01e - in situ - perMANOVA.R

```
library(phyloseq)
library(plyr)
library(vegan)
library(glmnet)
library(stats)

veganotu <- function(physeq) {
  OTU <- otu_table(physeq)
  
  if (taxa_are_rows(OTU)) {
    OTU <- t(OTU)
  }
  
  OTU <- as(OTU, "matrix")
  return(OTU)
}


########################################
## Variable Selection for Soil Chem Data
########################################

# Import ordination data (from BLOCK02)
NMDS <- readRDS(file = "raw_data/NMDS.coordinates.rds") 
MDS1 <- as.numeric(NMDS$MDS1)
MDS2 <- as.numeric(NMDS$MDS2)
NMDS[,c("rep","tree","enrichment","soil.type","MDS1","MDS2", "moisture", "LOI", "latitude", "longitude", "core.mass", "core.rocks", "hCore", "volume", "density")] <- list(NULL)
NMDS <- sapply(NMDS, as.numeric)

# stepwise regression
form <- as.formula(paste("MDS1 ~",paste(colnames(NMDS), collapse="+"),sep=""))
model <- lm(form, data = data.frame(cbind(MDS1,NMDS)))
NMDS1.model <- step(model)

form <- as.formula(paste("MDS2 ~",paste(colnames(NMDS), collapse="+"),sep=""))
model <- lm(form, data = data.frame(cbind(MDS2,NMDS)))
NMDS2.model <- step(model)

predictors.step <- unique(c(names(coef(NMDS1.model)), names(coef(NMDS2.model))))[-1]

############
## perMANOVA (Table S3) 
############

method = "bray" # bray or theta_YC
predictors <- "soil.chem" # "factors" | "soil.chem"

# Import
p <- readRDS(file = "raw_data/p_merged.final.filter.norm.rds") 

# Average all gradient fractions for 13C-PHB and 12C-PHB (controls)
sample_data(p)$fullCombo <- paste(sample_data(p)$Combo, sample_data(p)$enrichment, sample_data(p)$rep, sep = "_")
p <- merge_samples(p, "fullCombo", fun = mean)
sample_data(p)[,c("Combo","fullCombo","depth","env_biome","run.name","derito.label","BarcodeSequence","LinkerPrimerSequence", "tree", "soil.type", "enrichment", "fraction.type", "fraction","soil.collection", "run.data", "Original.barcode")] <- list(NULL)
sample_data(p)$tree <- unlist(lapply(sample_names(p), function(x) paste(unlist(strsplit(x, "_"))[1:2], collapse="_")))
sample_data(p)$soil.type <- unlist(lapply(sample_names(p), function(x) unlist(strsplit(x, "_"))[3]))
sample_data(p)$enrichment <- unlist(lapply(sample_names(p), function(x) unlist(strsplit(x, "_"))[4]))
p <- subset_taxa(p, taxa_sums(p) > 0)

# Make Counts Object
counts <- as.data.frame(as(t(otu_table(p)), "matrix"))
counts<-data.frame(veganotu(p))

# Get predictors
if (predictors == "soil.chem"){
 
  design <- as.data.frame(as(sample_data(p), "matrix"), stringsAsFactors = F)
  design <- data.frame(design[,c("enrichment",predictors.step)])
  design[,predictors.step] <- sapply(design[,predictors.step], as.numeric)
  
  if (method == "theta_YC"){
    form <- as.formula(paste("dis ~",paste(c(predictors.step), collapse="+"),sep=""))
  } else {
    form <- as.formula(paste("counts ~",paste(c(predictors.step), collapse="+"),sep=""))
  }
} else {
  design <- as.data.frame(as(sample_data(p), "matrix"), stringsAsFactors = F)
  design <- design[,c("tree","soil.type","enrichment")]
  
  if (method == "theta_YC"){
    form <- as.formula(paste("dis ~",paste(c("tree","soil.type","enrichment"), collapse="+"),sep=""))
  } else {
    form <- as.formula(paste("counts ~",paste(c("tree","soil.type","enrichment"), collapse="+"),sep=""))
  }
}

if (method == "theta_YC"){
  dis <- designdist(counts, method="1-(J/(A+B-J))",terms = c("quadratic"), abcd = FALSE)
  permanova <- adonis(form, design, permutations = 1000)
} else {
  permanova <- adonis(form, design, method = "bray", permutations = 1000)
}

if (predictors == "soil.chem"){
  names <-as.vector(c(attr(permanova$terms,"term.labels")[c(1:14)],"Residuals"))
  data <- permanova$aov.tab$R2[c(1:15)]
  prob <- permanova$aov.tab$Pr[c(1:15)]
} else {
  names <-as.vector(c(attr(permanova$terms,"term.labels")[c(1:3)],"Residuals"))
  data <- permanova$aov.tab$R2[c(1:4)]
  prob <- permanova$aov.tab$Pr[c(1:4)]
}

permanova.data <- data.frame(Factors = names, R_squared=data, p.value = prob, stringsAsFactors =FALSE)

# Save output
write.csv(permanova.data, file = paste("permanova",predictors,"info.csv", sep="."), row.names = F)
```
