## Supplementary material for "Phenolic acid-degrading *Paraburkholderia* prime decomposition in forest soil": BLOCK01f - in situ - Figure S6 - Diversity and Richness.R

```
library(phyloseq)
library(plyr)
library(ggplot2)
library(vegan)
library(ranacapa)

###############
##  Rarefaction (Supplementary Data)
###############

SIP <- "C13" # "C13" | "C12"
p <- subset_samples(readRDS(file = "p_merged.final.filter.rds"), enrichment == "C13")
sample_data(p)$combo <- paste(sample_data(p)$tree, sample_data(p)$soil.type, sep="_")
plot <- ggrare(p, step = 10, parallel = TRUE, color = "enrichment", label = "Sample")
plot

########################
##  Diversity & Richness (Figure S3)
########################
permutations = 100
indices = c("Observed", "Chao1", "ACE", "Shannon", "Simpson", "InvSimpson")
p <- readRDS(file = "raw_data/p_merged.final.filter.rds") 

# Subset (trend stays the same even if you subsample to the lowest fractions)
#x.data <- subset(x.data, variable == index & rep != "Standard")
p <- subset_samples(p, fraction %in% c("3-4","5","6","7","8"))
p <- subset_samples(p, rep != "Standard")

# Merge all fractions by rep
sample_data(p)$ComboID <- paste(sample_data(p)$enrichment, sample_data(p)$soil.type, sample_data(p)$tree, sample_data(p)$rep, sep="_")
p <- merge_samples(p, "ComboID")
sample_data(p)$ComboID <- NULL
sample_data(p)$Combo <- NULL

sample_data(p)$enrichment <- unlist(lapply(sample_names(p), function(x) unlist(strsplit(as.character(x), "_"))[1]))
sample_data(p)$soil.type <- unlist(lapply(sample_names(p), function(x) unlist(strsplit(as.character(x), "_"))[2]))
sample_data(p)$tree <- unlist(lapply(sample_names(p), function(x) paste(unlist(strsplit(as.character(x), "_"))[3:4], collapse="_")))
sample_data(p)$rep <- unlist(lapply(sample_names(p), function(x) unlist(strsplit(as.character(x), "_"))[5]))

saveRDS(p, file = "raw_data/p_merged.final.filter.repsAsOne.rds") # For BLOCK01f
#sample_sums(p)[order(sample_sums(p))] # Min. read depth = 9141

# Perform permutations of data
count = 1
for (i in 1:permutations){
  foo <- rarefy_even_depth(p)
  foo <- plot_richness(foo, c("enrichment"), measures=indices)  #Note: it doesn't matter that "treatment" is specified. All sample data is preserved
  
  if (count == 1){
    x.data <- foo$data
    x.data$perm <- i
  } else {
    foodata <- foo$data
    foodata$perm <- i
    x.data <- rbind(x.data, foodata)
  }
  count = count + 1
  print(count)
}


## Add in Pielou's Evenness Metric (done after the main calculation run)
# Grab data needed for calculation
observed <- subset(x.data, variable == "Observed")
shannon <- subset(x.data, variable == "Shannon")

#any(shannon$samples != observed$samples)  #sanity check = should be FALSE

# Calculate Evenness
J <- shannon$value/log(observed$value)  # Pielou's evenness
pielou <- shannon
pielou$variable <- "Pielou"
pielou$value <- J
pielou$se <- NA

# Add back to main dataframe  
x.data <- rbind(x.data, pielou)


########
# Plot 
########

#### USER INPUT : choose diversity metric
index = "Pielou" # options include : "Observed", "Chao1", "ACE", "Shannon", "Simpson", "InvSimpson", "Pielou

x.data <- subset(readRDS(file = "all.diversity.estimates.rds"), variable == index)

# Average all estimates from the random rarefying step
plot_data <- ddply(x.data, ~ enrichment + soil.type + tree + rep, summarise, average = mean(value))

# Plot
plot_data$enrichment <- factor(plot_data$enrichment, levels = c("C12", "C13"))
plot_data$tree <- factor(plot_data$tree, levels = c("Sugar_Maple", "Black_Locust","Red_Pine"))
plot_data$soil.type <- factor(plot_data$soil.type, levels = c("LnC", "MaB"))

plot<-ggplot(plot_data, aes(enrichment, average)) + geom_boxplot() + facet_wrap(~ soil.type+tree) 
plot <- plot + geom_point(data=plot_data, aes(x=enrichment, average), position=position_dodge(0.5), alpha=0.5)
plot <- plot + theme_bw()
plot

# T-Tests
for (tree.type in c("Sugar_Maple","Black_Locust","Red_Pine")){
  for (soil in c("LnC","MaB")){
    foo <- subset(plot_data, tree == tree.type & soil.type == soil)
    print(paste("Testing",tree.type,"&",soil,sep=" "))
    print(t.test(average ~ enrichment, foo))
  }  
}
```
