## Supplementary material for "Phenolic acid-degrading *Paraburkholderia* prime decomposition in forest soil": BLOCK01g - in situ - Differential Abundance - DESeq.R

```
library(phyloseq)
library(plyr)
library(DESeq2)

# Calculate Standard Error
std <- function(x) { 
  NA_Count<-is.na(x)
  NA_Count<-length(NA_Count["TRUE"])
  sd(x, na.rm=T)/sqrt(length(x)-NA_Count)
}

p <- readRDS(file = "raw_data/p_merged.final.filter.repsAsOne.rds")  

############
## Run DESeq

# Apply stricter filter against low abundance taxa (Indicator species won't work with low abundance data anyways)
p <- prune_taxa(names(which(taxa_sums(p) > mean(sample_sums(p)) * 0.005)), p)

# Run Analysis According to Site (differing by treatment handling)
data_list <- list(c("Sugar_Maple","LnC"),c("Sugar_Maple","MaB"),c("Black_Locust","LnC"),c("Black_Locust","MaB"),c("Red_Pine","LnC"))

DESeq.count = 1
for (data in data_list){
  foo <- subset_samples(p, soil.type == data[2] & tree == data[1])

  # Prepare counts (note: DESeq calculates the geometric mean, which means all the zeros in my columns are problematic)
  counts <- as.data.frame(as(t(otu_table(foo)), "matrix"))
  counts<-counts+1
  
  # Pull Experimental data from phyloseq object
  design <- as.data.frame(as(sample_data(foo), "matrix"), stringsAsFactors = F)
  design$enrichment <- factor(design$enrichment, levels = c("C13","C12"))
  
  # Sanity check
  #rownames(design) == colnames(counts)
  
  #Run DESeq
  DE <- DESeqDataSetFromMatrix(counts, design, formula(~ enrichment))
  dds <- DESeq(DE)
  results<-results(dds)
  results<-as.data.frame(results[order(results$log2FoldChange),])
  results <- subset(results, log2FoldChange < -3)

  # Save results    
  if (nrow(results) > 0){
    # Save output
    if (DESeq.count == 1){
      DESeq.results <- data.frame(ASV = row.names(results), soil.type = data[2], tree = data[1], logFold = (results$log2FoldChange)*-1, stat = results$stat, results$padj)
      DESeq.count = DESeq.count + 1
    } else {
      DESeq.results <- rbind(DESeq.results, data.frame(ASV = row.names(results), soil.type = data[2], tree = data[1], logFold = (results$log2FoldChange)*-1, stat = results$stat, results$padj))
    }
  }
}

saveRDS(DESeq.results, file = "DESeq.results.rds") # For BLOCK01g
```
