## Supplementary material for "Phenolic acid-degrading *Paraburkholderia* prime decomposition in forest soil": BLOCK01h - in situ - Figure 5 - Plot 13C Enriched ASVs.R

```
library(phyloseq)
library(plyr)
library(Hmisc)
library(ggplot2)

std <- function(x) { 
  NA_Count<-is.na(x)
  NA_Count<-length(NA_Count["TRUE"])
  sd(x, na.rm=T)/sqrt(length(x)-NA_Count)
}

#########################################
## Script to create Figure 5 and Table S1

## Calculate Response Ratio

# Import
x <- readRDS(file = "raw_data/DESeq.results.rds")

## Note: All ASVs identified by DESeq were manually curated to remove those with high standard deviation, typically found in low abundances
## see BLOCK01h

# Subset to manually verified enriched taxa
x <- subset(x, ASV %in% subset(readRDS(file = "raw_data/manually.curated.ASVs.rds"), status == "keep")$ASV) # These were manually veryfied using BLOCK06 based on consistency in the distribution across the CsCl gradient
x <- subset(x, ASV != "c10fd97955188a1a50f144960cfbcae4") # Accidentally deemed 'keep'

# Prepare Count Data
p <- prune_taxa(as.character(unique(x$ASV)), readRDS(file = "raw_data/p_merged.final.filter.norm.rds"))
p <- subset_samples(p, fraction %in% c("3-4","5","6","7","8"))
p <- subset_samples(p, rep != "Standard")
p <- subset_samples(p, enrichment == "C13")
p <- psmelt(p)

# average across heavy fractions
se <- ddply(p, ~ OTU + tree + soil.type + Phylum + Class + Order + Family + Genus, summarise, se = std(Abundance))
counts <- ddply(p, ~ OTU + tree + soil.type + Phylum + Class + Order + Family + Genus, summarise, Counts = mean(Abundance))
counts$se <- se$se  
colnames(counts)[1] <- "ASV"
x <- merge(x, counts, by = c("ASV","tree","soil.type"))

# Identify both soils
both_soils <- intersect(names(which(table(x$ASV, x$soil.type)[,1] > 0)), names(which(table(x$ASV, x$soil.type)[,2] > 0)))

# Identify all forest types
sm <- names(which(table(x$ASV, x$tree)[,1] > 0))
bl <- names(which(table(x$ASV, x$tree)[,2] > 0))
rp <- names(which(table(x$ASV, x$tree)[,3] > 0))
all_trees <- intersect(intersect(sm, bl), rp)

# designate
x$tree <- as.character(x$tree)
x$special.designation <- 'None'
x$special.designation[which(x$ASV %in% both_soils)] <- "All soils"
x$special.designation[which(x$ASV %in% all_trees)] <- "All forests"
x$special.designation[which(x$ASV %in% intersect(all_trees, both_soils))] <- "All forests | All soils"
x$special.designation[which(x$special.designation == "None")] <- x$tree[which(x$special.designation == "None")]

# Fix 'All soils' to read 'both soils; multiple forests' and 'both soils; single forest'
x$special.designation[which(x$ASV == "4a1c8b52a15697844431efb7363adb0e")] <- "Black_Locust" # all soils; Black Locust - only
x$special.designation[which(x$ASV == "11bb9511e708c810f45cba04c0f2453e")] <- "Sugar_Maple" # all soils; Sugar Maple - only
x$special.designation[which(x$special.designation == "All soils")] <- "Multiple forests | soils"
x$special.designation[which(x$ASV == "5e7528996d60eceb90f1232c95156c07")] <- "Multiple forests | soils" 
x$special.designation[which(x$ASV == "1832af5a2d44517bfacd2ec08754fcae")] <- "Multiple forests | soils" 

# Write output
write.csv(x, file = "Table SX - All enrASV.csv", row.names = F)

###############################################################################
## Barplot for ASVs with greater than 3 log2fold change relative to 12C control (Figure 5A)
###############################################################################

x <- unique(x[,c("ASV","Genus","logFold","Counts","se","special.designation")])
x$ASV <- as.character(x$ASV)

# Summarise
plot_me <- ddply(x, ~ ASV + Genus + special.designation, summarise, response.ratio = mean(logFold))

# Order by response.ratio
plot_me <- plot_me[rev(order(plot_me$response.ratio)),]
plot_me$rank <- 1:nrow(plot_me)

# Remove putative and unclassified
plot_me$Genus <- gsub("putative ","",plot_me$Genus)
plot_me$Genus <- gsub("unclassifed","",plot_me$Genus)

#Plot
plot <- ggplot(plot_me, aes(x = rank, y = response.ratio, fill = Genus))
plot <- plot + geom_bar(position="dodge", stat="identity") + theme(axis.text.x = element_text(angle = 45, hjust = 1)) + geom_hline(yintercept=3)
plot <- plot + coord_cartesian(ylim=c(3,11))
plot

plot <- ggplot(plot_me, aes(x = rank, y = response.ratio, fill = special.designation))
plot <- plot + geom_bar(position="dodge", stat="identity") + theme(axis.text.x = element_text(angle = 45, hjust = 1))
plot <- plot + coord_cartesian(ylim=c(3,11))

subset(plot_me, ASV == "2fea268498044452fdd476b1a5322e47") # P. xenovorans ASV
subset(plot_me, ASV == "6b64ff37fff389a42eccf7387ffac8b9") # P. madseniana RP11 ASV

######################################
## Repeat for total counts (Figure 5B)

# Prepare Count Data
p <- prune_taxa(as.character(unique(x$ASV)), readRDS(file = "raw_data/p_merged.final.filter.norm.rds"))
p <- subset_samples(p, fraction %in% c("3-4","5","6","7","8"))
p <- subset_samples(p, rep != "Standard")
p <- subset_samples(p, enrichment == "C13")
p <- psmelt(p)

# Summarise
se <- ddply(p, ~ OTU + Genus, summarise, se = std(Abundance))
counts <- ddply(p, ~ OTU + Genus, summarise, abundance = mean(Abundance))
counts$se <- se$se  
colnames(counts)[1] <- "ASV"
plot_me <- merge(counts, unique(x[,c("ASV","special.designation")]), by = "ASV")

# Order by response.ratio
plot_me <- plot_me[rev(order(plot_me$abundance)),]
plot_me$rank <- 1:nrow(plot_me)

# Remove putative and unclassified
plot_me$Genus <- gsub("putative ","",plot_me$Genus)
plot_me$Genus <- gsub("unclassifed","",plot_me$Genus)

# Plot
plot <- ggplot(plot_me, aes(x = rank, y = abundance, fill = Genus))
plot <- plot + geom_bar(position="dodge", stat="identity") + theme(axis.text.x = element_text(angle = 45, hjust = 1))
limits <- aes(ymax = abundance + se, ymin=abundance - se)
plot <- plot + geom_errorbar(limits, position=position_dodge(0.9), width=0.25)
plot

subset(plot_me, ASV == "2fea268498044452fdd476b1a5322e47") # P. xenovorans ASV
subset(plot_me, ASV == "6b64ff37fff389a42eccf7387ffac8b9") # P. madseniana RP11 ASV

head(plot_me, 10)$ASV
ggsave(plot, filename=paste('rank.abundance.PHB.degraders.barplot.pdf',sep="."), height=6, width=8)

plot <- ggplot(plot_me, aes(x = rank, y = abundance, fill = special.designation))
plot <- plot + geom_bar(position="dodge", stat="identity") + theme(axis.text.x = element_text(angle = 45, hjust = 1))
plot
```
