## Supplementary material for "Phenolic acid-degrading *Paraburkholderia* prime decomposition in forest soil": BLOCK01i - in situ - Plot Any ASV or Taxon.R

```
library(phyloseq)
library(plyr)
library(reshape2)
library(ggplot2)

std <- function(x) { 
  NA_Count<-is.na(x)
  NA_Count<-length(NA_Count["TRUE"])
  sd(x, na.rm=T)/sqrt(length(x)-NA_Count)
}

limits <- aes(ymax = average + error, ymin=average - error)

###########################
## USER INPUT: Select Taxon

method = "ASV"   ## "Taxa" | "ASV" (determines whether to plot ASV abundance grouped by taxonomic classification or cycle through a set of ASVs provided by the user)
need = "manually.verify"  # If you would like to create a dataframe to store ASVs deemed to meet the threshold for enrichment.

if (method == "Taxa"){
  rank = "Genus"
  taxa ="Burkholderia" 
  ASV = "foo"
} else {
  ASV <- "6b64ff37fff389a42eccf7387ffac8b9"
  # RP11ASV: "6b64ff37fff389a42eccf7387ffac8b9"
}

##### Run script
##### 
p <- readRDS(file = "raw_data/p_merged.final.filter.norm.rds")  

if (length(ASV) == 1 | method == "Taxa"){
  
  # Get Data and subset to taxon of interest
  if (method == "Taxa"){
    taxa_table <- as(tax_table(p)[,rank], "vector")
    KEEP <- rownames(tax_table(p))[which(taxa_table == taxa)]
    p_taxa <- prune_taxa(KEEP, p)
  } else {
    p_taxa <- prune_taxa(ASV, p)
  }
  
  # Prep dataframe
  plot_me <- data.frame(Counts = sample_sums(p_taxa))
  plot_me <- cbind(plot_me, sample_data(p_taxa)[,c("enrichment","fraction","tree","soil.type")]) 
  
  # Calculate standard error and mean
  error1 <- ddply(plot_me, ~ enrichment + tree + soil.type,  summarise, error = std(Counts))
  plot_me1 <- ddply(plot_me, ~ enrichment + tree + soil.type, summarise, average = mean(Counts))
  plot_me1 <- cbind(plot_me1, error = error1$error)

  error2 <- ddply(plot_me, ~ enrichment + fraction + tree + soil.type,  summarise, error = std(Counts))
  plot_me2 <- ddply(plot_me, ~ enrichment + fraction + tree + soil.type, summarise, average = mean(Counts))
  plot_me2 <- cbind(plot_me2, error = error2$error)
  
  # Set Factor Levels
  plot_me1$enrichment <- factor(plot_me1$enrichment, levels = c("C12", "C13"))
  plot_me2$enrichment <- factor(plot_me2$enrichment, levels = c("C12", "C13"))
  plot_me1$soil.type <- factor(plot_me1$soil.type, levels = c("LnC", "MaB"))
  plot_me2$soil.type <- factor(plot_me2$soil.type, levels = c("LnC", "MaB"))
  plot_me1$tree <- factor(plot_me1$tree, levels = c("Sugar_Maple", "Black_Locust","Red_Pine"))
  plot_me2$tree <- factor(plot_me2$tree, levels = c("Sugar_Maple", "Black_Locust","Red_Pine"))
  plot_me2$fraction <- factor(plot_me2$fraction, levels = c("1-2","3-4","5","6","7","8","9-10","11-12"))
  
  if (method == "Taxa"){
    plot1 <- ggplot(plot_me1, aes(x=tree, y=average, fill = enrichment)) + geom_bar(position="dodge", stat="identity") + ylab("Average Counts per Thousand Reads") + ggtitle(paste("Abundance Among Treatment for Heavy Fraction for Taxa",taxa,sep=": ")) + facet_grid(~ soil.type)
    plot1 <- plot1 + geom_errorbar(limits, position=position_dodge(0.9), width=0.25)
    print(plot1)
    
    writeLines("Save plots?")
    switch(menu(c("Discard","Continue","Exit")), step1 <- "Discard", step1 <- "Save to File", step1 <- "Exit")
    
    if (step1 == "Exit"){
      break
    } else if (step1 == "Save to File"){
      ggsave(plot1, filename=paste(rank,taxa,'heavy.barplot.pdf',sep="."), height=16, width=16)
    }
    
    plot2 <- ggplot(plot_me2, aes(x=fraction, y=average, fill = enrichment)) + geom_bar(position="dodge", stat="identity") + ylab("Average Counts per Thousand Reads") + ggtitle(paste("Abundance Across Density Gradient for Taxa",taxa,sep=": ")) + facet_wrap(~ soil.type + tree)
    plot2 <- plot2 + geom_errorbar(limits, position=position_dodge(0.9), width=0.25)
    print(plot2)
    
    writeLines("Save plots?")
    switch(menu(c("Discard","Continue","Exit")), step1 <- "Discard", step1 <- "Save to File", step1 <- "Exit")
    
    if (step1 == "Exit"){
      break
    } else if (step1 == "Save to File"){
      ggsave(plot2, filename=paste(rank,taxa,'fractions.barplot.pdf',sep="."), height=16, width=16)  
    }
    
  } else {
    plot1 <- ggplot(plot_me1, aes(x=tree, y=average, fill = enrichment)) + geom_bar(position="dodge", stat="identity") + ylab("Average Counts per Thousand Reads") + ggtitle(paste(tax_table(p_taxa),collapse=";")) + facet_grid(~ soil.type)
    plot1 <- plot1 + geom_errorbar(limits, position=position_dodge(0.9), width=0.25)
    print(plot1)
    print(ASV)
    
    writeLines("Save plots?")
    switch(menu(c("Discard","Continue","Exit")), step1 <- "Discard", step1 <- "Save to File", step1 <- "Exit")
    
    if (step1 == "Exit"){
      break
    } else if (step1 == "Save to File"){
      ggsave(plot1, filename=paste(ASV,'heavy.barplot.pdf',sep="."), height=16, width=16)
    }

    plot2 <- ggplot(plot_me2, aes(x=fraction, y=average, fill = enrichment)) + geom_bar(position="dodge", stat="identity") + ylab("Average Counts per Thousand Reads") + ggtitle(paste(tax_table(p_taxa),collapse=";")) + facet_wrap(~ soil.type + tree)
    plot2 <- plot2 + geom_errorbar(limits, position=position_dodge(0.9), width=0.25)
    print(plot2)
    print(ASV)
    
    writeLines("Save plots?")
    switch(menu(c("Discard","Continue","Exit")), step1 <- "Discard", step1 <- "Save to File", step1 <- "Exit")
    
    if (step1 == "Exit"){
      break
    } else if (step1 == "Save to File"){
      ggsave(plot2, filename=paste(ASV,'fractions.barplot.pdf',sep="."), height=16, width=16)
    }
    
  }
} else {
  # Get Data
  p_taxa <- prune_taxa(ASV, p)
  
  if (need == "manually.verify"){
    count1 = 1
  }
  
  for (n in ASV){
    foo <- prune_taxa(n, p_taxa)
    
    # Prep dataframe
    plot_me <- data.frame(Counts = sample_sums(foo))
    plot_me <- cbind(plot_me, sample_data(p_taxa)[,c("enrichment","fraction","tree","soil.type")]) 
    
    # Calculate standard error and mean
    error1 <- ddply(plot_me, ~ enrichment + tree + soil.type,  summarise, error = std(Counts))
    plot_me1 <- ddply(plot_me, ~ enrichment + tree + soil.type, summarise, average = mean(Counts))
    plot_me1 <- cbind(plot_me1, error = error1$error)
    
    error2 <- ddply(plot_me, ~ enrichment + fraction + tree + soil.type,  summarise, error = std(Counts))
    plot_me2 <- ddply(plot_me, ~ enrichment + fraction + tree + soil.type, summarise, average = mean(Counts))
    plot_me2 <- cbind(plot_me2, error = error2$error)
    
    # Set Factor Levels
    plot_me1$enrichment <- factor(plot_me1$enrichment, levels = c("C12", "C13"))
    plot_me2$enrichment <- factor(plot_me2$enrichment, levels = c("C12", "C13"))
    plot_me1$soil.type <- factor(plot_me1$soil.type, levels = c("LnC", "MaB"))
    plot_me2$soil.type <- factor(plot_me2$soil.type, levels = c("LnC", "MaB"))
    plot_me1$tree <- factor(plot_me1$tree, levels = c("Sugar_Maple", "Black_Locust","Red_Pine"))
    plot_me2$tree <- factor(plot_me2$tree, levels = c("Sugar_Maple", "Black_Locust","Red_Pine"))
    plot_me2$fraction <- factor(plot_me2$fraction, levels = c("1-2","3-4","5","6","7","8","9-10","11-12"))
    
    plot1 <- ggplot(plot_me1, aes(x=tree, y=average, fill = enrichment)) + geom_bar(position="dodge", stat="identity") + ylab("Average Counts per Thousand Reads") + ggtitle(paste(tax_table(foo),collapse=";")) + facet_grid(~ soil.type)
    plot1 <- plot1 + geom_errorbar(limits, position=position_dodge(0.9), width=0.25)
    print(plot1)
    print(n)
    
    if (need == "manually.verify"){
      print("good luck")
      
    } else {
      writeLines("Save plots?")
      switch(menu(c("Discard","Continue","Exit")), step1 <- "Discard", step1 <- "Save to File", step1 <- "Exit")
      
      if (step1 == "Exit"){
        break
      } else if (step1 == "Save to File"){
        ggsave(plot1, filename=paste(n,'heavy.barplot.pdf',sep="."), height=16, width=16)
      }
      
    }
    
    
    plot2 <- ggplot(plot_me2, aes(x=fraction, y=average, fill = enrichment)) + geom_bar(position="dodge", stat="identity") + ylab("Average Counts per Thousand Reads") + ggtitle(paste(tax_table(foo),collapse=";")) + facet_wrap(~ soil.type + tree)
    plot2 <- plot2 + geom_errorbar(limits, position=position_dodge(0.9), width=0.25)
    print(plot2)
    
    if (need == "manually.verify"){
      writeLines("Legit enrichment?")
      switch(menu(c("No","Yes","Exit")), step1 <- "Discard", step1 <- "Save to File", step1 <- "Exit")
      
      if (step1 == "Exit"){
        break
      } else if (step1 == "Save to File"){
        if (count1 == 1){
          results <- data.frame(ASV = n, status = "keep")
        } else {
          results <- rbind(results, data.frame(ASV = n, status = "keep"))
        }
      } else {
        if (count1 == 1){
          results <- data.frame(ASV = n, status = "discard")
          count1 = count1 + 1
        } else {
          results <- rbind(results, data.frame(ASV = n, status = "discard"))
        }
      }
    } else {
      writeLines("Save plots?")
      switch(menu(c("Discard","Continue","Exit")), step1 <- "Discard", step1 <- "Save to File", step1 <- "Exit")
      
      if (step1 == "Exit"){
        break
      } else if (step1 == "Save to File"){
        ggsave(plot2, filename=paste(n,'fractions.barplot.pdf',sep="."), height=16, width=16)
      }
    }
  }
}
```
