## Supplementary material for "Phenolic acid-degrading *Paraburkholderia* prime decomposition in forest soil": BLOCK02a - in vitro - GCMS Data Import.R

```
library(plyr)
library(ggplot2)

### Calculate Mols C
calc_mol_C = function(ppm,volume.L){
  # moles (n) = PV / RT
  
  # x = ppm
  temp.K=294.261
  pressure.atm=1
  R=0.08206
  
  ppm = as.numeric(ppm)
  mol.volume = (pressure.atm * volume.L) / (R * temp.K)  # mol gas in container
  mol.CO2 = mol.volume * (ppm / 1000000)  
  mol.C <- mol.CO2  # fraction of mol as CO2 ~ mol C
  
  return(mol.C)
}

### Join Data and Time into Single POSIX stamp
time_converter <- function(start, time_series){
  return(as.numeric(difftime(time_series, start), units='hours'))
}

## Set parameters
# Vessel volume (in L) 
v.volume <- 0.12

# Concentrations of Standards (in ppm / based on the method described on the Buckley Lab Git Hub)
stds <- data.frame(std = c(0,1,2,3,4,5,6), runID = c('std0','std1','std2','std3','std4','std5','std6'), ppm = c(0, 855, 1701, 3422, 8510, 17227, 34483))

#######################################
# Import Sampling Events and GC/MS Data

# Import Sampling Times
events <- read.csv(file="raw_data/events.csv", stringsAsFactors = F, header=T)
events$Measurer <- NULL

# Combine Data and Time
events$AllTime <- as.POSIXct(NA)
for (n in 1:nrow(events)){
  if (!is.na(as.POSIXct(strptime(paste(events$Date[n],events$Time[n],sep=" "), '%d/%m/%y %R'), tz="EST"))){
    events$AllTime[n] = as.POSIXct(strptime(paste(events$Date[n],events$Time[n],sep=" "), '%d/%m/%y %R'), tz="EST")
  } else if (!is.na(as.POSIXct(strptime(paste(events$Date[n],events$Time[n],sep=" "), '%d/%m/%Y %R'), tz="EST"))) {
    events$AllTime[n] = as.POSIXct(strptime(paste(events$Date[n],events$Time[n],sep=" "), '%d/%m/%Y %R'), tz="EST")
  } else {
    events$AllTime[n] = as.POSIXct(strptime(paste(events$Date[n],events$Time[n],sep=" "), '%d-%m-%y %R'), tz="EST")  
  }  
}

# Calculate Time Durations
start <- subset(events, Event == "Start")
sampling_events <- subset(events, Event == "Gas Sampling")

for (project in unique(sampling_events$Project)){
  foo_start <- subset(start, Project == project)
  foo_events <- subset(sampling_events, Project == project)
  addtime <- time_converter(foo_start$AllTime, foo_events$AllTime)
  assign(project, data.frame(timepoint = foo_events$Timepoint, time = addtime, Batch = project, Event = "Measurement"))
}

sampling_events <- rbind(get("PHB-priming")) # and any other projects that were looped

# Import GC/MS Data
x <- read.table(file="raw_data/co2.table", sep="\t", stringsAsFactors = F, header=T)
x$Batch <-  'PHB-priming'

# Add in Times
x <- merge(x,sampling_events, by=c("Batch","timepoint"))

# Make Standard Curves
standards <- x[grep("std", x$runID),]
standards$rack_pos <- unlist(lapply(standards$runID, function(x) unlist(strsplit(x, "_"))[2]))
standards$runID <- unlist(lapply(standards$runID, function(x) unlist(strsplit(x, "_"))[1]))
standards <- merge(standards, stds, by = "runID")
standards <- subset(standards, ion != "TIC")

# correct ion45 std concentrations to reflect natural abundance (1.1% )
standards$ppm[which(standards$ion == "ion45")] <- standards$ppm[which(standards$ion == "ion45")]*0.011

# how many undetected peaks?
standards[which(is.na(standards$peak.area)),]
standards <- subset(standards, !(is.na(peak.area)))

# Visualize
ggplot(subset(standards, ion == "ion44"), aes(ppm, peak.area, color = rack_pos)) + geom_point() + geom_smooth(method=lm, se=F)
ggplot(subset(standards, ion == "ion45"), aes(ppm, peak.area, color = rack_pos)) + geom_point() + geom_smooth(method=lm, se=F)

## No Need to Correct for Drift. The standards run at the start and end of the run identical.

# Remove zero standard
standards <- subset(standards, std != 0)
ggplot(subset(standards, ion == "ion44"), aes(ppm, peak.area, color = rack_pos)) + geom_point() + geom_smooth(method=lm, se=F)
ggplot(subset(standards, ion == "ion45"), aes(ppm, peak.area, color = rack_pos)) + geom_point() + geom_smooth(method=lm, se=F)

# Calculate regression coefficients
m.44 <- coef(lm(ppm ~ peak.area, data = subset(standards, ion == "ion44")))[2]
b.44 <- coef(lm(ppm ~ peak.area, data = subset(standards, ion == "ion44")))[1]

m.45 <- coef(lm(ppm ~ peak.area, data = subset(standards, ion == "ion45")))[2]
b.45 <- coef(lm(ppm ~ peak.area, data = subset(standards, ion == "ion45")))[1]

## Calculate Sample Concentration
x <- x[-grep("std", x$runID),]
x <- subset(x, ion != "TIC")

# how many undetected peaks?
x[which(is.na(x$peak.area)),]

# Calculate
x$concentration[which(x$ion == "ion44")] <- (x$peak.area[which(x$ion == "ion44")]*m.44 + b.44)
x$concentration[which(x$ion == "ion45")] <- (x$peak.area[which(x$ion == "ion45")]*m.45 + b.45)

# Calculate mol
x$mol.C <- calc_mol_C(x[,"concentration"], v.volume)  # for a 120 mL serum vial

# Adjust for 1 missed day between T8 and T9
foo<-subset(x, timepoint == "T9")
foo$mol.C <- foo$mol.C*4/3 # Three days were recorded out of 4. Artificially adjust for that 4th day.
x <- rbind(subset(x, timepoint != "T9"), foo)

# Save output
saveRDS(x, file = "raw_data/final.co2.data.rds")
```
