## Supplementary material for "Phenolic acid-degrading *Paraburkholderia* prime decomposition in forest soil": BLOCK02b - in vitro - Calculate Cumulative CO2.R

```
library(plyr)
library(ggplot2)

# Import data
x.CO2 <- readRDS(file = "raw_data/final.co2.data.rds")
x.main <- read.csv(file = "raw_data/main.csv", stringsAsFactors = F, header=T)
colnames(x.main)[1] <- "runID"
x <- merge(x.main, x.CO2, by = "runID")

## interpolate missed sample by averaging (sample 13C-glucose rep 3 was missed in T3)
x$concentration[which(x$timepoint == "T3" & x$runID == "63" & x$ion == "ion44")] <- mean(subset(x, timepoint == "T3" & treatment == "C13_glucose" & tree == "Black Locust" & ion == "ion44" & replicate %in% c(1,2,4))$concentration)
x$peak.area[which(x$timepoint == "T3" & x$runID == "63" & x$ion == "ion44")] <- mean(subset(x, timepoint == "T3" & treatment == "C13_glucose" & tree == "Black Locust" & ion == "ion44" & replicate %in% c(1,2,4))$peak.area)
x$mol.C[which(x$timepoint == "T3" & x$runID == "63" & x$ion == "ion44")] <- mean(subset(x, timepoint == "T3" & treatment == "C13_glucose" & tree == "Black Locust" & ion == "ion44" & replicate %in% c(1,2,4))$mol.C)

x$concentration[which(x$timepoint == "T3" & x$runID == "63" & x$ion == "ion45")] <- mean(subset(x, timepoint == "T3" & treatment == "C13_glucose" & tree == "Black Locust" & ion == "ion45" & replicate %in% c(1,2,4))$concentration)
x$peak.area[which(x$timepoint == "T3" & x$runID == "63" & x$ion == "ion45")] <- mean(subset(x, timepoint == "T3" & treatment == "C13_glucose" & tree == "Black Locust" & ion == "ion45" & replicate %in% c(1,2,4))$peak.area)
x$mol.C[which(x$timepoint == "T3" & x$runID == "63" & x$ion == "ion45")] <- mean(subset(x, timepoint == "T3" & treatment == "C13_glucose" & tree == "Black Locust" & ion == "ion45" & replicate %in% c(1,2,4))$mol.C)

## Normalize respiration to dry wt. soil (and account for soil sampling immediately post T6 - took 1.5 hrs)
# set microcosms where no soil was removed to zero
x$wt.removed.T7[which(is.na(x$wt.removed.T7))] <- 0

## Update quantities of soil remaining in microcosm
# adjust wt. removed for water content
water.added <- (x$WHC[which(x$timepoint == "T7")]*0.6)*10
soil.added <- x$soil.wt[which(x$timepoint == "T7")]
percent.moisture <- soil.added/(water.added+soil.added)
dry.wt.soil.removed <- x$wt.removed.T7[which(x$timepoint == "T7")] - (x$wt.removed.T7[which(x$timepoint == "T7")]*percent.moisture)
dry.wt.soil.removed <- dry.wt.soil.removed*1.05   # adjust assuming the soil last 5% moisture

# sutract dry wt. removed from dry. wt added
x$soil.wt[which(x$timepoint == "T7")] <- x$soil.wt[which(x$timepoint == "T7")] - dry.wt.soil.removed

## Normalize to mols C. respired to dry wt. soil
x$mol.C.dry.wt <- x$mol.C/x$soil.wt

#ddply(subset(x, ion == "ion45" & treatment == "C13_PHB"), ~ treatment + tree + timepoint, summarise, average = mean(mol.C.dry.wt))

## Calculate Cumulative Respiration
count = 1
for (i in unique(x$ion)){
  for (sample in unique(x$runID)){
    foo <- subset(x, runID == sample & ion == i)  
    foo <- foo[order(foo$time),]
    foo$cum.mol.C <- cumsum(foo$mol.C.dry.wt)
    
    if (count == 1){
      cumulative <- foo
      count = count + 1
    } else{
      cumulative <- rbind(cumulative, foo)
    }
  }
}

saveRDS(cumulative, file = "raw_data/final.co2.cumulative.rds")
```
