## Supplementary material for "Phenolic acid-degrading *Paraburkholderia* prime decomposition in forest soil": BLOCK02c - in vitro - Figure 1B - Plot Cumulative CO2.R

```
library(plyr)
library(ggplot2)
source("summarySE.R")

# Import data
cumulative <- readRDS(file = "raw_data/final.co2.cumulative.rds")

# Summarize
plot_me <- summarySE(cumulative, measurevar="cum.mol.C", groupvars=c("time","ion","tree","treatment"))

# Add in dummy zeros
foo <- unique(plot_me[,c("ion","tree","treatment")])
foo$time <- 0
plot_me <- rbind.fill(plot_me, foo)
plot_me$cum.mol.C[which(plot_me$time == 0)] <- 0
plot_me$sd[which(plot_me$time == 0)] <- 0

## Plot cumulative respiration

# set factor order
plot_me$treatment <- factor(plot_me$treatment, levels = c("water_control","inhibitor_control", "C12_PHB__C13_RP11","C13_glucose","C13_PHB","C13_PHB_inhibited","C13_PHB__C12_RP11"))
  
# ion 44
plot <- ggplot(subset(plot_me, ion == "ion44"), aes(time, cum.mol.C, color=treatment)) + geom_point() + geom_line() + facet_grid(~tree) + ylab("cumulative mols 12C per g dry wt. soil")
plot <- plot + geom_errorbar(aes(ymin=cum.mol.C-sd, ymax=cum.mol.C+sd), width=.2, position=position_dodge(0.05))
plot <- plot + scale_color_manual(values=c("#fca503", "#94928f", "#ed0e6f", "#850eed", "#0eb9ed", "#0e51ed", "#e20eed"))
print(plot)

## inhibitor had no affect in soil
## salinity had a negative effect on cell amendments (i.e. lower than water control)

# ion 45

# adjust for natural abundance (should do prior to calculating mean / sd if planning to report this figure)
water <- subset(cumulative, treatment == "water_control")
water <- reshape(water[, c("ion","time","treatment", "tree", "replicate","cum.mol.C")], direction = "wide", idvar =c("time","tree","treatment","replicate"), timevar = "ion")
water$nat.abund <- water$cum.mol.C.ion45/water$cum.mol.C.ion44
nat.abund <- ddply(water, ~ tree, summarise, average.nat.abund = mean(nat.abund))

adjust_me <- subset(plot_me, ion == "ion44")
adjust_me[which(adjust_me$tree == "Black Locust"), c("cum.mol.C","sd","se")]  <- adjust_me[which(adjust_me$tree == "Black Locust"), c("cum.mol.C","sd","se")] * nat.abund[which(nat.abund$tree == "Black Locust"),"average.nat.abund"]
adjust_me[which(adjust_me$tree == "Sugar Maple"), c("cum.mol.C","sd","se")] <- adjust_me[which(adjust_me$tree == "Sugar Maple"), c("cum.mol.C","sd","se")] * nat.abund[which(nat.abund$tree == "Sugar Maple"),"average.nat.abund"]
adjust_me[which(adjust_me$tree == "Red Pine"), c("cum.mol.C","sd","se")] <- adjust_me[which(adjust_me$tree == "Red Pine"), c("cum.mol.C","sd","se")] * nat.abund[which(nat.abund$tree == "Red Pine"),"average.nat.abund"]
colnames(adjust_me)[6] <- "nat.abund.mol.C"

ion45 <- subset(plot_me, ion == "ion45")
ion45 <- merge(ion45, adjust_me[,c("time","treatment", "tree", "nat.abund.mol.C")], by = c("time","treatment", "tree"))
ion45$cum.mol.C <- ion45$cum.mol.C - ion45$nat.abund.mol.C

## Plot by mols respired
plot <- ggplot(ion45, aes(time, cum.mol.C, color=treatment)) + geom_point() + geom_line() + facet_grid(~tree) + ylab("cumulative mols 13C (adjusted for natural abundance) per g dry wt. soil")
plot <- plot + geom_errorbar(aes(ymin=cum.mol.C-sd, ymax=cum.mol.C+sd), width=.2, position=position_dodge(0.05))
plot <- plot + scale_color_manual(values=c("#fca503", "#94928f", "#ed0e6f", "#850eed", "#0eb9ed", "#0e51ed", "#e20eed"))
print(plot)

## Plot by percent substrate respired

# calculate mols substrate
mgC13 <- 0.5 * 0.175
substrate <- mgC13 / (13 * 1000) # 0.5 mg C / (13 mg C/mmol)

# calculate percentage of substrate
percent.ion45 <- ion45
percent.ion45$cum.mol.C <- (percent.ion45$cum.mol.C/substrate)*100
percent.ion45$sd <- (percent.ion45$sd/substrate)*100

# plot
plot <- ggplot(percent.ion45, aes(time, cum.mol.C, color=treatment)) + geom_point() + geom_line() + facet_grid(~tree) + ylab("cumulative percent substrate respired")
plot <- plot + geom_errorbar(aes(ymin=cum.mol.C-sd, ymax=cum.mol.C+sd), width=.2, position=position_dodge(0.05))
plot <- plot + scale_color_manual(values=c("#fca503", "#94928f", "#ed0e6f", "#850eed", "#0eb9ed", "#0e51ed", "#e20eed"))
print(plot)

## Addendum

# Ratio between "C13_PHB__C12_RP11" and "C13_PHB" is supposed to be 1:20 (C13_PHB was diluted 20-fold to make the former.)
ddply(ion45, ~ treatment + tree, summarise, maximum = max(cum.mol.C))

# Ratio of "C13_PHB" to "C13_glucose"
foo <- ddply(ion45, ~ treatment + tree, summarise, maximum = max(cum.mol.C))
foo <- subset(foo, treatment == "C13_glucose" | treatment == "C13_PHB")
foo <- reshape(foo, direction = "wide", idvar = 'tree', timevar = 'treatment')
foo$ratio <- foo$maximum.C13_PHB/foo$maximum.C13_glucose
```
