## Supplementary material for "Phenolic acid-degrading *Paraburkholderia* prime decomposition in forest soil": BLOCK02d - in vitro - Net Priming.R

```
library(plyr)
library(reshape2)
library(ggplot2)
source("calculate_ratio.R")

# Import data
cumulative <- readRDS(file = "raw_data/final.co2.cumulative.rds")
cumulative <- subset(cumulative, timepoint == "T6")  ## At end of priming effect (just before subsampling)
#cumulative <- subset(cumulative, time == max(cumulative$time)) ## Full time-course

## Calculate ratio of C12 : C13
ratio.glucose <- calculate_ratio("N")[1]
ratio.PHB <- calculate_ratio("N")[2]


###################################################################################################
### Adjust mols 12C respired for each treatment based on 13C respired from added substrates / cells

##
## Calculate natural abundance information from water-only treatment
water <- subset(cumulative, treatment == "water_control")
water <- reshape(water[,c("ion","treatment", "tree", "replicate","soil.wt","cum.mol.C")], direction = "wide", idvar =c("tree","treatment","replicate", "soil.wt"), timevar = "ion")
water$nat.abund <- water$cum.mol.C.ion45/water$cum.mol.C.ion44
nat.abund <- ddply(water, ~ tree, summarise, average.nat.abund = mean(nat.abund))
water$nat.abund <- NULL
assign("water_control", water)

##
## Adjust simple treatments: Glucose-only and PHB-only

for (treat in c("C13_glucose", "C13_PHB")){
  foo <- subset(cumulative, treatment == treat)
  
  # subtract natural abundance 13C from totals mols 13C
  foo <- reshape(foo[,c("ion","treatment", "tree", "replicate","soil.wt","cum.mol.C")], direction = "wide", idvar =c("tree","treatment","replicate", "soil.wt"), timevar = "ion")
  foo <- merge(foo, nat.abund, by = "tree")
  foo$cum.mol.C13.nat.abund <- foo$cum.mol.C.ion44 * foo$average.nat.abund
  foo$cum.mol.C13.adj <- foo$cum.mol.C.ion45 - foo$cum.mol.C13.nat.abund
  foo$average.nat.abund <- NULL
  
  # adjust mols 12C respired based on mols 12C of substrate addition (i.e. for every mol 13C we measure, there were X mols 12C added with the substrate)
  if (treat == "C13_glucose"){
    foo$cum.mol.C12.substrate   <- foo$cum.mol.C13.adj * ratio.glucose
  } else {
    foo$cum.mol.C12.substrate   <- foo$cum.mol.C13.adj * ratio.PHB
  }
  
  # adjust
  foo$cum.mol.C12.adj <- foo$cum.mol.C.ion44 - foo$cum.mol.C12.substrate
  
  # save output
  assign(treat, foo)
}

##
## Inhibitor treatments

# Inhibitor-only: designed to measure how much 12C is respired from inhibitor (expected to be marginal, possibly lower if inhibitor proves toxic)
# C13_PHB + Inhibitor: designed to measure whether a decrease in priming occurs when pobA is inhibited

# determine how many additional mols 12C are produced from addition of inhibitor
inhib.only <- subset(cumulative, treatment == "inhibitor_control" & ion == "ion44")
inhib.only <- ddply(inhib.only, ~ tree + treatment, summarise, average.C12 = mean(cum.mol.C))
water.only <- ddply(water, ~ tree + treatment, summarise, average.C12 = mean(cum.mol.C.ion44))
inhib.vs.water <- rbind(inhib.only, water.only)
inhib.vs.water <- reshape(inhib.vs.water, direction = "wide", idvar =c("tree"), timevar = "treatment")
inhib.vs.water$average.C12.inhibitor_control - inhib.vs.water$average.C12.water_control

# No need to adjust for inhibitor (differences are negligible, negative or unbelievably high which suggests it may have stimulated phenol degraders, and is no longer a representative control for the 12C respired from just the inhibtor)

## Adjust PHB+Inhibitor treatment
PHB.inhib <- subset(cumulative, treatment == "C13_PHB_inhibited")

# subtract natural abundance 13C from mols 13C
PHB.inhib <- reshape(PHB.inhib[,c("ion","treatment", "tree", "replicate","soil.wt","cum.mol.C")], direction = "wide", idvar =c("tree","treatment","replicate", "soil.wt"), timevar = "ion")

# subtract natural abundance 13C from mols 13C
PHB.inhib <- merge(PHB.inhib, nat.abund, by = "tree")
PHB.inhib$cum.mol.C13.nat.abund <- PHB.inhib$cum.mol.C.ion44 * PHB.inhib$average.nat.abund
PHB.inhib$cum.mol.C13.adj <- PHB.inhib$cum.mol.C.ion45 - PHB.inhib$cum.mol.C13.nat.abund
PHB.inhib$average.nat.abund <- NULL

# adjust mols 12C respired based on mols 12C content of substrate addition
PHB.inhib$cum.mol.C12.substrate <- PHB.inhib$cum.mol.C13.adj * ratio.PHB

# adjust
PHB.inhib$cum.mol.C12.adj <- PHB.inhib$cum.mol.C.ion44 - PHB.inhib$cum.mol.C12.substrate

assign("C13_PHB_inhibited", PHB.inhib)

##
## Cell-amendment treatments

##  Calculate 12C mols due to addition of stimulated RP11

# Account for small amount of 13C-PHB added for stimulation (1:20 dilution)
RP11.activity <- subset(cumulative, treatment == "C13_PHB__C12_RP11")

# subtract natural abundance 13C from mols 13C
RP11.activity <- reshape(RP11.activity[,c("ion","treatment", "tree", "replicate","soil.wt","cum.mol.C")], direction = "wide", idvar =c("tree","treatment","replicate", "soil.wt"), timevar = "ion")
RP11.activity <- merge(RP11.activity, nat.abund, by = "tree")
RP11.activity$cum.mol.C13.nat.abund <- RP11.activity$cum.mol.C.ion44 * RP11.activity$average.nat.abund
RP11.activity$cum.mol.C13.adj <- RP11.activity$cum.mol.C.ion45 - RP11.activity$cum.mol.C13.nat.abund
RP11.activity$average.nat.abund <- NULL

# adjust mols 12C respired based on mols 12C content of substrate addition
RP11.activity$cum.mol.C12.substrate   <- RP11.activity$cum.mol.C13.adj * ratio.PHB

# adjust
RP11.activity$cum.mol.C12.adj.penultimate <- RP11.activity$cum.mol.C.ion44 - RP11.activity$cum.mol.C12.substrate

## Adjust estimate based on 12C produced from cells
RP11.necromass <- subset(cumulative, treatment == "C12_PHB__C13_RP11")

# subtract natural abundance 13C from mols 13C
RP11.necromass <- reshape(RP11.necromass[,c("ion","treatment", "tree", "replicate","soil.wt","cum.mol.C")], direction = "wide", idvar =c("tree","treatment","replicate", "soil.wt"), timevar = "ion")
RP11.necromass <- merge(RP11.necromass, nat.abund, by = "tree")
RP11.necromass$cum.mol.C13.nat.abund <- RP11.necromass$cum.mol.C.ion44 * RP11.necromass$average.nat.abund
RP11.necromass$cum.mol.C13.adj <- RP11.necromass$cum.mol.C.ion45 - RP11.necromass$cum.mol.C13.nat.abund
RP11.necromass$average.nat.abund <- NULL

# adjust for 99 atom % glucose used to label cells
RP11.necromass$cum.mol.C13.adj.necromass <- RP11.necromass$cum.mol.C13.adj * 1.01

# adjust mols 12C respired by 12C from necromass
RP11.activity <- merge(RP11.activity, RP11.necromass[,c("tree","replicate","cum.mol.C13.adj.necromass")], by = c("tree","replicate"))
RP11.activity$cum.mol.C12.adj <- RP11.activity$cum.mol.C12.adj.penultimate - RP11.activity$cum.mol.C13.adj.necromass

assign("RP11.activity", RP11.activity)

##############
# Compile Data
x <- do.call(rbind.fill, lapply(c("water_control","C13_glucose", "C13_PHB","C13_PHB_inhibited","RP11.activity"), as.symbol))
x$cum.mol.C12.adj[which(x$treatment == "water_control")] <- x$cum.mol.C.ion44[which(x$treatment == "water_control")]


#########################################################
# Plot dot+box plots for all treatments vs. water control

# combine C13_PHB treatments
x$treatment[which(x$treatment == "C13_PHB_inhibited")] <- "C13_PHB"
x$treatment <- factor(x$treatment, levels = c("water_control", "C13_glucose", "C13_PHB", "C13_PHB__C12_RP11"))

plot <- ggplot(x, aes(x=treatment, y=cum.mol.C12.adj, colour = treatment)) + geom_boxplot(outlier.shape = NA) + ylab("adjusted cumulative mol C12 per g dry wt. soil") + facet_grid(~tree)
plot <- plot + geom_point(position=position_jitterdodge(jitter.width = 0.1), alpha=0.5)
plot <- plot + scale_color_manual(values=c("#fca503", "#850eed", "#0eb9ed", "#e20eed"))
print(plot)

## Statistics
for (t in c("Sugar Maple","Black Locust","Red Pine")){
  print(paste("________",t,"________"))
  
  # Averages
  print(ddply(subset(x, tree == t), ~treatment, summarise, average = mean(cum.mol.C12.adj)))
  
  # Stats
  fit <- lm(cum.mol.C12.adj ~ treatment, data= subset(x, tree == t))
  print(TukeyHSD((aov(fit))))
  
}

########################################################
# Stats on net priming among ecoplots for PHB treatments
controls <- ddply(subset(x, treatment == "water_control"), ~ tree, summarize, mean.water = mean(cum.mol.C12.adj))
x.stat <- merge(subset(x, treatment == "C13_PHB")[,c("treatment","tree","cum.mol.C12.adj")], controls, by = "tree")
x.stat$net.prime <- x.stat$cum.mol.C12.adj-x.stat$mean.water
fit <- lm(net.prime ~ tree, data= x.stat)
print(TukeyHSD((aov(fit))))
ddply(x.stat, ~ tree, summarise, average = mean(net.prime))


#################################################
# Ratio of "C13_PHB__C12_RP11" to "water_control"
foo <- ddply(x, ~ treatment + tree, summarise, average = mean(cum.mol.C12.adj))
foo <- subset(foo, treatment == "C13_PHB__C12_RP11" | treatment == "water_control")
foo <- reshape(foo, direction = "wide", idvar = 'tree', timevar = 'treatment')
foo$ratio <- foo$average.C13_PHB__C12_RP11/foo$average.water_control
1-foo$ratio
```
