## Supplementary material for "Phenolic acid-degrading *Paraburkholderia* prime decomposition in forest soil": BLOCK02f - in vitro - Figure 1A - Instantaneous - Temporal Dynamics of Priming.R

```
library(plyr)
library(reshape2)
library(ggplot2)
source("summarySE.R")
source("calculate_ratio.R")

# Import data
spontaneous <- readRDS(file = "raw_data/final.co2.cumulative.rds")
spontaneous$cum.mol.C <- NULL

## Calculate ratio of C12 : C13
ratio.glucose <- calculate_ratio("N")[1]
ratio.PHB <- calculate_ratio("N")[2]


###################################################################################################
### Adjust mols 12C respired for each treatment based on 13C respired from added substrates / cells

##
## Calculate natural abundance information from water-only treatment
water <- subset(spontaneous, treatment == "water_control")
water <- reshape(water[,c("ion","time","treatment", "tree", "replicate","soil.wt","mol.C.dry.wt")], direction = "wide", idvar =c("time","tree","treatment","replicate", "soil.wt"), timevar = "ion")
water$nat.abund <- water$mol.C.dry.wt.ion45/water$mol.C.dry.wt.ion44
# ggplot(water, aes(nat.abund, time, color = tree)) + geom_point() + geom_smooth(method=lm, se=F) ## Intersting that natural abundance C is lower over time
nat.abund <- ddply(water, ~ tree + time, summarise, average.nat.abund = mean(nat.abund))
water$nat.abund <- NULL
assign("water_control", water)

##
## Inhibitor treatments

# Inhibitor-only: designed to measure how much 12C is respired from inhibitor (expected to be marginal, possibly lower if inhibitor proves toxic)
# C13_PHB + Inhibitor: designed to measure whether a decrease in priming occurs when pobA is inhibited

# determine how many additional mols 12C are produced from addition of inhibitor
inhib.only <- subset(spontaneous, treatment == "inhibitor_control" & ion == "ion44")
inhib.only <- ddply(inhib.only, ~ tree + treatment + time, summarise, average.C12 = mean(mol.C.dry.wt))
water.only <- ddply(water, ~ tree + treatment + time, summarise, average.C12 = mean(mol.C.dry.wt.ion44))
inhib.vs.water <- rbind(inhib.only, water.only)
inhib.vs.water <- reshape(inhib.vs.water, direction = "wide", idvar =c("time","tree"), timevar = "treatment")
inhib.vs.water$average.C12.inhibitor_control - inhib.vs.water$average.C12.water_control

# No need to adjust for inhibitor (differences are negligible, negative or unbelievably high which suggests it may have stimulated phenol degraders, and is no longer a representative control for the 12C respired from just the inhibtor)

##
## Adjust simple treatments: Glucose-only and PHB-only
for (treat in c("C13_glucose", "C13_PHB","C13_PHB_inhibited")){
  foo <- subset(spontaneous, treatment == treat)
  
  # subtract natural abundance 13C from totals mols 13C
  foo <- reshape(foo[,c("ion","time","treatment", "tree", "replicate","soil.wt","mol.C.dry.wt")], direction = "wide", idvar =c("time","tree","treatment","replicate", "soil.wt"), timevar = "ion")
  foo <- merge(foo, nat.abund, by = c("tree","time"))
  foo$mol.C.dry.wt.C13.nat.abund <- foo$mol.C.dry.wt.ion44 * foo$average.nat.abund
  foo$mol.C.dry.wt.C13.adj <- foo$mol.C.dry.wt.ion45 - foo$mol.C.dry.wt.C13.nat.abund
  foo$average.nat.abund <- NULL
  
  # adjust mols 12C respired based on mols 12C of substrate addition (i.e. for every mol 13C we measure, there were X mols 12C added with the substrate)
  if (treat == "C13_glucose"){
    foo$mol.C.dry.wt.C12.substrate   <- foo$mol.C.dry.wt.C13.adj * ratio.glucose
  } else {
    foo$mol.C.dry.wt.C12.substrate   <- foo$mol.C.dry.wt.C13.adj * ratio.PHB
  }

  # adjust
  foo$mol.C.dry.wt.C12.adj <- foo$mol.C.dry.wt.ion44 - foo$mol.C.dry.wt.C12.substrate
  
  # save output
  assign(treat, foo)
}

##
## Cell-amendment treatments

##  Calculate 12C mols due to addition of stimulated RP11

# Account for small amount of 13C-PHB added for stimulation (1:20 dilution)
RP11.activity <- subset(spontaneous, treatment == "C13_PHB__C12_RP11")

# subtract natural abundance 13C from mols 13C
RP11.activity <- reshape(RP11.activity[,c("ion","time","treatment", "tree", "replicate","soil.wt","mol.C.dry.wt")], direction = "wide", idvar =c("time","tree","treatment","replicate", "soil.wt"), timevar = "ion")
RP11.activity <- merge(RP11.activity, nat.abund, by = c("time","tree"))
RP11.activity$mol.C.dry.wt.C13.nat.abund <- RP11.activity$mol.C.dry.wt.ion44 * RP11.activity$average.nat.abund
RP11.activity$mol.C.dry.wt.C13.adj <- RP11.activity$mol.C.dry.wt.ion45 - RP11.activity$mol.C.dry.wt.C13.nat.abund
RP11.activity$average.nat.abund <- NULL

# adjust mols 12C respired based on mols 12C content of substrate addition
RP11.activity$mol.C.dry.wt.C12.substrate   <- RP11.activity$mol.C.dry.wt.C13.adj * ratio.PHB

# adjust
RP11.activity$mol.C.dry.wt.C12.adj.penultimate <- RP11.activity$mol.C.dry.wt.ion44 - RP11.activity$mol.C.dry.wt.C12.substrate

## Adjust estimate based on 12C produced from cells
RP11.necromass <- subset(spontaneous, treatment == "C12_PHB__C13_RP11")

# subtract natural abundance 13C from mols 13C
RP11.necromass <- reshape(RP11.necromass[,c("ion","time","treatment", "tree", "replicate","soil.wt","mol.C.dry.wt")], direction = "wide", idvar =c("tree","time","treatment","replicate", "soil.wt"), timevar = "ion")
RP11.necromass <- merge(RP11.necromass, nat.abund, by = c("time","tree"))
RP11.necromass$mol.C.dry.wt.C13.nat.abund <- RP11.necromass$mol.C.dry.wt.ion44 * RP11.necromass$average.nat.abund
RP11.necromass$mol.C.dry.wt.C13.adj <- RP11.necromass$mol.C.dry.wt.ion45 - RP11.necromass$mol.C.dry.wt.C13.nat.abund
RP11.necromass$average.nat.abund <- NULL

# adjust for 99 atom % glucose used to label cells
RP11.necromass$mol.C.dry.wt.C13.adj.necromass <- RP11.necromass$mol.C.dry.wt.C13.adj * 1.01

# adjust mols 12C respired by 12C from necromass
RP11.activity <- merge(RP11.activity, RP11.necromass[,c("time","tree","replicate","mol.C.dry.wt.C13.adj.necromass")], by = c("time","tree","replicate"))
RP11.activity$mol.C.dry.wt.C12.adj <- RP11.activity$mol.C.dry.wt.C12.adj.penultimate - RP11.activity$mol.C.dry.wt.C13.adj.necromass

assign("RP11.activity", RP11.activity)

##############
# Compile Data
x <- do.call(rbind.fill, lapply(c("water_control","C13_glucose", "C13_PHB","C13_PHB_inhibited","RP11.activity"), as.symbol))
x$mol.C.dry.wt.C12.adj[which(x$treatment == "water_control")] <- x$mol.C.dry.wt.ion44[which(x$treatment == "water_control")]

saveRDS(x, file = "raw_data/spontaneous.soil.priming.mols.12-CO2.rds")

###############################
# Plot spontaneous priming effect (mols 12C primed minus mols 12C in water-only control)

# Subtract average of water-only control per rep
water2 <- subset(x, treatment == "water_control")
water2 <- ddply(water2, ~ tree + time, summarise, control = mean(mol.C.dry.wt.C12.adj))
x <- merge(x, water2, by = c("tree","time"))
x$mol.C.dry.wt.C12.adj <- x$mol.C.dry.wt.C12.adj - x$control

# Summarize
plot_me <- summarySE(x, measurevar="mol.C.dry.wt.C12.adj", groupvars=c("time","tree","treatment"))

# Add in dummy zeros
foo <- unique(plot_me[,c("tree","treatment")])
foo$time <- 0
plot_me <- rbind.fill(plot_me, foo)
plot_me$mol.C.dry.wt.C12.adj[which(plot_me$time == 0)] <- 0
plot_me$sd[which(plot_me$time == 0)] <- 0

# remove water control from plot
plot_me <- subset(plot_me, treatment != "water_control")

# set factor order
#plot_me$treatment <- factor(plot_me$treatment, levels = c("water_control","C13_glucose","C13_PHB","C13_PHB_inhibited","C13_PHB__C12_RP11"))
plot_me$treatment <- factor(plot_me$treatment, levels = c("C13_glucose","C13_PHB","C13_PHB_inhibited","C13_PHB__C12_RP11"))

# plot
plot <- ggplot(plot_me, aes(time, mol.C.dry.wt.C12.adj, color=treatment)) + geom_point() + geom_line() + facet_grid(~tree) + ylab("spontaneous mols 12C primed per g dry wt. soil")
plot <- plot + geom_errorbar(aes(ymin=mol.C.dry.wt.C12.adj-sd, ymax=mol.C.dry.wt.C12.adj+sd), width=.2, position=position_dodge(0.05))
#plot <- plot + scale_color_manual(values=c("#fca503","#850eed", "#0eb9ed", "#0e51ed", "#e20eed"))
plot <- plot + scale_color_manual(values=c("#850eed", "#0eb9ed", "#0e51ed", "#e20eed"))

print(plot)
```
