## Supplementary material for "Phenolic acid-degrading *Paraburkholderia* prime decomposition in forest soil": BLOCK02g - in vitro - Statistics on Temporal Dynamics of Priming.R

```
library(plyr)

### USER INPUT
data <- "spontaneous"   # "spontaneous" | "cumulative"

# Import
x <- readRDS(file = paste("raw_data/",data,".soil.priming.mols.12-CO2.rds",sep="")) # From BLOCK02e

# run Tukey on all contrasts & only keep significant ones
count <- 1
for (t in unique(x$time)){
  for (trunk in unique(x$tree)){
    if (data == "spontaneous"){
      fit <- lm(mol.C.dry.wt.C12.adj ~ treatment, data=subset(x, time == t & tree == trunk))
    } else {
      fit <- lm(cum.mol.C12.adj ~ treatment, data=subset(x, time == t & tree == trunk))
    }
    
    fit <- TukeyHSD((aov(fit)))
    fit <- as.data.frame(fit$treatment)
    fit$tree <- trunk
    fit$time <- t

    if (count == 1){
      results <- subset(fit, `p adj` < 0.05)
      count = count + 1
    } else {
      results <- rbind(results, subset(fit, `p adj` < 0.05))
    }
  }
}

# simplify
results$contrast <- row.names(results)
results$contrast1 <- unlist(lapply(results$contrast, function(x) unlist(strsplit(x, "-"))[1]))
results$contrast2 <- unlist(lapply(results$contrast, function(x) unlist(strsplit(x, "-"))[2]))
results[,c("diff","lwr","upr","contrast")] <- list(NULL)
row.names(results)<-NULL
results <- results[,c(2:5,1)]
results <- results[order(results$tree),]

# results
results <- merge(results, unique(readRDS(file = "raw_data/final.co2.cumulative.rds")[,c("timepoint","time")]), by ="time")

# output all differences to water control
write.csv(subset(results, contrast1 == "water_control" | contrast2 == "water_control"), file = paste(data,"sig.diff.vs.control.csv",sep="."), row.names = F)
```
