## Supplementary material for "Phenolic acid-degrading *Paraburkholderia* prime decomposition in forest soil": BLOCK02h - in vitro - Amplicon Library Preprocessing.R

```
library(phyloseq)
library(plyr)

############################
## Merge Two Sequencing Runs
p.1 <- readRDS(file = "raw_data/p_priming.run1.rds")
p.2 <- readRDS(file = "raw_data/p_priming.run2.rds")

saveRDS(sample_names(p.1), file = "run1.sampleIDs.rds") # for making Figure 5

p.1 <- merge_phyloseq(otu_table(p.1), tax_table(p.1))
p.2 <- merge_phyloseq(otu_table(p.2), tax_table(p.2))

# Merge
p <- merge_phyloseq(p.1,p.2)

# Add in Sample Info
sample_file <- read.csv(file = "raw_data/in.vitro.sample.info.csv",header = T, stringsAsFactors = F)
rownames(sample_file) = sample_file$SampleID
sample_file$SampleID <- NULL

# Finalize
p_final <- merge_phyloseq(p, sample_data(sample_file))
p <- p_final

######################################################
## Pre-process to Filter Low Abundance and Sparse Taxa

# Keep information about the total # of sequences and OTUs
total_seqs <- sum(sample_sums(p))
total_otus <- length(taxa_names(p))

#######################################
## Filter Low Abundance and Sparse OTUs

# Set Filters
filter1 <- mean(sample_sums(p)) * 0.0005  #
filter2 <- mean(sample_sums(p)) * 0.01

# Hard Filter OTUs below 0.05% of each sample's Total Abundance (i.e. summed across all samples)
p <- prune_taxa(names(which(taxa_sums(p) > filter1)), p)

# Soft Filter OTUs which occur in less than three samples (with two exceptions)
present_absent<-otu_table(p) 
present_absent[present_absent > 0] <- 1

# 1) Remove all singletons except those with a max relative abundance greater than 1% in any sample
REMOVE_singletons <- rownames(present_absent)[which(rowSums(present_absent) == 1)]
p_assess <- prune_taxa(REMOVE_singletons, p)
KEEP_singletons <- names(which(apply(otu_table(p_assess), 1, function(x) max(x) > filter2)))
REMOVE_singletons <- setdiff(REMOVE_singletons, KEEP_singletons)

# 2) Remove all doubletons except those with a max relative abundance greater than 1% in any sample OR occur in the same forest and soil type
REMOVE_doubletons <- rownames(present_absent)[which(rowSums(present_absent) == 2)]
p_assess <- prune_taxa(REMOVE_doubletons, p)

KEEP_doubletons <- names(which(apply(otu_table(p_assess), 1, function(x) max(x) > filter2)))
CHECK_doubletons <- prune_taxa(names(which(apply(otu_table(p_assess), 1, function(x) max(x) < filter2))), p_assess)

KEEP_paired_doubles <- vector()
for (taxa in taxa_names(CHECK_doubletons)){
  foo <- prune_taxa(taxa, CHECK_doubletons)
  foo <- prune_samples(names(which(sample_sums(foo) > 0)), p)
  
  if (nrow(unique(sample_data(foo)[,"tree"])) == 1) {
    KEEP_paired_doubles <- c(KEEP_paired_doubles, taxa)
  }
}

KEEP_doubletons <- c(KEEP_doubletons, KEEP_paired_doubles)

# 3) Filter all singletons and doubletons, then add back in those that passed criteria
KEEP <- rownames(present_absent)[which(rowSums(present_absent) >= 3)]
KEEP <- c(KEEP, KEEP_singletons, KEEP_doubletons)
p_filtered <- prune_taxa(KEEP, p)

print(paste("The Percent of Sequences Remaining After Filtering is",round((sum(sample_sums(p_filtered))/total_seqs)*100, 2),sep=" "))
print(paste("The Number of OTUs was Reduced from", total_otus,"to", length(taxa_names(p_filtered)), "(i.e.", round((length(taxa_names(p_filtered))/total_otus)*100,1),"% of OTUs remain)", sep=" "))

saveRDS(p_filtered, file = "raw_data/p_priming.final.filter.rds")  

"The Percent of Sequences Remaining After Filtering is 97.69"
"The Number of OTUs was Reduced from 5137 to 1541 (i.e. 30 % of OTUs remain)"


##################################################
# Normalize Abundances - Counts Per Thousand Reads

p_filtered <- readRDS(file = "raw_data/p_priming.final.filter.rds")  

x.totals <- colSums(otu_table(p_filtered))/1000

for (i in 1:length(x.totals)){
  otu_table(p_filtered)[,i] <- otu_table(p_filtered)[,i]/x.totals[i]
}

saveRDS(p_filtered, file = "raw_data/p_priming.final.filter.norm.rds")  
```
