## Supplementary material for "Phenolic acid-degrading *Paraburkholderia* prime decomposition in forest soil": BLOCK02i - in vitro - Indicator Analysis.R

```
library(indicspecies)
library(phyloseq)
library(plyr)
library(reshape2)

# Import data
p <- readRDS(file = "raw_data/p_priming.final.filter.norm.rds")  

# Combine PHB and PHB+inhibitor treatments
sample_data(p)[which(sample_data(p)[,"treatment"] == "C13_PHB_inhibited"), "treatment"] <- "C13_PHB"

# Remove 'field samples' and cell amendments (uninformative for this analysis)
p <- subset_samples(p, treatment != "field" & treatment != "C13_PHB__C12_RP11")

# Remove cDNA samples (activity had subsided by day 7)
p <- subset_samples(p, amplicon == "DNA")

# Remove sequencing data from first pass (all trends are consistent, but counts are 2-fold higher across the board)
p <- subset_samples(p, !(sample_names(p) %in% readRDS(file = "run1.sampleIDs.rds")))


###################
## Indicator by ASV

factor <- "treatment"

count = 1
for (ecoplot in c("Sugar Maple","Red Pine","Black Locust")){
  foo <- subset_samples(p, tree == ecoplot)
  foo <- subset_taxa(foo, taxa_sums(foo) > 0)
  
  # Separate Metadata
  design <- as.data.frame(as(sample_data(foo), "matrix"), stringsAsFactors = F)
  groups <- design[,factor]
  
  # Make Counts Matrix
  counts <- as.data.frame(as(t(otu_table(foo)), "matrix"))
  
  # Remove NA
  if (any(is.na(groups))){
    counts <- counts[-which(is.na(groups)),]
    groups <- groups[-which(is.na(groups))]
    
  }
  
  # Run Indicator Speices
  indval = multipatt(counts, groups, control = how(nperm=999))
  
  # Store Results in Table
  results <- indval$sign
  
  # Remove NA
  if(length(which(is.na(results$p.value)))>0){
    results <- results[-which(is.na(results$p.value)),]  
  }
  
  # Filter for signficance
  #results <- results[which(results$p.value < 0.2),] # permutations of 999 gives p-value of 0.001, which is adjusted to 0.06 (i.e. above threshold).
  results <- results[which(results$stat > 0.8),] # permutations of 999 gives p-value of 0.001, which is adjusted to 0.06 (i.e. above threshold).
  
  # Define contrasts of interest
  relevant_indicators <- data.frame(index = c(1,2,4,5), DIY.indicator = c("glucose_only","PHB_only","glu_and_PHB","glu_and_water"))
  
  # Filter out only combinations of interest
  results <- results[which(results$index %in% relevant_indicators$index),]
  
  # Clarify what indices correspond to
  results$OTU <- rownames(results)
  results <- merge(results, relevant_indicators, by = "index")
  results$index <- NULL
  
  # Order by Indicator Value
  results <- results[rev(order(results$stat)),]
  
  # Add in Factor Info
  results$factor <- factor
  results$tree <- ecoplot

  # Add in Taxonomy
  foo2 <- subset_taxa(foo, taxa_names(foo) %in% results$OTU)
  foo2 <- psmelt(foo2)
  tax <- unique(foo2[,c("OTU","Phylum","Class","Order","Family","Genus","Species")])
  
  # Add in Relative Abundance
  abund <- ddply(foo2, ~ OTU + tree, summarise, avg.RelAbund = mean(Abundance))
  abund <- merge(abund, tax, by = "OTU")

  if (count == 1){
    x <- merge(results, abund, by = "OTU")
    count = count + 1
  } else {
    x <- rbind(x, merge(results, abund, by = "OTU"))
  }
}

# order by relative abundance
x <- x[rev(order(x$avg.RelAbund)),]

# Filter out low abundance (these are inconsistent among ecoplots, and driven by stochasticity)
results <- subset(x, avg.RelAbund > 0.34)
write.csv(results, file = "indicator.analyses.abund.csv", quote = F, row.names = F)
table(results$DIY.indicator)


########################
## Indicator by Taxonomy

x <- psmelt(p)

# average
x <- ddply(x, ~ Genus + Sample + tree + treatment, summarise, total = sum(Abundance))

count = 1
for (ecoplot in c("Sugar Maple","Red Pine","Black Locust")){
  foo <- subset(x, tree == ecoplot)
  
  groups <- unique(foo[,c("Sample","treatment")])
  counts <- reshape(foo[,c("Sample","Genus","total")], direction = "wide", idvar = "Sample", timevar = "Genus")
  colnames(counts) <- gsub("total.","",colnames(counts))
  row.names(counts) <- counts$Sample
  counts$Sample <- NULL
  groups <- groups$treatment
  
  # Run Indicator Speices
  indval = multipatt(counts, groups, control = how(nperm=999))
  
  # Store Results in Table
  results <- indval$sign
  
  # Remove NA
  if(length(which(is.na(results$p.value)))>0){
    results <- results[-which(is.na(results$p.value)),]  
  }
  
  # Filter for signficance
  results <- results[which(results$p.value < 0.15),] # permutations of 999 gives p-value of 0.001, which is adjusted to 0.06 (i.e. above threshold).
  
  # Define contrasts of interest
  relevant_indicators <-data.frame(index = c(1,2,4,5), DIY.indicator = c("glucose_only","PHB_only","glu_and_PHB","glu_and_water"))
  
  # Filter out only combinations of interest
  results <- results[which(results$index %in% relevant_indicators$index),]
  
  # Clarify what indices correspond to
  results$OTU <- rownames(results)
  results <- merge(results, relevant_indicators, by = "index")
  results$index <- NULL
  
  # Order by Indicator Value
  results <- results[rev(order(results$stat)),]
  
  # Add in Factor Info
  results$factor <- factor
  results$tree <- ecoplot
  
  if (nrow(results) < 0){
    if (count == 1){
      final_result <- results
      count = count + 1
    } else {
      final_result <- rbind(final_result, results)
    }    
  }
}
```
