## Supplementary material for "Phenolic acid-degrading *Paraburkholderia* prime decomposition in forest soil": BLOCK02ii - in vitro - Wilcoxon tests.R

```
library(indicspecies)
library(phyloseq)
library(plyr)
library(reshape2)

## User Input
method <- "aggregate"  # By 'tree' or by 'aggregate'

# Import data
p <- readRDS(file = "raw_data/p_priming.final.filter.norm.rds")  

# Combine PHB and PHB+inhibitor treatments
sample_data(p)[which(sample_data(p)[,"treatment"] == "C13_PHB_inhibited"), "treatment"] <- "C13_PHB"

# Remove 'field samples' and cell amendments (uninformative for this analysis)
p <- subset_samples(p, treatment != "field" & treatment != "C13_PHB__C12_RP11")

# Remove cDNA samples (activity had subsided by day 7)
p <- subset_samples(p, amplicon == "DNA")

# Remove sequencing data from first pass 
p <- subset_samples(p, !(sample_names(p) %in% readRDS(file = "run1.sampleIDs.rds")))

# Convert to dataframe
x <- psmelt(p)

# Cycle through all genera looking for statistical differences

if (method == "aggregate"){
  x <- ddply(x, ~ Sample + treatment + Family, summarise, Total = sum(Abundance))
} else {
  x <- ddply(x, ~ Sample + tree + treatment + Family, summarise, Total = sum(Abundance))
}

count = 1
for (family in unique(x$Family)){
  foo <- subset(x, Family == family)
  
  if (method == "aggregate"){
    
    # multiple comparisons (Doesn't change anything if you use wilcoxon versus Tukey)
    stat <- pairwise.wilcox.test(foo$Total, foo$treatment, p.adj = "bonf")
    stat <- as.numeric(unlist(stat)[c(3,4,6)])
    keep <- which(stat < 0.05)
    
    #stat <- data.frame(TukeyHSD(aov(lm(Total ~ treatment, data = foo)))$treatment, stringsAsFactors = F)
    #keep <- which(stat$p.adj < 0.05)
    
    if (length(keep) > 0){
      if (count == 1){
        results <- data.frame(Taxon = rep(family, length(keep)), p.value = stat[keep], contrast = c("glu.vs.PHB","glu.vs.water","PHB.vs.water")[keep], stringsAsFactors = F)
        #results <- data.frame(Taxon = rep(family, length(keep)), p.value = stat$p.adj[keep], contrast = c("glu.vs.PHB","glu.vs.water","PHB.vs.water")[keep], stringsAsFactors = F)
        count = count + 1  
      } else {
        results <- rbind(results, data.frame(Taxon = rep(family, length(keep)), p.value = stat[keep], contrast = contrasts[keep], stringsAsFactors = F))
        #results <- rbind(results, data.frame(Taxon = rep(family, length(keep)), p.value = stat$p.adj[keep], contrast = c("glu.vs.PHB","glu.vs.water","PHB.vs.water")[keep], stringsAsFactors = F))
        
      }
    }    
  } else {
    for (species in unique(foo$tree)){
      foo2 <- subset(foo, tree == species)
      
      # multiple comparisons
      stat <- data.frame(TukeyHSD(aov(lm(Total ~ treatment, data = foo2)))$treatment, stringsAsFactors = F)
      keep <- which(stat$p.adj < 0.05)
      
      if (length(keep) > 0){
        if (count == 1){
          results <- data.frame(Taxon = rep(family, length(keep)), tree = species, p.value = stat$p.adj[keep], contrast = c("glu.vs.PHB","glu.vs.water","PHB.vs.water")[keep], stringsAsFactors = F)
          count = count + 1  
        } else {
          results <- rbind(results, data.frame(Taxon = rep(family, length(keep)), tree = species, p.value = stat$p.adj[keep], contrast = c("glu.vs.PHB","glu.vs.water","PHB.vs.water")[keep], stringsAsFactors = F))
        }
      }
    }
  }
}
```
