## Supplementary material for "Phenolic acid-degrading *Paraburkholderia* prime decomposition in forest soil": BLOCK02j - in vitro - Plot Any ASV or Taxon.R

```
library(phyloseq)
library(plyr)
library(ggplot2)


###########################
## USER INPUT: Select Taxon

method = "Taxa"   ## By Taxa or by ASV

if (method == "Taxa"){
  rank = "Genus"
  taxa = "Burkholderia-Caballeronia-Paraburkholderia" # "Burkholderia-Caballeronia-Paraburkholderia"
  ASV = "foo"
} else {
  ASV <- "5b18237cf385326c2cda6c3441ba8d40"
  # ASV <- names(taxa_sums(p)[rev(order(taxa_sums(p)))][1:20]) ## Top 20 most abundant
  #ASV <- results$OTU
}


##### Run script
##### 
p <- readRDS(file = "raw_data/p_priming.final.filter.norm.rds")  

# Remove sequencing data from first pass 
p <- subset_samples(p, !(sample_names(p) %in% readRDS(file = "run1.sampleIDs.rds")))

if (length(ASV) == 1 | method == "Taxa"){
  
  # Get Data and subset to taxon of interest
  if (method == "Taxa"){
    taxa_table <- as(tax_table(p)[,rank], "vector")
    KEEP <- rownames(tax_table(p))[which(taxa_table == taxa)]
    p_taxa <- prune_taxa(KEEP, p)
  } else {
    p_taxa <- prune_taxa(ASV, p)
  }
  
  # Prep dataframe
  plot_me <- data.frame(Counts = sample_sums(p_taxa))
  plot_me <- cbind(plot_me, sample_data(p_taxa)[,c("tree","treatment","amplicon")]) 

  # Set factor order  
  plot_me$tree <- factor(plot_me$tree, levels = c("Sugar Maple","Black Locust","Red Pine"))
  plot_me$amplicon <- factor(plot_me$amplicon, levels = c("cDNA","DNA"))
  plot_me$treatment <- factor(plot_me$treatment, levels = c("field","water_control","C13_glucose","C13_PHB","C13_PHB_inhibited","C13_PHB__C12_RP11"))
  
  if (method == "Taxa"){
    
    plot <- ggplot(plot_me, aes(x=treatment, y=Counts, colour = tree)) + geom_boxplot(outlier.shape = NA) + ylab("Average Counts per Thousand Reads") + ggtitle(taxa) + facet_grid(~ amplicon)
    plot <- plot + geom_point(position=position_jitterdodge(), alpha=0.5)
    print(plot)

    writeLines("Save plots?")
    switch(menu(c("Discard","Continue","Exit")), step1 <- "Discard", step1 <- "Save to File", step1 <- "Exit")
    
    if (step1 == "Exit"){
      break
    } else if (step1 == "Save to File"){
      ggsave(plot, filename=paste("raw_data/",rank,taxa,'priming.plot.pdf',sep="."), height=16, width=16)
    }
  } else {
    
    plot <- ggplot(plot_me, aes(x=treatment, y=Counts, colour = tree)) + geom_boxplot(outlier.shape = NA) + ylab("Average Counts per Thousand Reads") + ggtitle(paste(tax_table(p_taxa),collapse=";"))  + facet_grid(~ amplicon)
    plot <- plot + geom_point(position=position_jitterdodge(), alpha=0.5)
    print(plot)
    print(ASV)
    
    writeLines("Save plots?")
    switch(menu(c("Discard","Continue","Exit")), step1 <- "Discard", step1 <- "Save to File", step1 <- "Exit")
    
    if (step1 == "Exit"){
      break
    } else if (step1 == "Save to File"){
      ggsave(plot, filename=paste("raw_data/",ASV,'priming.plot.pdf',sep="."), height=16, width=16)
    }
  }
} else {
  # Get Data
  p_taxa <- prune_taxa(ASV, p)
  
  for (n in ASV){
    foo <- prune_taxa(n, p_taxa)
    
    # Prep dataframe
    plot_me <- data.frame(Counts = sample_sums(foo))
    plot_me <- cbind(plot_me, sample_data(foo)[,c("tree","treatment","amplicon")]) 
    
    # Set factor order  
    plot_me$tree <- factor(plot_me$tree, levels = c("Sugar Maple","Black Locust","Red Pine"))
    plot_me$amplicon <- factor(plot_me$amplicon, levels = c("cDNA","DNA"))
    plot_me$treatment <- factor(plot_me$treatment, levels = c("field","water_control","C13_glucose","C13_PHB","C13_PHB_inhibited","C13_PHB__C12_RP11"))

    plot <- ggplot(plot_me, aes(x=treatment, y=Counts, colour = tree)) + geom_boxplot(outlier.shape = NA) + ylab("Average Counts per Thousand Reads") + ggtitle(paste(n,paste(tax_table(foo), collapse=";"))) + facet_grid(~ amplicon)
    plot <- plot + geom_point(position=position_jitterdodge(), alpha=0.5)
    print(n)
    print(plot)
    
    writeLines("Save plots?")
    switch(menu(c("Discard","Continue","Exit")), step1 <- "Discard", step1 <- "Save to File", step1 <- "Exit")
    
    if (step1 == "Exit"){
      break
    } else if (step1 == "Save to File"){
      ggsave(plot1, filename=paste("raw_data/",n,'priming.plot.pdf',sep="."), height=16, width=16)
    }
  }
}
```
