## Supplementary material for "Phenolic acid-degrading *Paraburkholderia* prime decomposition in forest soil": BLOCK02k - in vitro - Figure 2.R

```
library(phyloseq)
library(plyr)
library(ggplot2)


std <- function(x) { 
  NA_Count<-is.na(x)
  NA_Count<-length(NA_Count["TRUE"])
  sd(x, na.rm=T)/sqrt(length(x)-NA_Count)
}

limits <- aes(ymax = average + error, ymin=average - error)

# Import data
p <- readRDS(file = "raw_data/p_priming.final.filter.norm.rds")  
taxa_table <- as(tax_table(p)[,"Genus"], "vector")
KEEP <- rownames(tax_table(p))[which(taxa_table == "Burkholderia-Caballeronia-Paraburkholderia")]
p_taxa <- prune_taxa(KEEP, p)
x <- psmelt(p_taxa)
x$tree <- factor(x$tree, levels = c("Sugar Maple","Black Locust","Red Pine"))
x$amplicon <- factor(x$amplicon, levels = c("cDNA","DNA"))
x$treatment <- factor(x$treatment, levels = c("field","water_control","C13_glucose","C13_PHB","C13_PHB_inhibited","C13_PHB__C12_RP11"))


##################################
# Information on Field Populations

control = "field"  # "field" | "water_control"
x.control <- subset(x, treatment == control)

ddply(x.control, ~ tree + amplicon + OTU, summarise, total = mean(Abundance))
ddply(x.control, ~ amplicon + OTU, summarise, total = mean(Abundance))

# RP11 in field
foo <- subset(x.control, amplicon == "DNA" & OTU == "5b18237cf385326c2cda6c3441ba8d40")

# Average abundance of taxa in heavy fractions
foo <- psmelt(p_taxa)
foo <- subset(foo, fraction %in% c("3-4","5","6","7","8") & enrichment == "C13")
ddply(foo, ~ tree + soil.type, summarise, average = mean(Abundance))


############################
# Figure 2A - PHB responders
x.PHB <- subset(x, treatment %in% c("field","water_control","C13_glucose","C13_PHB","C13_PHB_inhibited"))
x.PHB <- subset(x, !(Sample %in% readRDS(file = "run1.sampleIDs.rds")))

# average "C13_PHB" & "C13_PHB_inhibited"
x.PHB$treatment <- gsub("C13_PHB_inhibited","C13_PHB",x.PHB$treatment)

# stats
for (tre in c("Sugar Maple", "Black Locust", "Red Pine")){
  print(tre)
  stats <- ddply(subset(x.PHB, tree == tre), ~ Sample + treatment + tree + amplicon, summarise, total = sum(Abundance))
  stats <- subset(stats, treatment != "field")
  fit <- aov(lm(total ~ treatment, data = stats))
  print(TukeyHSD(fit))
}

# plot
plot_me1 <- ddply(x.PHB, ~ treatment + OTU + tree, summarise, average = mean(Abundance))
plot_me1$treatment <- factor(plot_me1$treatment, levels = c("field","water_control","C13_glucose","C13_PHB"))

# order bars
plot_me1 <- plot_me1[rev(order(plot_me1$tree, plot_me1$average)),]
plot_me1$OTU <- factor(plot_me1$OTU, levels = ddply(plot_me1, ~ OTU, summarise, major = mean(average))[order(ddply(plot_me1, ~ OTU, summarise, major = mean(average))$major),"OTU"])

# plot
plot <- ggplot(plot_me1, aes(x=treatment, y=average, fill = OTU)) + geom_bar(position="stack", stat="identity") + ylab("Counts per Thousand Reads") + facet_grid(~tree)
plot
ggsave(plot, filename="Figure2.ASV.phylotypes.barplot.pdf", height=16, width=16)

# plot error bars
foo <- ddply(x.PHB, ~ treatment + Sample + tree, summarise, total = sum(Abundance))
error <- ddply(foo, ~ treatment + tree,  summarise, error = std(total))
plot_me2 <- ddply(foo, ~ treatment + tree, summarise, average = mean(total))
plot_me2 <- cbind(plot_me2, error = error$error)
plot_me2$treatment <- factor(plot_me2$treatment, levels = c("water_control","C13_glucose","C13_PHB"))

plot <- ggplot(plot_me2, aes(x=treatment, y=average)) + geom_bar(position="dodge", stat="identity") + ylab("Counts per Thousand Reads") + facet_grid(~tree)
plot <- plot + geom_errorbar(limits, position=position_dodge(0.9), width=0.25)  
plot
ggsave(plot, filename="Figure5.ASV.phylotypes.errorbars.pdf", height=16, width=16)


#############################
# Figure 2B - cDNA versus DNA
x.cDNA <- subset(x, treatment %in% c("C13_PHB","C13_PHB_inhibited"))

# average "C13_PHB" & "C13_PHB_inhibited"
x.cDNA$treatment <- gsub("C13_PHB_inhibited","C13_PHB",x.cDNA$treatment)
x.cDNA <- ddply(x.cDNA, ~ tree + amplicon + sample, summarise, total = sum(Abundance))
x.cDNA <- ddply(x.cDNA, ~ tree + amplicon, summarise, average = mean(total))

# plot
plot <- ggplot(x.cDNA, aes(x=amplicon, y=average)) + geom_bar(position="stack", stat="identity") + ylab("Counts per Thousand Reads") + facet_grid(~tree)
plot
ggsave(plot, filename="Figure2.ASV.cDNA.vs.DNA.barplot.pdf", height=16, width=16)


######################################
# Figure S5A - Cell-eating Streptomyces

taxa_table <- as(tax_table(p)[,"Family"], "vector")
KEEP <- rownames(tax_table(p))[which(taxa_table == "Streptomycetaceae")]
p_taxa <- prune_taxa(KEEP, p)
x <- psmelt(p_taxa)
x$tree <- factor(x$tree, levels = c("Sugar Maple","Black Locust","Red Pine"))
x$amplicon <- factor(x$amplicon, levels = c("cDNA","DNA"))
x$treatment <- factor(x$treatment, levels = c("field","water_control","C13_glucose","C13_PHB","C13_PHB_inhibited","C13_PHB__C12_RP11"))

x.necro <- subset(x, treatment %in%  c("field","water_control","C13_glucose","C13_PHB","C13_PHB_inhibited","C13_PHB__C12_RP11"))

# average "C13_PHB" & "C13_PHB_inhibited"
x.necro$treatment <- gsub("C13_PHB_inhibited","C13_PHB",x.necro$treatment)
x.necro <- ddply(x.necro, ~ treatment + OTU + tree + amplicon + Genus, summarise, average = mean(Abundance))
x.necro$treatment <- factor(x.necro$treatment, levels = c("field","water_control","C13_glucose","C13_PHB","C13_PHB__C12_RP11"))

# order bars
x.necro <- x.necro[rev(order(x.necro$tree, x.necro$amplicon, x.necro$average)),]
x.necro$OTU <- factor(x.necro$OTU, levels = ddply(x.necro, ~ OTU, summarise, major = mean(average))[order(ddply(x.necro, ~ OTU, summarise, major = mean(average))$major),"OTU"])

# plot
plot <- ggplot(x.necro, aes(x=treatment, y=average, fill = OTU)) + geom_bar(position="stack", stat="identity") + ylab("Counts per Thousand Reads") + facet_grid(~tree+amplicon)
plot

# repeat with genus
x.necro <- ddply(x.necro, ~ treatment + Genus + tree + amplicon, summarise, total = sum(average))

# plot
plot <- ggplot(x.necro, aes(x=treatment, y=total, fill = Genus)) + geom_bar(position="stack", stat="identity") + ylab("Counts per Thousand Reads") + facet_grid(~tree+amplicon)
plot

#############################
# Figure S5B - cDNA versus DNA
x.cDNA <- subset(x, treatment %in% c("C13_PHB__C12_RP11"))

# average "C13_PHB" & "C13_PHB_inhibited"
x.cDNA <- ddply(x.cDNA, ~ tree + amplicon + sample, summarise, total = sum(Abundance))
x.cDNA <- ddply(x.cDNA, ~ tree + amplicon, summarise, average = mean(total))

# plot
plot <- ggplot(x.cDNA, aes(x=amplicon, y=average)) + geom_bar(position="stack", stat="identity") + ylab("Counts per Thousand Reads") + facet_grid(~tree)
plot
```
