## Supplementary material for "Phenolic acid-degrading *Paraburkholderia* prime decomposition in forest soil": BLOCK02l - in vitro - Sirois et al. 2018 - Figure S8.R

```
library(phyloseq)
library(plyr)
library(ggplot2)
library(indicspecies)

normalize_to_cpt <- function(p){
  return(apply(otu_table(p), 2, function(x) x/(sum(x)/1000)))
}


#######################################
# Overlap with Sirois and Buckley, 2018 - a study focused on DNA-degrading bacterial populations

## Pre-process and filter data
##

# pre-process Sara's data to remove spike ASV
p <- readRDS(file = "raw_data/p_eDNA.final.rds")  

# Spike ASV: "296015136dead5777a07024affaded9a" # Streptococcus mutans htrA 
p <- subset_taxa(p, taxa_names(p) != "296015136dead5777a07024affaded9a")

# Remove empty sample table columns
keep <- unlist(lapply(sample_data(p), function(x) length(which(is.na(x))) == length(x)))
keep <- names(keep[which(keep != TRUE)])
sample_data(p) <- sample_data(p)[,keep]

# Keep information about the total # of sequences and OTUs prior to filtering for later
total_seqs <- sum(sample_sums(p))
total_otus <- length(taxa_names(p))

# Remove ASVs that could not be assigned at the domain level (except those with > 500 reads)
remove <- taxa_names(subset_taxa(p, Domain == "uncl_sequence" & taxa_sums(p) < 500))
p <- prune_taxa(setdiff(taxa_names(p), remove), p)

# Perform abundance filtering ('soft filtering') 
softfilter1 <- mean(sample_sums(p)) * 0.0001  # to 0.01% of total (i.e. 1 in 10000 across ALL samples)
softfilter2 <- mean(sample_sums(p)) * 0.01  # threshold of abundance to keep singletons and doubletons

# Hard Filter OTUs below 0.01% of each sample's Total Abundance (i.e. summed across all samples)
p <- prune_taxa(names(which(taxa_sums(p) > softfilter1)), p)

# Soft Filter OTUs which occur in less than three samples (with two exceptions)
present_absent<-otu_table(p) 
present_absent[present_absent > 0] <- 1

# 1) Remove all singletons except those with a max relative abundance greater than softfilter in any sample
REMOVE_singletons <- rownames(present_absent)[which(rowSums(present_absent) == 1)]
p_assess <- prune_taxa(REMOVE_singletons, p)
KEEP_singletons <- names(which(apply(otu_table(p_assess), 1, function(x) max(x) > softfilter2)))
REMOVE_singletons <- setdiff(REMOVE_singletons, KEEP_singletons)

# 2a) Remove all doubletons except those with a max relative abundance greater than softfilter in any sample OR same plot
REMOVE_doubletons <- rownames(present_absent)[which(rowSums(present_absent) == 2)]
p_assess <- prune_taxa(REMOVE_doubletons, p)

KEEP_doubletons <- names(which(apply(otu_table(p_assess), 1, function(x) max(x) > softfilter2)))
CHECK_doubletons <- prune_taxa(names(which(apply(otu_table(p_assess), 1, function(x) max(x) < softfilter2))), p_assess)

# 2b) Remove all doubletons except those with a max relative abundance greater than softfilter from the same treatment/soil type (etc.)
factor <- "LandType"

# Kept doubleton OTUs that co-occured in similar treatments
KEEP_paired_doubles <- vector()
for (taxa in taxa_names(CHECK_doubletons)){
  foo <- prune_taxa(taxa, CHECK_doubletons)
  foo <- prune_samples(names(which(sample_sums(foo) > 0)), p)
  
  if (nrow(unique(sample_data(foo)[,factor])) == 1) {
    KEEP_paired_doubles <- c(KEEP_paired_doubles, taxa)
  }
}

KEEP_doubletons <- c(KEEP_doubletons, KEEP_paired_doubles)

# 3) Filter all singletons and doubletons, then add back in those that passed criteria
KEEP <- rownames(present_absent)[which(rowSums(present_absent) >= 3)]
KEEP <- c(KEEP, KEEP_singletons, KEEP_doubletons)
p <- prune_taxa(KEEP, p)

# Remove Chloroplast & Mitochondria
p <- subset_taxa(p, Class != "Chloroplast" & Family != "mitochondria")

print(paste("The Percent of Sequences Remaining After Filtering is",round((sum(sample_sums(p))/total_seqs)*100, 2),sep=" "))
print(paste("The Number of OTUs was Reduced from", total_otus,"to", length(taxa_names(p)), "in the 'filtered' datasets (i.e.", round((length(taxa_names(p))/total_otus)*100,1),"% of OTUs remain)", sep=" "))

# Normalize Abundances - Scaled to Counts Per Thousand Reads
sample_totals <- sample_sums(p)
otu_table(p) <- otu_table(normalize_to_cpt(p), taxa_are_rows = TRUE)
saveRDS(p, file = "raw_data/p_eDNA.filtered.norm.rds")


###########################################
## Figure S8 - Plot changes in Streptomyces
p <- readRDS(file = "raw_data/p_eDNA.filtered.norm.rds")  

# Subset Sirois to Streptomycetaceae
p_taxa <- prune_taxa(taxa_names(subset_taxa(readRDS(file = "raw_data/p_eDNA.filtered.norm.rds"), Family == "Streptomycetaceae")), p)

# common ASVs between soil priming and Sirois and Buckley, 2018
common <- taxa_names(subset_taxa(readRDS(file = "raw_data/p_priming.final.filter.norm.rds"), taxa_names(readRDS(file = "raw_data/p_priming.final.filter.norm.rds")) %in% taxa_names(p_taxa)))

tax_table(subset_taxa(p, taxa_names(p) %in% common))

# filter low abundant 
#p_taxa <- subset_taxa(p_taxa, taxa_sums(p_taxa) > 20)

# prepare for plotting
x <- psmelt(p_taxa)
x <- x[,colSums(is.na(x))<nrow(x)]

x <- ddply(x, ~ day + LandType + moisture + temp + experiment + OTU + Genus, summarise, average = mean(Abundance))
x <- subset(x, experiment == "eDNA.habitat")
x$day <- factor(x$day, levels = c("0","7","21","35","49"))
x$Genus <- as.character(x$Genus)
x <- subset(x, Genus != "E1B-B2-114")

# denote ASVs in soil priming experiment
x$Genus[which(x$OTU %in% common)] <- "in.priming"

plot <- ggplot(x, aes(x=day, y=average, fill = OTU)) + geom_bar(position="stack", stat="identity") + ylab("Counts per Thousand Reads") + facet_grid(~LandType)
plot

# set factor order
x$Genus <- factor(x$Genus, levels = rev(c("Streptomyces","uncl_Streptomycetaceae","in.priming","Kitasatospora")))

x <- ddply(x, ~ day + LandType + Genus, summarise, total = sum(average))
plot <- ggplot(x, aes(x=day, y=total, fill = Genus)) + geom_bar(position="stack", stat="identity") + ylab("Counts per Thousand Reads") + facet_grid(~LandType)
plot
```
