## Supplementary material for "Phenolic acid-degrading *Paraburkholderia* prime decomposition in forest soil": BLOCK03a - pobA global - Curating IMG metagenome data.R

```
# Import Data
x <- read.csv(file = "raw_data/IMG.pobA.metagenomes.jan-2019.csv", stringsAsFactors = F, header = T)
x$Status <- NULL
meta <- read.csv(file = "raw_data/IMG.pobA.metagenomes.sample.info.csv", stringsAsFactors = F, header = T)
colnames(meta)[which(colnames(meta) == "Genome.Name...Sample.Name")] <- "Genome.Name"
colnames(meta)[which(colnames(meta) == "Gene.Count.....assembled")] <- "Total.Genes"
colnames(meta)[which(colnames(meta) == "Genome.Size.....assembled")] <- "Genome.Size"

x <- merge(x, meta, all.y= T, by = "Genome.Name")
x$Gene.Count[is.na(x$Gene.Count)] <- 0

# Clean-up
x[,c("Sequencing.Center","Sequencing.Status","Domain")] <- list(NULL)

# Order
x <- x[rev(order(x$Gene.Count)),]

# Fix crappy metadata
x$Habitat.original <- x$Habitat
x$Habitat <- gsub("soil","Soil",x$Habitat)
x$Habitat <- gsub("Prmafrost Soil","Permafrost",x$Habitat)
x$Habitat <- gsub("Permafrost Soil","Permafrost",x$Habitat)
x$Habitat <- gsub("unplanted Soil","Soil",x$Habitat)
x$Habitat <- gsub("Terrestrial Soil","Soil",x$Habitat)
x$Habitat <- gsub("Tropical rainforest Soil ","Tropical Rainforest Soil",x$Habitat)
x$Habitat <- gsub("Terrestrial","Soil",x$Habitat)
x$Habitat <- gsub("bulk Soil","Soil",x$Habitat)
x$Habitat <- unlist(lapply(x$Habitat, function(x) toupper(x)))
x$Habitat <- trimws(x$Habitat)
x$Habitat[grep("RHIZOSPHERE",x$Habitat)] <- "RHIZOSPHERE"
x$Habitat[grep("FERROUS",x$Habitat)] <- "FERROUS MICROBIAL MAT"
x$Habitat[grep("HOTSPRINGS",x$Habitat)] <- "HOT SPRING"
x$Habitat[grep("HOT SPRINGS",x$Habitat)] <- "HOT SPRING"
x$Habitat[grep("FEC",x$Habitat)] <- "FECES"
x$Habitat[grep("GUT|CECUM",x$Habitat)] <- "GUT"
x$Habitat[grep("RUMEN",x$Habitat)] <- "RUMEN"
x$Habitat[grep("ACID MINE",x$Habitat)] <- "ACID MINE DRAINAGE"
x$Habitat[grep("REACTOR",x$Habitat)] <- "BIOREACTOR"
x$Habitat[grep("GROUNDWATER",x$Habitat)] <- "GROUNDWATER"
x$Habitat[grep("PHYLLOSPHERE",x$Habitat)] <- "PHYLLOSPHERE"
x$Habitat[grep("ROOT",x$Habitat)] <- "ROOT"
x$Habitat[grep("LITTER",x$Habitat)] <- "LITTER"
x$Habitat[grep("BIOFILM",x$Habitat)] <- "BIOFILM"
x$Habitat[grep("CITY",x$Habitat)] <- "URBAN"
x$Habitat[grep("LAKE",x$Habitat)] <- "LAKE"
x$Habitat[grep("MARINE|OCEAN",x$Habitat)] <- "MARINE"
x$Habitat[grep("ENDO",x$Habitat)] <- "ENDOSPHERE"
x$Habitat[grep("HUMAN",x$Habitat)] <- "HUMAN"
x$Habitat[grep("FOOD WASTE|COMPOST",x$Habitat)] <- "COMPOST"
x$Habitat[grep("SEW|WASTE|WASTEWATER TREATMENT|SLUDGE",x$Habitat)] <- "SEWAGE"
x$Habitat[grep("PEAT",x$Habitat)] <- "PEATLAND"
x$Habitat[grep("TROPICAL FOREST SOIL",x$Habitat)] <- "FOREST SOIL"
x$Habitat[grep("BOG FOREST SOIL",x$Habitat)] <- "FOREST SOIL"

# Fix weird duplicated assemblies
weirdos <- x[which(duplicated(x$Gene.Count))[which(which(duplicated(x$Gene.Count)) %in% which(duplicated(x$Genome.Name)))],]
weirdos <- unique(weirdos[,c("Gene.Count","Genome.Name")])

# Cycle through weird duplicates (these are all assembled data, but some projects were re-analyzed. It is unclear which number is correct, so go with the conservative higher assembly count)
for (n in 1:nrow(weirdos)){
  foo <- subset(x, Gene.Count == weirdos[n,1] & Genome.Name == weirdos[n,2])
  foo <- subset(foo, Genome.Size == max(foo$Genome.Size))  

  x <- rbind(subset(x, !(Gene.Count == weirdos[n,1] & Genome.Name == weirdos[n,2])),foo)  
}

# Filter Small and Empty genomes
x <- subset(x, Genome.Size > 1000000)

# Normalize
x.norm <- subset(x, x$Gene.Count != 0)
x.norm$cpm <- x.norm$Gene.Count/(x.norm$Genome.Size/1000000)  # Normalize to total size of assembly
foo <- subset(x, x$Gene.Count == 0)
foo$cpm <- 0
x <- rbind(x.norm, foo)
x <- x[rev(order(x$cpm)),]

# Save
saveRDS(x, file = "raw_data/IMG.pobA.metagenome.counts.final.rds")
```
