## Supplementary material for "Phenolic acid-degrading *Paraburkholderia* prime decomposition in forest soil": BLOCK03b - pobA global - Plotting IMG metagenome data.R

```
library(plyr)
library(ggplot2)
library(Hmisc)
library(reshape2)

## Figure 5B

# Import data
x <- readRDS(file = "raw_data/IMG.pobA.metagenome.counts.final.rds")

# remove low abundance sample types
x <- subset(x, cpm > 1)
x <- subset(x, Habitat %in% names(which(table(x$Habitat) > 50)))
x <- subset(x, Ecosystem.Category %in% names(which(table(x$Ecosystem.Category) > 50)))

# Order
x <- x[rev(order(x$cpm)),]
x$Habitat <- factor(x$Habitat, levels = unique(x$Habitat))

# Plot by Ecosystem and Habitat
plot <- ggplot(x, aes(x=Habitat, y=cpm)) + geom_jitter(width = 0.15, alpha = 0.5)
plot + theme(axis.text.x = element_text(angle = 90, hjust = 1))
plot

# Plot Boxplot
plot <- ggplot(x, aes(x=Habitat, y=cpm)) + geom_boxplot(outlier.shape = NA)
ggsave(plot, filename = "pobA.environmental.distribution.boxes.pdf", height=7.5, width=6)

# stats
wilcox.test(subset(x, Habitat == "FOREST SOIL")$cpm, subset(x, Habitat == "SOIL")$cpm)
wilcox.test(subset(x, Habitat == "MARINE")$cpm, subset(x, Habitat == "FRESHWATER")$cpm)
```
