## Supplementary material for "Phenolic acid-degrading *Paraburkholderia* prime decomposition in forest soil": BLOCK03c - pobA global - Plotting IMG genome data.R

```
library(plyr)
library(ggplot2)

## Figure 5A

# Import
x <- read.csv(file = "raw_data/IMG.pobA.genomes.all.csv", stringsAsFactors = F, header = T)
x <- unique(x) # some redundancy in IMG
colnames(x)[3] <- "genomeID"

# Add in taxonomy
totals <- read.csv(file = "raw_data/IMG.pobA.genome.taxonomy.csv", stringsAsFactors = F, header = T)
totals$imgID <- NULL
totals <- unique(totals)
assign_me <- subset(x, is.na(x$Phylum))
assign_me$Phylum <- NULL
foo <- merge(assign_me, totals, by = "genomeID")
foo[which(foo$Phylum == "Proteobacteria"), "Phylum"] <- "Epsilonproteobacteria"
x <- rbind(subset(x, !(is.na(x$Phylum))),foo)

# Sum
x$Count <- 1
counts <- ddply(x, ~ Phylum, summarise, present = sum(Count))

# Normalize to total genomes of each phylum in IMG
totals <- read.csv(file = "raw_data/IMG.total.genomes.csv", stringsAsFactors = F, header = T)
counts <- merge(counts, totals, by = "Phylum")

counts$prop.prez <- counts$present/counts$total.genomes
counts$prop.abs <- (counts$total.genomes-counts$present)/counts$total.genomes

# Plot
plot_me <- data.frame(Taxa = rep(counts$Phylum, 2), measure = c(rep("present",15),rep("absent",15)), proportion = c(counts$prop.prez, counts$prop.abs), stringsAsFactors = F)

# order by percentage
counts <- counts[rev(order(counts$prop.prez)),]
plot_me$Taxa <- factor(plot_me$Taxa, levels = counts$Phylum)

# make stacked bar plots
plot <- ggplot(plot_me, aes(y = proportion, x = Taxa, fill = measure)) + geom_bar(stat="identity", position = "fill")
plot

########################################
# Which Genera most commonly encode pobA
x <- read.csv(file = "raw_data/IMG.pobA.genomes.all.csv", stringsAsFactors = F, header = T)
x <- unique(x) # some redundancy in IMG
colnames(x)[3] <- "genomeID"
x$Phylum <- NULL

taxonomy <- read.csv(file = "raw_data/IMG.ranks.genomes.csv", stringsAsFactors = F, header = T)
x <- merge(x, taxonomy, by = "genomeID")

# Genus with greatest frequency of pobA
totals <- read.csv(file = "raw_data/IMG.total.genomes.by.genus", stringsAsFactors = F, header = T)
colnames(totals)[1] <- "Genus"
foo <- x
foo$Count <- 1
foo <- ddply(foo, ~ Genus, summarise, present = sum(Count))
foo <- merge(foo, totals, by = "Genus")
foo$percent <- foo$present/foo$Genomes.Count
foo <- subset(foo, Genomes.Count > 10)  # filter low abundant genera (those with 1 or 2 genomes skew the 'percent')
foo <- foo[rev(order(foo$percent)),]
foo$rank <- 1:nrow(foo)
subset(foo, Genus %in% c("Pseudomonas", "Burkholderia", "Paraburkholderia", "Streptomyces","Phenylobacterium","Novosphingobium","Conexibacter","Collimonas"))

# Plot pie charts for genera
toi <- c("Alphaproteobacteria","Betaproteobacteria","Gammaproteobacteria","Actinobacteria")

for (taxa in toi){
  
  # Subset
  foo <- subset(x, Class == taxa)
  
  # Sum
  foo$Count <- 1
  counts <- ddply(foo, ~ Family, summarise, present = sum(Count))
  counts$percentage <- counts$present/nrow(subset(x, Class == taxa))

  # Bunch all counts less than 1%
  counts[which(counts$Family == "unclassified"), "Family"] <- "Other"
  counts[which(counts$percentage < 0.03), "Family"] <- "Other"
  counts <- ddply(counts, ~ Family, summarise, total.percent = sum(percentage))
  counts <- counts[rev(order(counts$total.percent)),] 
  
  pdf(paste(taxa,".piechart.pobA.JGI.genomes.pdf",sep="")) 
  pie(counts$total.percent, labels = counts$Family)
  dev.off() 
  
}
```
