## Supplementary material for "Phenolic acid-degrading *Paraburkholderia* prime decomposition in forest soil": BLOCK04a - pobA paralogs - Survey of genomes in IMG.R

```
#################################
# pobA duplication in IMG/ER Data
x <- readRDS(file = "IMG.pobA.genome.counts.with.tax.rds")

# Overview
x.table <- table(x$Genus, x$Gene.Count)

# filter genera with few genome reps
x.table <- x.table[which(rowSums(x.table) > 10),]

#Percent single-copy, duplicated or more
foo <- data.frame(Genus = names(x.table[,1]), single_copy = x.table[,1]/rowSums(x.table))
foo <- merge(foo, data.frame(Genus = names(x.table[,1]), duplicate = x.table[,2]/rowSums(x.table)), by = "Genus")
foo <- merge(foo, data.frame(Genus = names(x.table[,1]), triplicate = x.table[,3]/rowSums(x.table)), by = "Genus")
foo <- merge(foo, data.frame(Genus = names(x.table[,1]), quadruplicate = x.table[,4]/rowSums(x.table)), by = "Genus")
foo <- merge(foo, data.frame(Genus = names(x.table[,1]), quintuplate = x.table[,5]/rowSums(x.table)), by = "Genus")

# order by duplicate frequency
foo <- foo[rev(order(foo$duplicate)),]
foo$rank <- 1:nrow(foo)
head(foo)
subset(foo, Genus %in% c("Pseudomonas", "Burkholderia", "Paraburkholderia", "Streptomyces","Phenylobacterium","Novosphingobium","Conexibacter","Collimonas"))

# order by triplicate frequency
foo <- foo[rev(order(foo$triplicate)),]
head(foo)
subset(foo, Genus %in% c("Pseudomonas", "Burkholderia", "Paraburkholderia", "Streptomyces","Phenylobacterium","Novosphingobium","Conexibacter","Collimonas"))

# which freaks have four?
foo <- foo[rev(order(foo$quadruplicate)),]
head(foo)
subset(foo, Genus %in% c("Pseudomonas", "Burkholderia", "Paraburkholderia", "Streptomyces","Phenylobacterium","Novosphingobium","Conexibacter","Collimonas"))

# which freaks have five?
foo <- foo[rev(order(foo$quintuplate)),]
head(foo)
subset(foo, Genus %in% c("Pseudomonas", "Burkholderia", "Paraburkholderia", "Streptomyces","Phenylobacterium","Novosphingobium","Conexibacter","Collimonas"))

# in Burkholderia
burks <- subset(x, Family == "Burkholderiaceae")
subset(x, Family == "Burkholderiaceae" & Genus == "unclassified")
table(burks$Genus, burks$Gene.Count)

alphs <-subset(x, Class == "Alphaproteobacteria")
table(alphs$Genus, alphs$Gene.Count)

# in Actinos
actino <- subset(x, Phylum == "Actinobacteria")
colSums(table(actino$Genus, actino$Gene.Count))/sum(colSums(table(actino$Genus, actino$Gene.Count)))

foo <- data.frame(Genus = names(x.table[,1]), single_copy = x.table[,1]/rowSums(x.table))
foo <- merge(foo, data.frame(Genus = names(x.table[,1]), duplicate = x.table[,2]/rowSums(x.table)), by = "Genus")
foo <- merge(foo, data.frame(Genus = names(x.table[,1]), triplicate = x.table[,3]/rowSums(x.table)), by = "Genus")
foo <- merge(foo, data.frame(Genus = names(x.table[,1]), quadruplicate = x.table[,4]/rowSums(x.table)), by = "Genus")
foo <- merge(foo, data.frame(Genus = names(x.table[,1]), quintuplate = x.table[,5]/rowSums(x.table)), by = "Genus")
```
