## Supplementary material for "Phenolic acid-degrading *Paraburkholderia* prime decomposition in forest soil": BLOCK04b - pobA paralogs - MLSA with paralogs.R

```
library(ape)
library(plyr)
library(ggtree)
library(ggplot2)

# Import pobA annotations
x <- readRDS(file = "raw_data/pobA.hmm.output.rds")

### summarise number of pobA per genome
# get genomeID
genomeID <- unlist(lapply(as.vector(x$gene), function(x) unlist(strsplit(x, ";"))[2]))
genomeID <- unlist(lapply(genomeID, function(x) paste(unlist(strsplit(x, "_"))[1:2], collapse="_")))
x$genomeID <- genomeID

# add taxonomy
tax <- readRDS(file = "raw_data/MLSA.genomes.tax.rds") 
x <- merge(x, tax, by = "genomeID")

# summarise
x.sum <- x
x.sum$count <- 1
x.sum <- ddply(x.sum, ~ genomeID + Genus + Species, summarise, total.pobA = sum(count))

# Identify duplicates
dups <- subset(x, genomeID %in% subset(x.sum, total.pobA >= 2)$genomeID)

### import MLSA tree and integrade pobA information
x.tree <- read.tree(file = "raw_data/MLSA.RAxML.tree.nwk")
root <- "GCF_000007565.2"

# consolidate tree with genomes with pobA information (while keeping the root)
x.tree <- drop.tip(x.tree, setdiff(x.tree$tip.label, c(x$genomeID,"GCF_000007565.2")))

# root to Pseudomonas (outgroup)
x.tree <- root(x.tree, root)

### re-introduce tips that were removed by RAxML, given they were identical across the full 13,366 positions (n = 2071)
removed <- read.table(file = "raw_data/MLSA.identical.sequences.tsv", sep = "\t", stringsAsFactors = F, header = T)

# Add back in tips to each representative sequence
for (tip in intersect(removed$sequence1, x.tree$tip.label)){
  for (t in subset(removed, sequence1 == tip)$sequence2){
    x.tree <- bind.tip(x.tree, t, edge.length=0, where=which(x.tree$tip.label==tip), position=0)
  }
}

# final consolidate tree with pobA information
x.tree <- drop.tip(x.tree, setdiff(x.tree$tip.label, c(x$genomeID, root)))
saveRDS(x.tree, file = "raw_data/MLSA.pobA.duplication.final.rds") # For BLOCK04c

# Subset tree
x <- subset(x, genomeID %in% x.tree$tip.label)

### Mark tips with duplicated pobA
x.info <- unique(x[,c("genomeID","Genus")])

# add back in root
x.info <- rbind(x.info, data.frame(genomeID = "GCF_000007565.2", Genus = "Pseudomonas"))

# mark dups
x.info$dup <- 0
x.info$dup[which(x.info$genomeID %in% dups$genomeID)] <- 1
x.info$dup <- as.factor(x.info$dup)
colnames(x.info)[1] <- "label"

# add in isolation source information scraped from BioSample database
x.source <- read.csv(file = "raw_data/MLSA.genome.isolate.source.csv", header = T, stringsAsFactors = F)

# find where 'host' designation is present and isolation source is absent
host <- subset(x.source, ncbi.designator == "host")
host <- subset(host, !(is.na(host$description)))
source <- subset(x.source, ncbi.designator == "isolation source")
source <- subset(source, !(is.na(source$description)))

missing <- subset(host, genomeID %in% setdiff(host$genomeID, source$genomeID))
missing <- subset(missing, description != "environmental")
missing$description <- "host"
missing$ncbi.designator <- "isolation source"

# add back in new info
source <- rbind(source, missing)

# manually curate descriptors
source$description <- gsub(","," ",source$description)
new.descriptions <- read.csv(file = "raw_data/MLSA.curated.genome.isolate.source.csv", header = T, stringsAsFactors = F)
source <- merge(source, new.descriptions, by = "description")
colnames(source)[2] <- "label"

# remove stragglers
source <- subset(source, !(label %in% setdiff(source$label, x.info$label)))

# add to main dataframe
x.info <- merge(x.info, source[,c("label","new.descriptor")], by = "label", all.x = T)
saveRDS(x.info, file = "raw_data/isolate.source.info.rds") # For BLOCK04c

# only color descriptions for duplicated genomes
x.info$new.descriptor[which(x.info$dup == 0)] <- NA
x.info$RP11 <- 0
x.info$RP11[grep("GCF_900116445.1", x.info$label)] <- 1
x.info$RP11 <- as.factor(x.info$RP11)

# prep tree file for ploting
x.foo <- as_tibble(x.info)
x.tree <- as_tibble(x.tree)
x.tree <- full_join(x.tree, x.foo, by = "label")
x.tree <- as.treedata(x.tree)

# Plot Taxonomy and clade
plot <- ggtree(x.tree, branch.length='none', layout='circular') + theme(legend.position="right")
plot + geom_tippoint(aes(color=dup), alpha=0.25) + geom_treescale()
```
