## Supplementary material for "Phenolic acid-degrading *Paraburkholderia* prime decomposition in forest soil": BLOCK04c - pobA paralogs - Isolate source and paralogs.R

```
library(ape)
library(geiger)
library(phytools)
library(ggtree)

x.tree <- readRDS(file = "raw_data/MLSA.pobA.duplication.final.rds")
x.info <- readRDS(file = "raw_data/isolate.source.info.rds") 

# Plot whole tree
plot.phylo(x.tree, type = "u", show.tip.label = FALSE, cex = 10)

# Ralstonia
ralstonia <- data.frame(genomeID = tips(x.tree, "6857"), clade = "ralstonia", stringsAsFactors = F)

# Cupriavidus
cupria <- data.frame(genomeID = tips(x.tree, "6781"), clade = "cupriavidus", stringsAsFactors = F)

# Pandoraea
pando <- data.frame(genomeID = tips(x.tree, "6737"), clade = "pandoraea", stringsAsFactors = F)

# Paraburkholderia
para <- data.frame(genomeID = tips(x.tree, "6597"), clade = "paraburkholderia", stringsAsFactors = F)
para <- rbind(para, data.frame(genomeID = tips(x.tree, "6562"), clade = "paraburkholderia", stringsAsFactors = F))
para <- rbind(para, data.frame(genomeID = tips(x.tree, "6518"), clade = "paraburkholderia", stringsAsFactors = F))

# Caballeronia
caba <- data.frame(genomeID = tips(x.tree, "6672"), clade = "caballeronia", stringsAsFactors = F)

# Burkholderia clades
burk1 <- data.frame(genomeID = tips(x.tree, "5200"), clade = "burkholderia_clade1", stringsAsFactors = F)
burk2 <- data.frame(genomeID = tips(x.tree, "5616"), clade = "burkholderia_clade2", stringsAsFactors = F)
burk3 <- data.frame(genomeID = tips(x.tree, "5830"), clade = "burkholderia_clade3", stringsAsFactors = F)
burk4 <- data.frame(genomeID = tips(x.tree, "3512"), clade = "burkholderia_clade4", stringsAsFactors = F)
burk3.3 <- data.frame(genomeID = tips(x.tree, "6149"), clade = "burkholderia_clade3.3", stringsAsFactors = F)
burk3.2 <- data.frame(genomeID = setdiff(tips(x.tree, "5980"), tips(x.tree, "6149")), clade = "burkholderia_clade3.2", stringsAsFactors = F)
burk3.1 <- data.frame(genomeID = setdiff(tips(x.tree, "5830"), c(burk3.3$genomeID, burk3.2$genomeID)), clade = "burkholderia_clade3.1", stringsAsFactors = F)

# Plot pie-charts
for (clade in c("burk2","para","burk1","burk3.1","burk3.2","burk3.3","cupria")){
  foo <- get(clade)
  
  labels <- names(table(subset(x.info, label %in% foo$genomeID & dup == 1)$new.descriptor))
  values <- as.numeric(table(subset(x.info, label %in% foo$genomeID & dup == 1)$new.descriptor))
  
  pdf(paste("pie.duplicated", clade,"pdf",sep=".")) 
  pie(values, labels, main = paste("Duplicated in ", clade," (n=",sum(values),")",sep=""))
  dev.off() 
  
  labels <- names(table(subset(x.info, label %in% foo$genomeID & dup == 0)$new.descriptor))
  values <- as.numeric(table(subset(x.info, label %in% foo$genomeID & dup == 0)$new.descriptor))
  pdf(paste("pie.non.duplicated", clade,"pdf",sep=".")) 
  pie(values, labels, main = paste("Non-duplicated in ", clade," (n=",sum(values),")",sep=""))
  dev.off() 
}

for (clade in c("burk4","ralstonia","caba","pando")){
  foo <- get(clade)

  labels <- names(table(subset(x.info, label %in% foo$genomeID)$new.descriptor))
  values <- as.numeric(table(subset(x.info, label %in% foo$genomeID)$new.descriptor))
  
  pdf(paste("pie.non.duplicated", clade,"pdf",sep=".")) 
  pie(values, labels, main = paste(clade," (n=",sum(values),")",sep=""))
  dev.off() 
}

### Fisher exact test

# Soil
foo <- subset(x.info, !(is.na(new.descriptor)))
test_me <- matrix(c(nrow(subset(foo, new.descriptor == "soil" & dup == 1)), nrow(subset(foo, new.descriptor == "soil" & dup == 0)), nrow(subset(foo, new.descriptor != "soil" & dup == 1)), nrow(subset(foo, new.descriptor != "soil" & dup == 0))), nrow = 2)
fisher.test(test_me)
```
