## Supplementary material for "Phenolic acid-degrading *Paraburkholderia* prime decomposition in forest soil": BLOCK04d - pobA paralogs - phylogenetic conservation and dispersion.R

```
library("caper")

# Prepare input for consenTRAIT

# Import other dioxygenases
x <- read.table(file = "raw_data/consenTRAIT.dioxygenases.out", stringsAsFactors = F, header = F)
x[,c("V2","V5","V10","V11","V12","V13","V14","V15","V18","V19","V20","V21","V22")] <- list(NULL)
colnames(x) <- c("gene","gene.length","hmm","length","evalue","score","bias","hit.start","hit.end")

# get genome ID
x$genomeID <- unlist(lapply(x$gene, function(x) unlist(strsplit(x, "\\|"))[2]))
x$genomeID <- unlist(lapply(x$genomeID, function(x) paste(unlist(strsplit(x, "_"))[1:2], collapse="_")))

# Filter short hits
x$hit.length <- x$hit.end-x$hit.start
x$percent.length <- x$hit.length/x$length
x <- subset(x, percent.length > 0.9)

# Filter low scoring hits
x <- subset(x, score > 200)

# reshape and add to consenTRAIT table
x$count <- 1
x <- reshape(x[,c("genomeID","hmm","count")], direction = "wide", idvar = "genomeID", timevar = "hmm")
colnames(x) <- gsub("count.","",colnames(x))
x[is.na(x)] <- 0

# count genomes with both subunits
x$benzoate <- 0
x$benzoate[which(x$benzo_1_2_benA == 1 & x$benzo_1_2_benB == 1)] <- 1

x$protocatechuate <- 0
x$protocatechuate[which(x$protocat_alph == 1 & x$protocat_beta == 1)] <- 1
x[,c("benzo_1_2_benA","benzo_1_2_benB","protocat_alph","protocat_beta")] <- list(NULL)

colnames(x) <- c("V1","V3","V4","V5","V6")

colnames(consenTRAIT.info) <- c("V1","V2")
consenTRAIT.info <- merge(consenTRAIT.info, x, all.x = T, by = "V1")
consenTRAIT.info[is.na(consenTRAIT.info)] <- 0

# add informatino about which genomes encoded pobA
pobA <- readRDS(file = "all.pobA.from.hmms.for.consenTRAIT.rds")

consenTRAIT.info[grep("GCF_900230185.1", consenTRAIT.info$V1),]
consenTRAIT.info$V7 <- 0
consenTRAIT.info$V7[which(consenTRAIT.info$V1 %in% unique(pobA$genomeID))] <- 1

# V2 : pobA duplication  (n=391)
# V3 : catechol (n=3386)
# V4 : gentisate (n=1039)
# V5 : benzoate (n=3295)
# V6 : protocatechuate (n=3494)
# V7 : pobA (n=3380)

write.table(consenTRAIT.info, file = "consenTRAIT.trait.info.tsv", sep = "\t", col.names = F, row.names = F, quote=F)

#consenTRAIT.r -p 10 consenTRAIT.MLSA.tree.nwk GCF_000007565.2 consenTRAIT.trait.info.tsv


####################################
# Summarize results from consenTRAIT

x <- read.table(file = "Tau_D.txt", sep="\t", header=T, stringsAsFactors = F)
x$tree <- NULL
trait_list <- c("pobA_dup","catechol","gentisate","benzoate","protocatechuate","pobA")
x <- data.frame(trait = trait_list, tauD = colMeans(x))
dist <- cophenetic.phylo(readRDS(file = "pobA.duplication.MLSA.final.rds"))
x <- rbind(x, data.frame(trait = "whole tree", tauD = max(dist)))
write.csv(x, file = "consenTRAIT.tauD.table.csv", row.names = F, quote = F)

#########################################################
# Calculate Fritz and Purvis' phylogenetic dispersion (D)

x.info <- read.table(file = "consenTRAIT.trait.info.tsv", sep = "\t", stringsAsFactors = F, header = F)
colnames(x.info) <- c("genomeID","pobA_dup","catechol","gentisate","benzoate","protocatechuate","pobA")
x.tree <- read.tree(file = "RAxML_bipartitions.MLSA.burks.final.tree.cleaned.nwk")
x.tree <- root(x.tree, "GCF_000007565.2")

for (trait in trait_list){
  foo <- x.info[,c("genomeID",trait)]
  colnames(foo)[2] <- "trait"
  PD <- comparative.data(phy = x.tree, data = foo, names.col = genomeID, vcv = TRUE, na.omit = FALSE, warn.dropped = TRUE)
  PD$phy <- di2multi(PD$phy, tol = 0.00000001)   # 0.000005
  result = phylo.d(data=PD, binvar = trait, permut = 1000)
  print(trait)
  print(result)  
}

x.D <- data.frame(trait = c("pobA_dup","catechol","gentisate","benzoate","protocatechuate","pobA"), D = c("0.0979","-0.0317","0.0796","0.111","0.249","0.0365"), p_random = c("0","0","0","0","0","0"), p_brownian = c("0.109","0.57","0.119","0.14","0.041","0.356"))
write.csv(x.D, file = "phylogenetic.dispersion.table.csv", row.names = F, quote = F)
```
