## Supplementary material for "Phenolic acid-degrading *Paraburkholderia* prime decomposition in forest soil": BLOCK06 - Plot Any Taxa or OTU.R

```
library(phyloseq)
library(plyr)
library(reshape2)
library(ggplot2)
source("get.study.R")
source("get_geogroups.R")
source("fix.tillage.R")

## Specify Functions 
##

std <- function(x) { 
  NA_Count<-is.na(x)
  NA_Count<-length(NA_Count["TRUE"])
  sd(x, na.rm=T)/sqrt(length(x)-NA_Count)
}

limits <- aes(ymax = average + error, ymin=average - error)

########
########  USER INPUT
########

### Choose Studies
###
#study_list = c("liming","chazy","fullcyc","chazy.SIP","bioblitz")  ## "soil.health" | "liming" | "chazy" | "chazy.SIP" | "bioblitz" | "fullcyc"
study_list = c("chazy")  ## "soil.health" | "liming" | "chazy" | "chazy.SIP" | "bioblitz" | "fullcyc"

### Choose Method
###
method = "Taxa"   ## By Taxa or by OTU

### Choose Dataset
###
dataset = "normalized"  ## "normalized" | "css" | "filtered.rare"

### Choose Plot Style
###
plot.type = "dot"  ## "dot" | "bar"  # It is recommended to use the 'dot' version, particularly for SIP experiments (where error bars are inflated due to aggregation of time-course information)

### Choose Soil Health Variable to use Display
###
fact = "health.category" # "health.category", "tillage", "organic_matter.category", "pH.category", "ace_soil_protein_index.category", "active_carbon.category", "respiration.category", "surface_hardness.category", "subsurface_hardness.category", "aggregate_stability.category", "water_capacity.category"

# Display all options for variables
#head(sample_data(readRDS(file = "p_SSU.soil.health.filter.norm.rds")), 5)

### Select Taxa
###
rank = "Phylum"
taxa = "Actinobacteria"

#tax_table(p)[grep("Burkholderia", tax_table(p)[,"Genus"]),]
#tax_table(p)[grep("Nitrososphaera", tax_table(p)[,"Genus"]),]

#unique(tax_table(p_taxa)[,"Genus"])
#head(taxa_sums(p_taxa)[rev(order(taxa_sums(p_taxa)))])
#tax_table(prune_taxa("efdd7620bb28b2e6d806cf42d41cfe00",p_taxa))

# Caulobacter rhizosphaera : "43d5f162125cd7bf8ece35b3170c1042" (Caulobacter) # Cellulolytic | Caulobacter rhizosphaera | KCTC 52515 ($30 USD from Korean Culture Collection)
# Ellin329 (from GreenGenes) ~ "Rhizobiales_Incertae_Sedis" (Family) or Rhizomicrobium (Genus)
# Phenylobacterium: c("1abee12a36fcc29ba296ab598cf9a439","6fa5e4fe0449e0fe7e771ed0b1101c51","f34ff488e9d783a9bf687a20f21720b4","f8d3bd6d0f955e41d54c4d8f3f33f38b")

# DEEP-SIP
#taxa1 <- c("Devosia","Rhizobium","Ensifer","Agrobacterium","Sphingomonas","Sphingopyxis","Brevundimonas","Caulobacter","Asticcacaulis","Pseudoxanthomonas","Microbacterium","Lentzea","Kitasatospora","Streptomyces","Sporocytophaga","Sorangium","Nannocystis","Dactylosporangium","Catellatospora","Luteolibacter","Vampirovibrio","Pseudomonas","Xiphinematobacter","Rhodobacter","01D2Z36")
#taxa2 <- c("Microbacterium","Lentzea","Kitasatospora","Streptomyces","Sporocytophaga","Sorangium","Nannocystis","Dactylosporangium","Catellatospora","Luteolibacter","unclassified Vampirovibrionales","Pseudomonas","Candidatus_Xiphinematobacter","01D2Z36")
#taxa3 <- c("unclassified 01D2Z36","unclassified putative 01D2Z36")
#taxas <- c(taxa1, taxa2, taxa3)
#taxas <- unique(taxas)

# Help Finding Taxa Name
#tax_table(p)[grep("01D", tax_table(p)[,"Genus"]),]

#foo <- p_taxa
#foo <- prune_taxa(names(head(taxa_sums(foo)[rev(order(taxa_sums(foo)))])), foo)
#tax_table(foo)

### Or select OTU(s) 
###
#
OTU <- c("a7db197d3a76ed1d4fa13a99ff1cd999")  

#OTU <- readRDS(file = "DA101.ASVs.rds")
#OTU <- readRDS(file = "Armatimonadetes.ASVs.rds")

#OTU <- names(head(taxa_sums(p_taxa)[rev(order(taxa_sums(p_taxa)))]))
#OTU <- "46fb50c57230a2b1b88e41d382f10118"

#OTU <- unique(subset(readRDS(file = "all.indicators.buckleyDB.may.2018.rds"), Family == "Rhizobiaceae" & substrate == "cellulose")$OTU)
#OTUs <- c("2fea268498044452fdd476b1a5322e47","cb26644aaef39f96743f1f85614baef1","ab0d1fdbcfdea9360c49602d66386b1f")

### Additional Study Specific Options
###
only_heavy <- ""    # Set if you'd only like to compare abundance in the heavy fractions (1-10) does not factor if 'by_fraction' is specified
#show_geo_groups <- ""  # Set if you'd like to display a second plot with the geogroups labeled.
#remove_zeros <- ""  # This is only an option for dotplots and is only useful for time-course SIP studies, since the averages are downweighted for say, SIP-cellulose incubations, b/c all of the zeros in earlier timepoints.
#by_month <- ""     # Set if you'd like to breakdown chazy study by month
#by_day <- ""       # Set if you'd like to breakdown SIP studies by day
#by_fraction <- ""  # Set if you'd like to breakdown SIP studies by fraction

## Some Examples of Responders to Soil Health and in other studies
# Negative associated OTUs: 
#:"0073ff6a194e692f063aa9540e7d9334"
# Positively associated OTUs:
#: c("a7db197d3a76ed1d4fa13a99ff1cd999") # Pasture-associated organism


#p <- get_study("liming", "bacteria", "normalized")
#p <- subset_samples(p, treatment == "limed" & horizon == "Oe")
#p <- subset_taxa(p, Genus %in% c("Caulobacter","Rhizomicrobium","Rhizobium","Brevundimonas","Phenylobacterium","Devosia","Sphingomonas","Pseudoxanthomonas"))
#mean(sample_sums(p))

##
## MAIN
##

# Always append soil.health data to study_list
study_list <- append(study_list, "soil.health")

#plot <- ggplot(subset(plot_me, substrate == "cellulose"), aes(x=enrichment, y=Counts, colour = get(x_fill))) + geom_boxplot(outlier.shape = NA) + ylab("Average Counts per Thousand Reads") 
#plot <- plot + geom_point(position=position_jitterdodge(), alpha=0.5)
#plot

for (study in unique(study_list)){
    
    ### Import Data
    ###
    p <- get_study(study, "Bacteria", dataset)
    
    ### Set Factors of Interest for Each Individual Study
    ### 
    
    # List possible factors
    #colnames(sample_data(p)[,colSums(is.na(sample_data(p)))<nrow(sample_data(p))])
    
    if (study == "soil.health"){
      # Prepare factor levels  
      factors <- c(fact)
      
      if (fact == "tillage"){
        p <- fix_tillage(p)
      }
      
      # Prepare formula
      formula <- as.formula(paste("~ ",paste(factors, collapse="+"),sep=""))
      x_axis <- fact
      x_fill <- fact

      if (exists("facet")){
        rm(facet)  
      }
      
      # Clean-up for soil health data (breaks due to bioblitz code)
      if (exists("factor_levels")){
        rm(factor_levels)
      }
      
    } else if (study == "liming"){
      
      # Prepare factor levels  
      factors <- c("treatment","horizon","site","time")
      factor_levels <- list(c("control", "limed"),c("Oe", "Oa"),c("Honnedaga", "Woods"),c("pre", "post"))  # this will order your factors as you wish. Must be in same order as factors 
      
      # Prepare formula
      formula <- as.formula(paste("~ ",paste(factors, collapse="+"),sep=""))
      x_axis <- "treatment"
      x_fill <- "time"
      facet <- as.formula("~ site + horizon")
      
    } else if (study == "chazy"){
      
      ## Determine whether to use time
      
      if(exists("by_month")){
        # Prepare factor levels  
        factors <- c("treatment","time")
        factor_levels <- list(c("NTR", "NTH","PTR","PTH","PAS"),c("April", "June","July","September","October","November"))  # this will order your factors as you wish. Must be in same order as factors 
        facet <- as.formula("~ time")
        
        # clean-up
        rm(by_month)
        
      } else {
        # Prepare factor levels  
        factors <- c("treatment")
        
        # Temporary
        #sample_data(p)[which(sample_data(p)[,"treatment"] == "NTR"),"treatment"] <- "No-till"
        #sample_data(p)[which(sample_data(p)[,"treatment"] == "NTH"),"treatment"] <- "No-till"
        #sample_data(p)[which(sample_data(p)[,"treatment"] == "PTR"),"treatment"] <- "Till"
        #sample_data(p)[which(sample_data(p)[,"treatment"] == "PTH"),"treatment"] <- "Till"
        #sample_data(p)[which(sample_data(p)[,"treatment"] == "PAS"),"treatment"] <- "Pasture"
        #factor_levels <- list(c("No-till", "Till","Pasture"))
        
        factor_levels <- list(c("NTR", "NTH","PTR","PTH","PAS"))  # this will order your factors as you wish. Must be in same order as factors 
        
        if (exists("facet")){
          rm(facet)  
        }
      }
      
      # Prepare formula
      formula <- as.formula(paste("~ ",paste(factors, collapse="+"),sep=""))
      x_axis <- "treatment"
      x_fill <- "treatment"
      
    } else if (study == "chazy.SIP"){
      
      # Prepare factor levels  
      factors <- c("enrichment","fraction","substrate","day","treatment")
      substrates <- c("water","xylose","cellulose")
      days <- c(1,3,7,14,30)
      
      if (exists("by_day") & exists("by_fraction")){
        formula <- as.formula(paste("~ ",paste(c("enrichment","fraction","day","substrate","treatment"), collapse="+")))
        x_axis <- "fraction"
        x_fill <- "enrichment"
        facet <- as.formula("~ treatment + substrate")
        
        factors <- c("enrichment","fraction","substrate","day","treatment")
        factor_levels <- list(c("C12","C13","incubated"), seq(from= 1, 30, by=1), substrates, days, c("NTH","PTH"))
        
        rm(by_day)
        rm(by_fraction)
        
      } else if (exists("by_day")){
        
        # Only compare heavy fractions
        if (exists("only_heavy")){
          p <- subset_samples(p, fraction %in% c(1,2,3,4,5,6,7,8,9,10,11,12) | enrichment == "incubated")  
        }
        
        formula <- as.formula(paste("~ ",paste(c("enrichment","day","substrate","treatment"), collapse="+")))
        x_axis <- "day"
        x_fill <- "enrichment"
        facet <- as.formula("~ substrate + treatment")
        factor_levels <- list(c("C12","C13","incubated"), substrates, days, c("NTH","PTH"))
        factors <- c("enrichment","substrate","day","treatment")
        
        rm(by_day)
        
      } else if (exists("by_fraction")){
        formula <- as.formula(paste("~ ",paste(c("enrichment","fraction","substrate","treatment"), collapse="+")))
        x_axis <- "fraction"
        x_fill <- "enrichment"
        facet <- as.formula("~ substrate + treatment")
        
        factors <- c("enrichment","fraction","substrate","treatment")
        factor_levels <- list(c("C12","C13","incubated"), seq(from= 1, 30, by=1), substrates, c("NTH","PTH"))
        
        rm(by_fraction)
        
      } else {
        
        # Only compare heavy fractions
        if (exists("only_heavy")){
          p <- subset_samples(p, fraction %in% c(1,2,3,4,5,6,7,8,9,10,11,12) | enrichment == "incubated")  
        }
        
        formula <- as.formula(paste("~ ",paste(c("enrichment","substrate","treatment"), collapse="+")))
        x_axis <- "treatment"
        x_fill <- "enrichment"
        facet <- as.formula("~ substrate")
        factors <- c("enrichment","substrate","treatment")
        factor_levels <- list(c("C12","C13","incubated"), substrates, c("NTH","PTH"))
        
      }
      
    } else if (study == "bioblitz"){
      # Prepare factor levels  
      factors <- c("sample.type")
      factor_levels <- list(c("Soil","Rhizosphere","Phyllosphere","Lichen","Fungus","Animal","Aquatic"))
      
      # Prepare formula
      formula <- as.formula(paste("~ ",paste(factors, collapse="+"),sep=""))
      x_axis <- "sample.type"
      x_fill <- "sample.type"
      
      # remove if exists from previous dataset
      if (exists("facet")){
        rm(facet)  
      }
      
    } else if (study == "fullcyc"){
      
      # Prepare factor levels  
      factors <- c("enrichment","fraction","substrate","day")
      substrates <- c("control","glucose", "xylose","lactate", "oxalic.acid","glycine","amino.acids","cellulose","palmitic.acid","vanillin")
      days <- c(1,3,6,14,30,48)
      
      if (exists("by_day") & exists("by_fraction")){
        # Prepare formula
        formula <- as.formula(paste("~ ",paste(c("enrichment","fraction","substrate","day"), collapse="+")))
        x_axis <- "fraction"
        x_fill <- "enrichment"
        facet <- as.formula("~ substrate")
        
        factors <- c("enrichment","fraction","substrate","day","rep")
        factor_levels <- list(c("C12","C13","incubated"), seq(from= 4, 29, by=1), substrates, days)
        
        rm(by_day)
        rm(by_fraction)
        
      } else if (exists("by_day")){
        
        # Only compare heavy fractions
        if (exists("only_heavy")){
          p <- subset_samples(p, fraction %in% c(1,2,3,4,5,6,7,8,9,10,11,12) | enrichment == "incubated")  
        }
        
        # Prepare formula
        formula <- as.formula(paste("~ ",paste(c("enrichment","substrate","day"), collapse="+")))
        x_axis <- "enrichment"
        x_fill <- "enrichment"
        facet <- as.formula("~ substrate + day")
        
        factors <- c("enrichment","substrate","day","rep")
        factor_levels <- list(c("C12","C13","incubated"), substrates, days)
        
        rm(by_day)
        
      } else if (exists("by_fraction")){
        
        # Prepare formula
        formula <- as.formula(paste("~ ",paste(c("enrichment","fraction","substrate"), collapse="+")))
        x_axis <- "fraction"
        x_fill <- "enrichment"
        facet <- as.formula("~ substrate")
        
        factors <- c("enrichment","fraction","substrate","rep")
        factor_levels <- list(c("C12","C13","incubated"), seq(from= 4, 29, by=1), substrates)
        
        rm(by_fraction)
        
      } else {
        
        # Only compare heavy fractions
        if (exists("only_heavy")){
          p <- subset_samples(p, fraction %in% c(1,2,3,4,5,6,7,8,9,10,11,12) | enrichment == "incubated")  
        }
        
        # Prepare formula
        formula <- as.formula(paste("~ ",paste(c("enrichment","substrate"), collapse="+")))
        x_axis <- "substrate"
        x_fill <- "enrichment"
        
        factors <- c("enrichment","substrate","rep")
        factor_levels <- list(c("C12","C13","incubated"), substrates)
        
        # remove if exists from previous dataset
        if (exists("facet")){
          rm(facet)  
        }
      }
    }
    
    ### Plot
    ###
    
    # Get Data and subset to taxon of interest
    if (method == "Taxa"){
      taxa_table <- as(tax_table(p)[,rank], "vector")
      KEEP <- rownames(tax_table(p))[which(taxa_table == taxa)]
      p_taxa <- try(prune_taxa(KEEP, p), silent = TRUE)
      
      # Catch missing Taxa
      if (class(p_taxa) == "try-error") {
        print("This taxa was not found in your dataset.")
        next()
      } 
      
      if (study == "soil.health"){
        # Print geogroup info
        get_geogroups(KEEP,p_taxa)
      }
      
    } else {
      print(OTU)
      p_taxa <- try(prune_taxa(OTU, p), silent = TRUE)
      
      # Catch missing OTU
      if (class(p_taxa) == "try-error") {
        print("This OTU was not found in your dataset.")
        next()
      } 
      
      if (study == "soil.health"){
        # Print geogroup info
        get_geogroups(OTU,p_taxa)
      }
    }
    
    # Prep counts dataframe with factors
    plot_me <- data.frame(Counts = sample_sums(p_taxa))
    
    if (exists("show_geo_groups") & study == "soil.health"){
      plot_me <- cbind(plot_me, sample_data(p_taxa)[,c(factors,"geo_group")])
      plot_me$geo_group <- as.factor(plot_me$geo_group)
      plot_me <- ddply(plot_me, ~ get(factors) + geo_group, summarise, average = mean(Counts))
      colnames(plot_me) <- c(factors,"geo_group","Counts")
      plot_me[,factors] <- as.numeric(plot_me[,factors])
      
    } else {
      plot_me <- cbind(plot_me, sample_data(p_taxa)[,factors])
    }
    
    # Collapse SIP data by replicates
    if (study == "fullcyc"){
      formula2 <- as.formula(paste(as.character(formula)[1],as.character(formula)[2], "+rep", sep=""))
      plot_me <- ddply(plot_me, formula2, summarise, counts = mean(Counts))
      colnames(plot_me)[grep("counts", colnames(plot_me))] <- "Counts"
    }
    
    # Calculate standard error and mean for barplot plot type
    if (plot.type != "dot"){
      error <- ddply(plot_me, formula,  summarise, error = std(Counts))
      plot_me <- ddply(plot_me, formula, summarise, average = mean(Counts))
      plot_me <- cbind(plot_me, error = error$error)
    } else {
      if (exists("remove_zeros")){
        plot_me <- subset(plot_me, Counts > 0)
        rm(remove_zeros)
      }
    }
    
    ## Special treatment for fullcyc: 
    ##     i) Add in Control for every substrate
    ##     ii) remove NA fractions
    
    if (study == "fullcyc"){
      
      # Double the control in SIP studies
      #foo <- subset(plot_me, substrate == "control")
      
      #for (substrate in substrates){
      #  foo$substrate <- substrate
      #  plot_me <- rbind(plot_me, foo)  
      #}
      
      # Remove NA and incubated
      if (exists("by_fraction")){
        plot_me <- subset(plot_me, !is.na(fraction))
      }
      
      plot_me <- subset(plot_me, enrichment != "incubated")  
    } 
    
    # Special treatment for chazy.SIP i) Add in Control for every substrate
    if (study == "chazy.SIP"){
      
      # Double the control in SIP studies
      foo.NTH <- subset(plot_me, substrate == "control" & treatment == "NTH")
      foo.PTH <- subset(plot_me, substrate == "control" & treatment == "PTH")
      
      for (subs in c("xylose","cellulose")){
        foo.NTH$substrate <- subs
        plot_me <- rbind(plot_me, foo.NTH)  
        foo.PTH$substrate <- subs
        plot_me <- rbind(plot_me, foo.PTH)
      }
      
      # Remove NA
      if (exists("by_fraction")){
        plot_me <- subset(plot_me, !is.na(fraction))
      }
      
      plot_me <- subset(plot_me, substrate != "control")
    } 
    
    # Set Factor Levels
    if (exists("factor_levels")){
      
      for (n in 1:length(factor_levels)){
        plot_me[,factors[n]] <- factor(plot_me[,factors[n]], levels = unlist(factor_levels[n]))
      }
      
      rm(factor_levels)
    }
    
    ## Cycle Through Plots by Day for SIP studies
    if (study == "fullcyc" | study == "chazy.SIP"){
      if(exists("by_day")){
        for (d in days){
          foo <- subset(plot_me, day == d)
          
          # Skip if empty
          if (nrow(foo) == 0){
            next
          }
          
          # remove substrates with zero counts (frees up space on plot)
          if (plot.type == "dot"){
            remove <- ddply(foo, ~ substrate, summarise, total = sum(Counts))
            foo <- subset(foo, !(substrate %in% as.character(subset(remove, total == 0)$substrate)))
          } else {
            remove <- ddply(foo, ~ substrate, summarise, total = sum(average))
            foo <- subset(foo, !(substrate %in% as.character(subset(remove, total == 0)$substrate)))
          }
          
          # Create base plot
          if (exists("show_geo_groups") & study == "soil.health"){
            if (plot.type == "dot"){
              #plot <- ggplot(plot_me, aes(x=get(x_axis), y=Counts, colour = geo_group)) + geom_boxplot(outlier.shape = NA) + ylab("Average Counts per Thousand Reads") 
              #plot <- plot + geom_point(position=position_jitterdodge(), alpha=0.5)
              plot <- ggplot(plot_me, aes(x = get(x_axis), y = Counts, colour = geo_group)) + geom_smooth(method=lm, se = FALSE) + geom_point()+ ylab("Average Counts per Thousand Reads") 
              
            } else {
              plot <- ggplot(foo, aes(x=get(x_axis), y=average, fill = geo_group)) + geom_bar(position="dodge", stat="identity") + ylab("Average Counts per Thousand Reads")
            }
            
            rm("show_geo_groups")
            
          } else {
            if (plot.type == "dot"){
              plot <- ggplot(plot_me, aes(x=get(x_axis), y=Counts, colour = get(x_fill))) + geom_boxplot(outlier.shape = NA) + ylab("Average Counts per Thousand Reads") 
              plot <- plot + geom_point(position=position_jitterdodge(), alpha=0.5)
              
            } else {
              plot <- ggplot(foo, aes(x=get(x_axis), y=average, fill = get(x_fill))) + geom_bar(position="dodge", stat="identity") + ylab("Average Counts per Thousand Reads")
            }
          }
          
          # Add in title
          if (method == "Taxa"){
            plot <- plot + ggtitle(paste("Relative Abundance of:",taxa,"in",study,"on day",d,sep=" "))
          } else {
            if (length(OTU) == 1){
              plot <- plot + ggtitle(paste("Relative Abundance of:",paste(tax_table(p_taxa),collapse=";"),"in",study,"on day",d,sep=" "))
            } else {
              plot <- plot + ggtitle(paste("Relative Abundance of:",paste(taxa_names(p_taxa),collapse=","),"in",study,"on day",d,sep=" "))
            }
          }
          
          # Add facet
          if (exists("facet")){
            plot <- plot + facet_grid(facet)
          }
          
          # Add error bars (there are no replicates in fullcyc)
          if (study == "chazy.SIP" & plot.type != "dot"){
            plot <- plot + geom_errorbar(limits, position=position_dodge(0.9), width=0.25)  
          }
          
          # Add better labels
          plot <- plot + guides(fill=guide_legend(title=x_axis)) + xlab(x_axis)
          
          # Print to screen
          print(paste("Day",d,sep=" "))
          print(plot)
          
          writeLines("Save plots?")
          switch(menu(c("Discard","Continue","Exit")), step1 <- "Discard", step1 <- "Save to File", step1 <- "Exit")
          
          if (step1 == "Exit"){
            break
          } else if (step1 == "Save to File"){
            if (method == "Taxa"){
              ggsave(plot, filename=paste(taxa,study,d,'relabund',plot.type,'pdf',sep="."), height=16, width=16)
            } else {
              if (length(OTU) > 3){
                ggsave(plot, filename=paste(paste(OTU[1:3], collapse="."),"abr",study,d,'relabund',plot.type,'pdf',sep="."), height=16, width=16)
                
              } else {
                ggsave(plot, filename=paste(paste(OTU, collapse="."),study,d,'relabund',plot.type,'pdf',sep="."), height=16, width=16)
              }
            }
            
            ## Print Stats
            if (plot.type == "dot"){
              # Remove zeros
              #plot_me <- subset(plot_me, Counts > 0)
              
              # Averages
              print(ddply(plot_me, ~get(x_axis), summarise, average = mean(Counts)))
              
              # Stats
              fit <- lm(Counts ~get(x_axis), data= plot_me)
              print(TukeyHSD((aov(fit))))
              
              print("Wilcoxon")
              print(wilcox.test(Counts ~get(x_axis), data= plot_me))
              
            }
          }
        }
      } else {
        foo <- plot_me
        
        # Skip if empty      
        if (nrow(foo) == 0){
          next
        }
        
        # remove substrates with zero counts (frees up space on plot)
        if (plot.type == "dot"){
          remove <- ddply(foo, ~ substrate, summarise, total = sum(Counts))
          foo <- subset(foo, !(substrate %in% as.character(subset(remove, total == 0)$substrate)))
        } else {
          remove <- ddply(foo, ~ substrate, summarise, total = sum(average))
          foo <- subset(foo, !(substrate %in% as.character(subset(remove, total == 0)$substrate)))
        }
        
        # Create base plot
        if (exists("show_geo_groups") & study == "soil.health"){
          if (plot.type == "dot"){
            #plot <- ggplot(plot_me, aes(x=get(x_axis), y=Counts, colour = geo_group)) + geom_boxplot(outlier.shape = NA) + ylab("Average Counts per Thousand Reads") 
            #plot <- plot + geom_point(position=position_jitterdodge(), alpha=0.5)
            plot <- ggplot(plot_me, aes(x = get(x_axis), y = Counts, colour = geo_group)) + geom_smooth(method=lm, se = FALSE) + geom_point()+ ylab("Average Counts per Thousand Reads") 
            
          } else {
            plot <- ggplot(foo, aes(x=get(x_axis), y=average, fill = geo_group)) + geom_bar(position="dodge", stat="identity") + ylab("Average Counts per Thousand Reads")
          }
          
          rm("show_geo_groups")
          
        } else {
          if (plot.type == "dot"){
            plot <- ggplot(plot_me, aes(x=get(x_axis), y=Counts, colour = get(x_fill))) + geom_boxplot(outlier.shape = NA) + ylab("Average Counts per Thousand Reads") 
            plot <- plot + geom_point(position=position_jitterdodge(), alpha=0.5)
            
          } else {
            plot <- ggplot(foo, aes(x=get(x_axis), y=average, fill = get(x_fill))) + geom_bar(position="dodge", stat="identity") + ylab("Average Counts per Thousand Reads")
          }
        }
        
        # Add in title    
        if (method == "Taxa"){
          plot <- plot + ggtitle(paste("Relative Abundance of:",taxa,"in",study,sep=" "))
        } else {
          if (length(OTU) == 1){
            plot <- plot + ggtitle(paste("Relative Abundance of:",paste(tax_table(p_taxa),collapse=";"),"in",study,sep=" "))
          } else {
            plot <- plot + ggtitle(paste("Relative Abundance of:",paste(taxa_names(p_taxa),collapse=","),"in",study,sep=" "))
          }
        }
        
        # Add facet
        if (exists("facet")){
          plot <- plot + facet_grid(facet)
        }
        
        # Add error bars
        if (plot.type != "dot"){
          plot <- plot + geom_errorbar(limits, position=position_dodge(0.9), width=0.25)  
        }
        
        # Add better labels
        plot <- plot + guides(fill=guide_legend(title=x_axis)) + xlab(x_axis)
        
        # Print to screen
        print(plot)
        
        writeLines("Save plots?")
        switch(menu(c("Discard","Continue","Exit")), step1 <- "Discard", step1 <- "Save to File", step1 <- "Exit")
        
        if (step1 == "Exit"){
          break
        } else if (step1 == "Save to File"){
          if (method == "Taxa"){
            ggsave(plot, filename=paste(taxa,study,'relabund',plot.type,'pdf',sep="."), height=16, width=16)
          } else {
            if (length(OTU) > 3){
              ggsave(plot, filename=paste(paste(OTU[1:3], collapse="."),"abr",study,'relabund',plot.type,'pdf',sep="."), height=16, width=16)  
              
            } else {
              ggsave(plot, filename=paste(paste(OTU, collapse="."),study,'relabund',plot.type,'pdf',sep="."), height=16, width=16)  
            }
            
          }
          
          ## Print Stats
          if (plot.type == "dot"){
            # Remove zeros
            #plot_me <- subset(plot_me, Counts > 0)
            
            # Averages
            print(ddply(plot_me, ~get(x_axis), summarise, average = mean(Counts)))
            
            # Stats
            fit <- lm(Counts ~ get(x_axis), data= plot_me)
            print(TukeyHSD((aov(fit))))
            
            print("Wilcoxon")
            print(wilcox.test(Counts ~get(x_axis), data= plot_me))
            
          }
        }
      }
    } else {
      
      # Create base plot
      if (exists("show_geo_groups") & study == "soil.health"){
        if (plot.type == "dot"){
          #plot <- ggplot(plot_me, aes(x=get(x_axis), y=Counts, colour = geo_group)) + geom_boxplot(outlier.shape = NA) + ylab("Average Counts per Thousand Reads") 
          #plot <- plot + geom_point(position=position_jitterdodge(), alpha=0.5)
          plot <- ggplot(plot_me, aes(x = get(x_axis), y = Counts, colour = geo_group)) + geom_smooth(method=lm, se = FALSE) + geom_point()+ ylab("Average Counts per Thousand Reads") 
          
        } else {
          plot <- ggplot(plot_me, aes(x=get(x_axis), y=average, fill = geo_group)) + geom_bar(position="dodge", stat="identity") + ylab("Average Counts per Thousand Reads") 
        }
        
        rm("show_geo_groups")
        
      } else {
        if (plot.type == "dot"){
          plot <- ggplot(plot_me, aes(x=get(x_axis), y=Counts, colour = get(x_fill))) + geom_boxplot(outlier.shape = NA) + ylab("Average Counts per Thousand Reads") 
          plot <- plot + geom_point(position=position_jitterdodge(), alpha=0.5)
          
        } else {
          plot <- ggplot(plot_me, aes(x=get(x_axis), y=average, fill = get(x_fill))) + geom_bar(position="dodge", stat="identity") + ylab("Average Counts per Thousand Reads") 
        }
      }
      
      
      # Add title
      if (method == "Taxa"){
        plot <- plot  + ggtitle(paste("Relative Abundance of:",taxa,"in",study,sep=" "))
      } else {
        if (length(OTU) == 1){
          plot <- plot + ggtitle(paste("Relative Abundance of:",paste(tax_table(p_taxa),collapse=";"),"in",study,sep=" "))
        } else {
          plot <- plot + ggtitle(paste("Relative Abundance of:",paste(taxa_names(p_taxa),collapse=","),"in",study,sep=" "))
        }
      }
      
      # Add facet
      if (exists("facet")){
        plot <- plot + facet_grid(facet)
        rm(facet)
      }
      
      # Add error bars
      if (plot.type != "dot"){
        plot <- plot + geom_errorbar(limits, position=position_dodge(0.9), width=0.25)
      }
      
      # Add better labels
      plot <- plot + guides(fill=guide_legend(title=x_axis)) + xlab(x_axis)
      
      # Print to screen
      print(plot)
      
      writeLines("Save plots?")
      switch(menu(c("Discard","Continue","Exit")), step1 <- "Discard", step1 <- "Save to File", step1 <- "Exit")
      
      if (step1 == "Exit"){
        break
      } else if (step1 == "Save to File"){
        if (method == "Taxa"){
          ggsave(plot, filename=paste(taxa,study,'relabund',plot.type,'pdf',sep="."), height=16, width=16)
        } else {
          if (length(OTU) > 3){
            ggsave(plot, filename=paste(paste(OTU[1:3], collapse="."),"abr",study,'relabund',plot.type,'pdf',sep="."), height=16, width=16)
            
          } else {
            ggsave(plot, filename=paste(paste(OTU, collapse="."),study,'relabund',plot.type,'pdf',sep="."), height=16, width=16)
          }
        }
        
        ## Print Stats
        if (plot.type == "dot"){
          # Remove zeros
          #plot_me <- subset(plot_me, Counts > 0)
          
          # Averages
          print(ddply(plot_me, ~get(x_axis), summarise, average = mean(Counts)))
          
          # Stats
          fit <- lm(Counts ~get(x_axis), data= plot_me)
          print(TukeyHSD((aov(fit))))
          
          print("Wilcoxon")
          print(wilcox.test(Counts ~get(x_axis), data= plot_me))
          
        }
      }
    }
}
```
