## Supplementary material for "Phenolic acid-degrading *Paraburkholderia* prime decomposition in forest soil": calculate_ratio.R

```
## Calculate ratio of C12 : C13
calculate_ratio <- function(decision){
  
  if (decision == "N"){
    # ratio of C12 : C13 determined from masses of 13C and 12C substrate mixed
    mg.C13.glucose <- 36.6  # 99 atom % 13C
    mg.C12.glucose <- 172.6 # 1.1 atom % 13C
    
    mg.C13.PHB <- 42.6  # 99 atom % 13C
    mg.C12.PHB <- 200.4 # 1.1 atom % 13C
    
    # calculation (adjust mass of 13C by 99%)
    ratio.glucose = (mg.C13.glucose*0.01+mg.C12.glucose)/mg.C13.glucose*0.99
    ratio.PHB = (mg.C13.PHB*0.01+mg.C12.PHB)/mg.C13.PHB*0.99
  
  } else {
    # use isotopic data from COIL
    coil <- read.csv(file = "raw_data/COIL.data.csv", header = T, stringsAsFactors = F)
    
    # Convert atom percent to atom proportion
    coil$atom.C13 <- coil$atom.C13/100
    
    # Solutions containing 13C-substrates were diluted 1:1 with a 12C-glucose dilution solution 100 mg/mL
    # The labeled glucose and PHB solutions were at concentrations of 5 mg/mL and 2 mg/mL, respectively
    # 100uL of sample solution was mixed with 100 uL of 12C-glucose dilution solution and run on EA-IRMS
    
    ## mols C13 contributed from C12 dilution
    mg.glucose.dilution = (0.1 * 100)  # (100 uL 100 mg/mL C12-glucose)
    mols.glucose.dilution = (mg.glucose.dilution/1000)/180.156  # 180.156 g/mol
    mols.C.dilution = mols.glucose.dilution*6  # 6 carbons per mol of glucose
    mols.C13.dilution = mols.C.dilution*coil$atom.C13[which(coil$amendment == "C12-glucose")]
    
    ## Glucose
    
    # mols C13 in total sample
    mg.glucose = (0.1 * 5)+(0.1 * 100)  # (100 uL 5 mg/mL C13-glucose) * (100 uL 100 mg/mL C12-glucose)
    mols.glucose = (mg.glucose/1000)/180.156  # 180.156 g/mol
    mols.C = mols.glucose*6  # 6 carbons per mol of glucose
    mols.C13 = mols.C*coil$atom.C13[which(coil$amendment == "C13-glucose")]
    
    # mols C13 contributed to mix from sample
    mols.C13.glucose.COIL = mols.C13 - mols.C13.dilution
    
    # mols C in sample
    mg.glucose.sample = (0.1 * 5)  # (100 uL 5 mg/mL glucose)
    mols.glucose.sample = (mg.glucose.sample/1000)/180.156  # 180.156 g/mol
    mols.C.sample = mols.glucose.sample*6  # 6 carbons per mol of glucose
    
    # atom % C13 of undiluted sample
    (mols.C13.glucose.COIL/mols.C.sample)*100
    
    # calculate ratio of C12:C13
    ratio.glucose <- (1-(mols.C13.glucose.COIL/mols.C.sample))/(mols.C13.glucose.COIL/mols.C.sample)
    
    ## PHB
    
    # mols C13 in total sample
    mols.C = (((0.1 * 2)/1000)/138.12)*7 + (((0.1 * 100)/1000)/180.156)*6 # 138.12 g/mol PHB + 180.156 g/mol glucose
    # 7 carbons per mol of PHB and * 6 carbons
    mols.C13 = mols.C*coil$atom.C13[which(coil$amendment == "C13-PHB")]
    
    # mols C13 contributed to mix from sample
    mols.C13.PHB.COIL = mols.C13 - mols.C13.dilution
    
    # mols C in sample
    mg.PHB.sample = (0.1 * 2)  # (100 uL 2 mg/mL PHB)
    mols.PHB.sample = (mg.PHB.sample/1000)/138.12  # 138.12 g/mol
    mols.C.sample = mols.PHB.sample*7  # 7 carbons per mol of PHB
    
    # atom % C13 of undiluted sample
    (mols.C13.PHB.COIL/mols.C.sample)*100
    
    # calculate ratio of C12:C13
    ratio.PHB <- (1-(mols.C13.PHB.COIL/mols.C.sample))/(mols.C13.PHB.COIL/mols.C.sample)
    
  }
  
  return(c(ratio.glucose, ratio.PHB)) 
}
```
