## Supplementary material for "Phenolic acid-degrading *Paraburkholderia* prime decomposition in forest soil": Wilhelm and DeRito - Supplementary Figures.pdf

**Figure S1**. Photographs of field work measuring the respiration of $^{13}$C-labeled PHB (A-D) and images of soils from each forest type (E, F, and G). Soils dosed with PHB were eventually collected for DNA extraction and density gradient ultracentrifugation to recover $^{13}$C-enriched ‘heavy’ DNA corresponding to the PHB-degrading populations. In (A), a close-up of metal chambers used for collecting soil $CO_2$ efflux deployed in the red pine ecoplot. In (B), the application of 150 µL of $^{13}$C-labeled or unlabeled PHB to soil in chambers embedded in the sugar maple ecoplot. In (C), sampling headspace for GC/MS analysis from triplicates in sugar maple plantation soil. In (D), a view of the landscape during field sampling in November of 2016. Macro images of field soils reveal differences in root density among forest types, where the dense root structures of sugar maple (E) where absent in red pine (F) and black locust (G). All root material was removed by sieving prior to microcosm soil priming experiment. Image scale differs slightly among E, F and G.

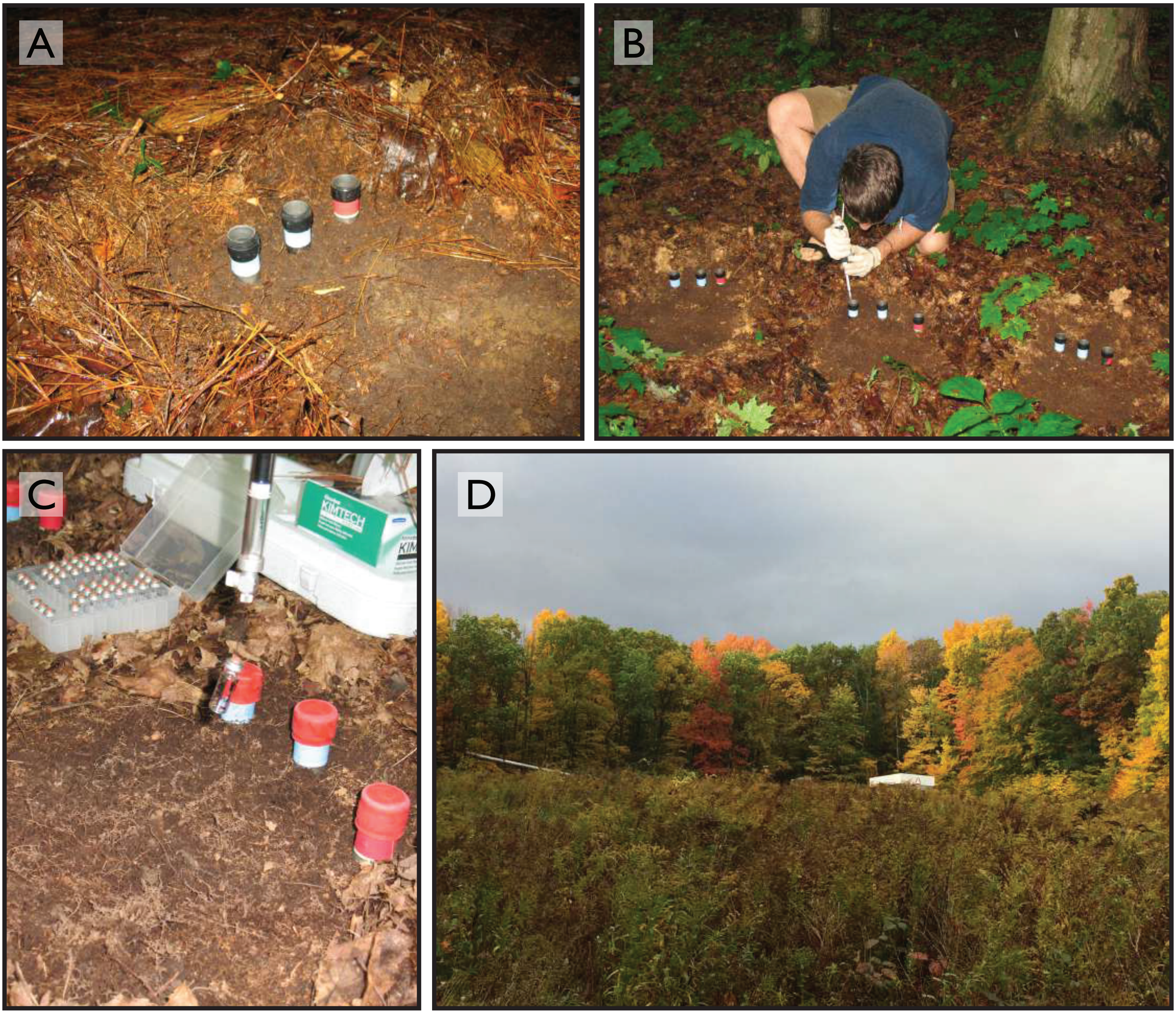


Photo credit: Dr. Eugene Madsen

**Figure S1** (continued)

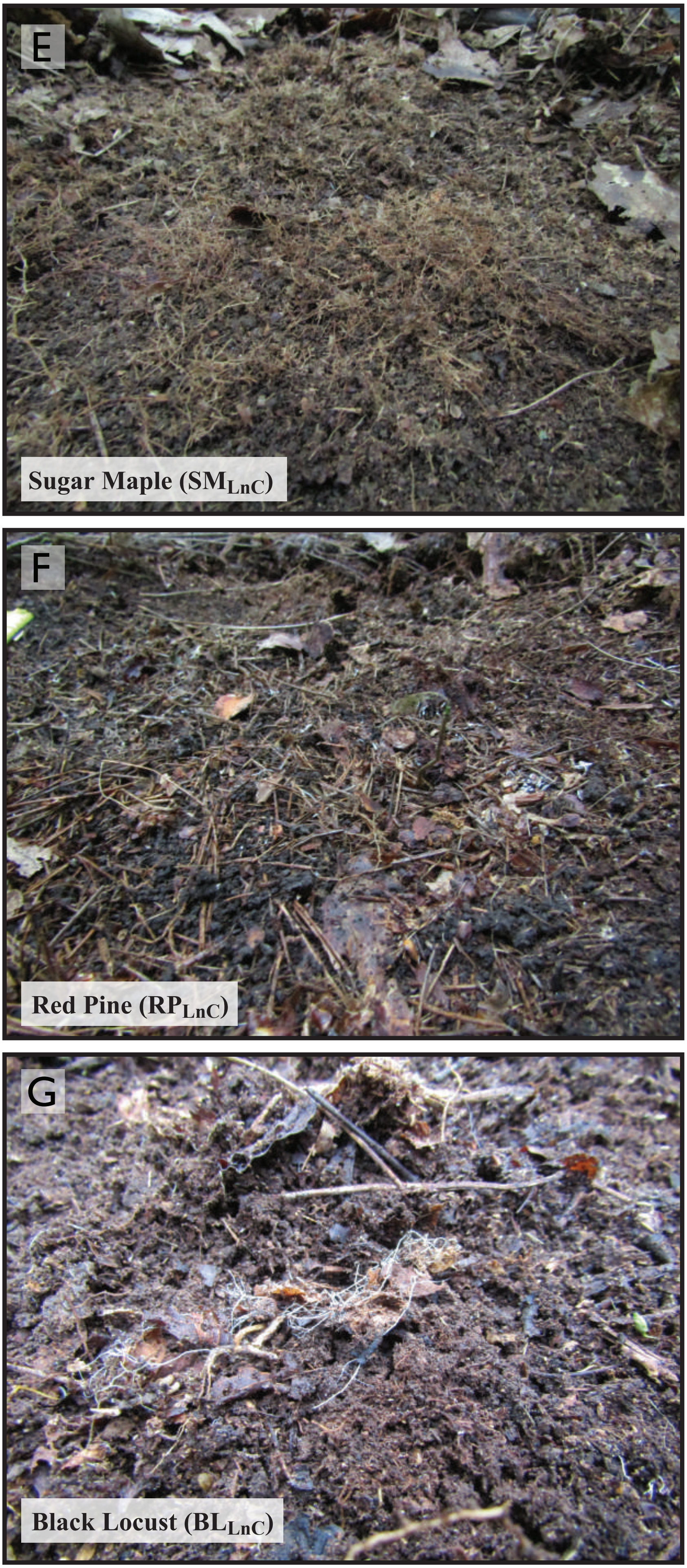


Photo credits: Dr. Roland Wilhelm

**Figure S2**. The design of the soil priming experiment displaying preparation methods, a numerically accurate representation of sample numbers, and a rationale for each treatment.

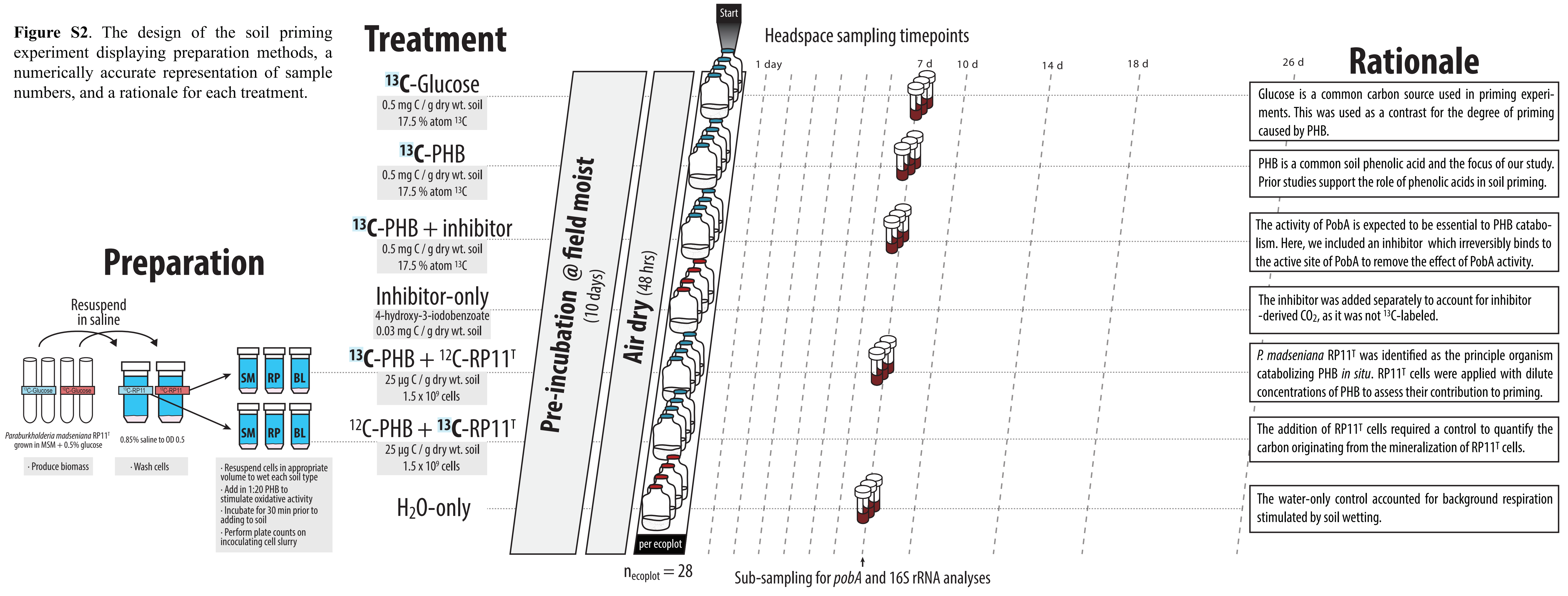

**Figure S3**. Trends in soil microcosm treatments (glucose and RP11$^T$ cell amendments) where negative soil priming was observed (A and B) corresponded with the relative abundance of *Streptomycetaceae* phylotypes (C, D, and E). In (A), the spontaneous priming of native SOC over time and (B) the cumulative respiration of amended substrate is shown. Phylotypes were named according to the top representative BLAST hit to a type strain (100% identity). Taxonomic classifications correspond to a 'species group' since the V4 region of the 16S rRNA gene cannot resolve closely related species. In (C), stacked barplots display the relative abundance of all phylotypes in DNA extracts. In (D), the total relative abundance of *Streptomycetaceae* in DNA and cDNA pools are shown for soil microcosms amended with $^{13}$C-labeled RP11$^T$ cells.

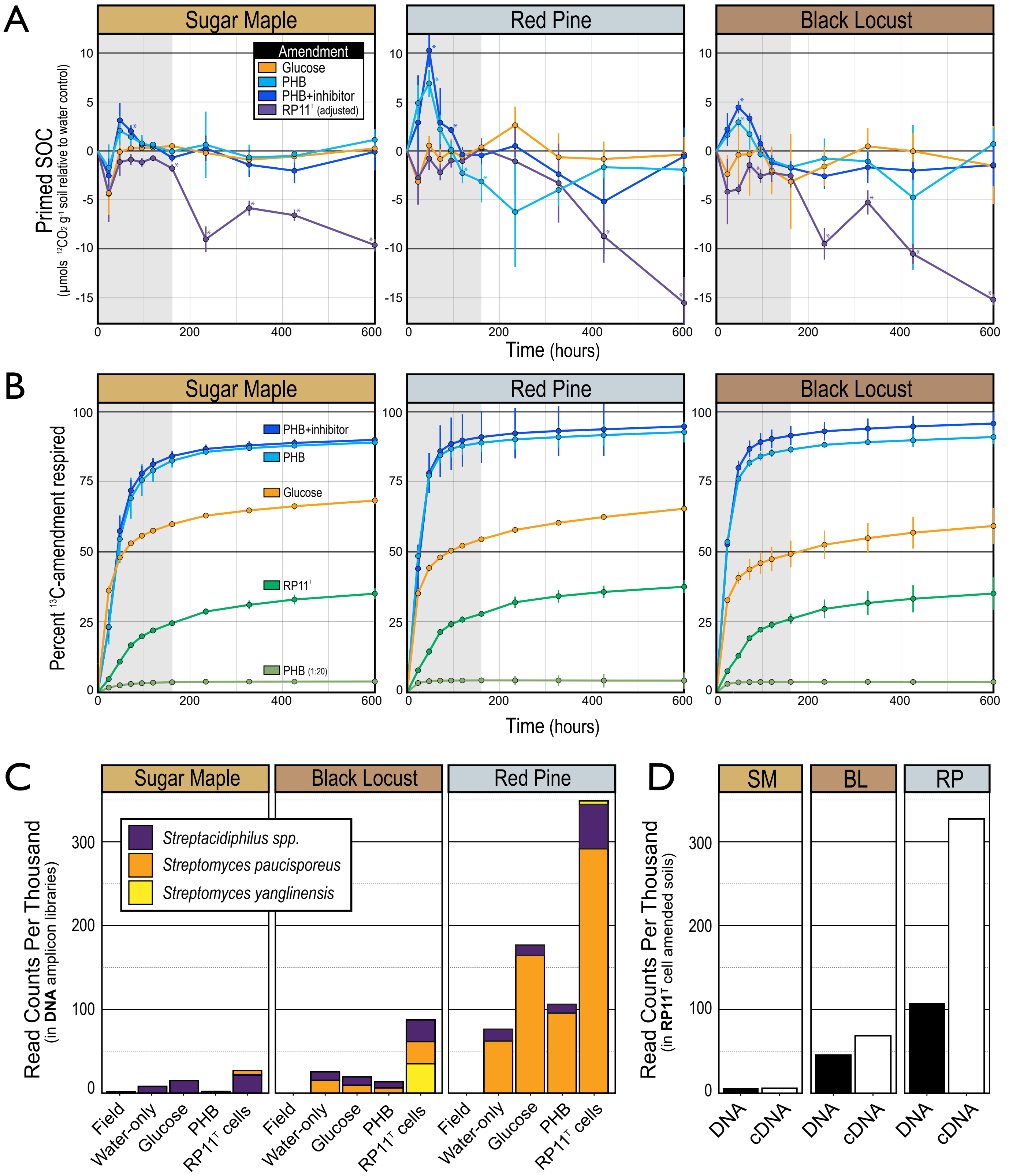

**Figure S4**. The size of active bacterial populations differed among ecoplots and was directly proportional to soil organic matter, as revealed in (A) RNA yields from DNAse treated soil extractions, (B) the abundance of copies of 16S rRNA according to RT-qPCR, and (C) the percent soil organic matter of field soil. In (A), RNA yields were measured from unamended and PHB-amended field soil in parallel for all ecoplots. In (B), RT-qPCR was performed on the RNA extracts measured in (A). Letters denote significant differences ($p < 0.05$) based on Tukey's Honest Significant Difference tests (n = 3).

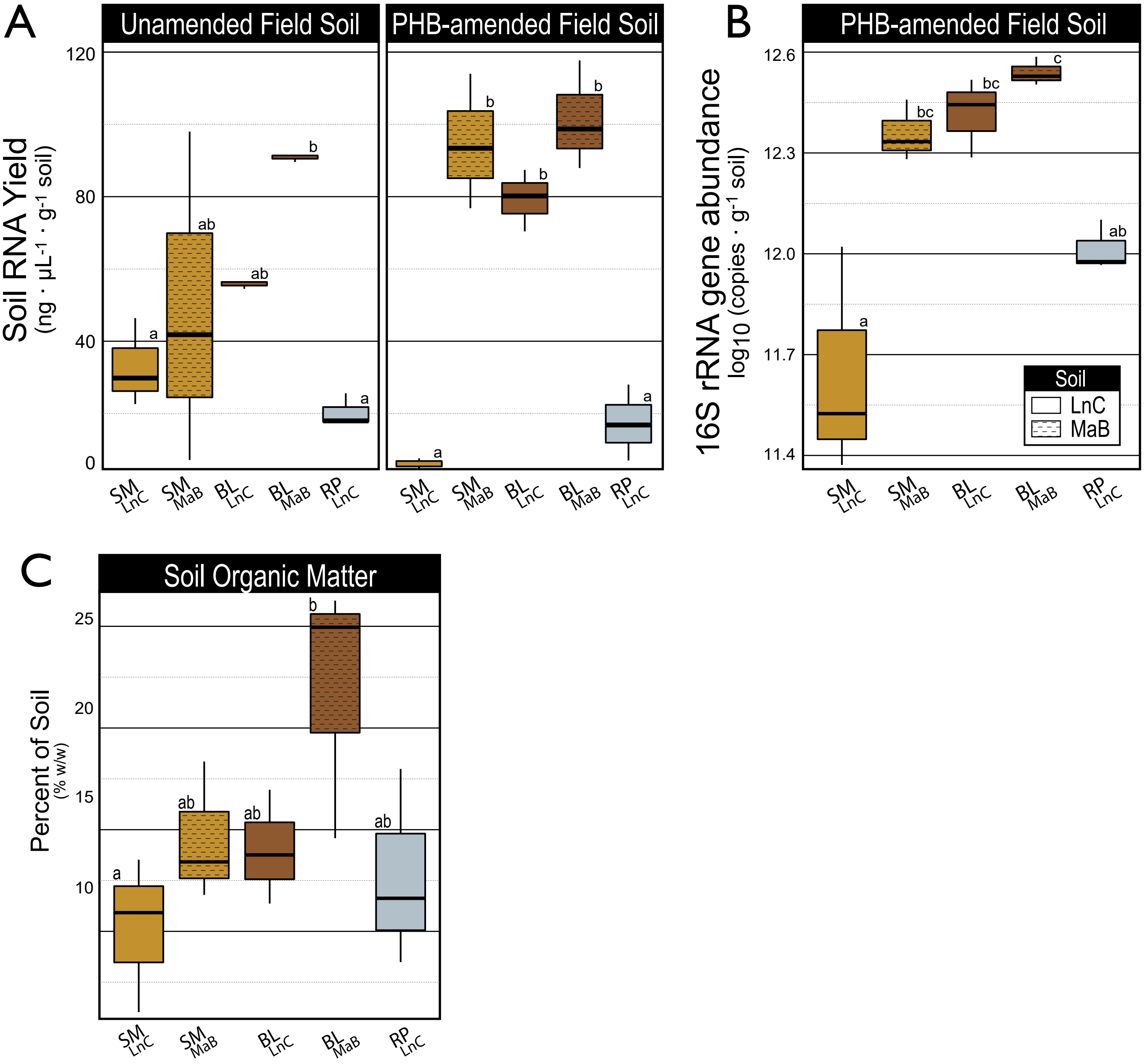

**Figure S5**. Species richness (A) and Pielou's evenness (B) of $^{13}C$-enriched versus control 16S rRNA gene libraries iin heavy gradient fractions (gradient fractions F3-F8) in all ecoplots. Asterisks denote significant differences ($p < 0.05$) based on Mann–Whitney tests (n=3 per). The average read depth per 16S rRNA gene library was  80,000 sequences, with a minimum of 2,200.

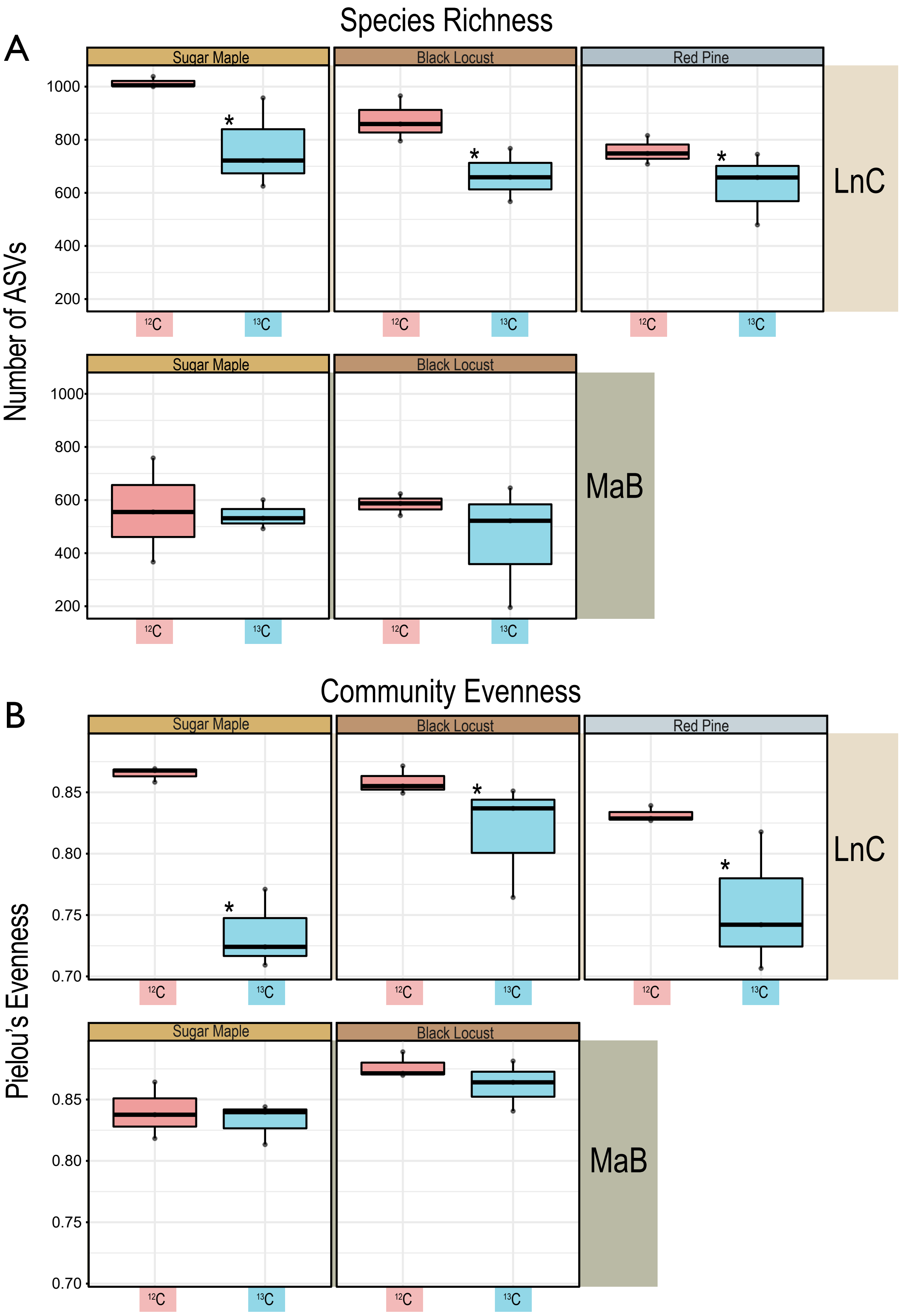

**Figure S6**. The $^{13}C$-enrichment of a *Solicoccozyma* phylotype evidenced by their increased relative abundance in heavy fractions of the CsCl gradients. Blue colored bars correspond to soil DNA extracts from soils incubated with $^{13}C$-labeled PHB, and pink bars to corresponding soils incubated with unlabeled PHB. The 18S rRNA gene was incidentally amplified using our universal bacterial primers (515f/805r). *Solicoccozyma* are common forest soil decomposers [124] and have been reported to degrade PHB [125,126].

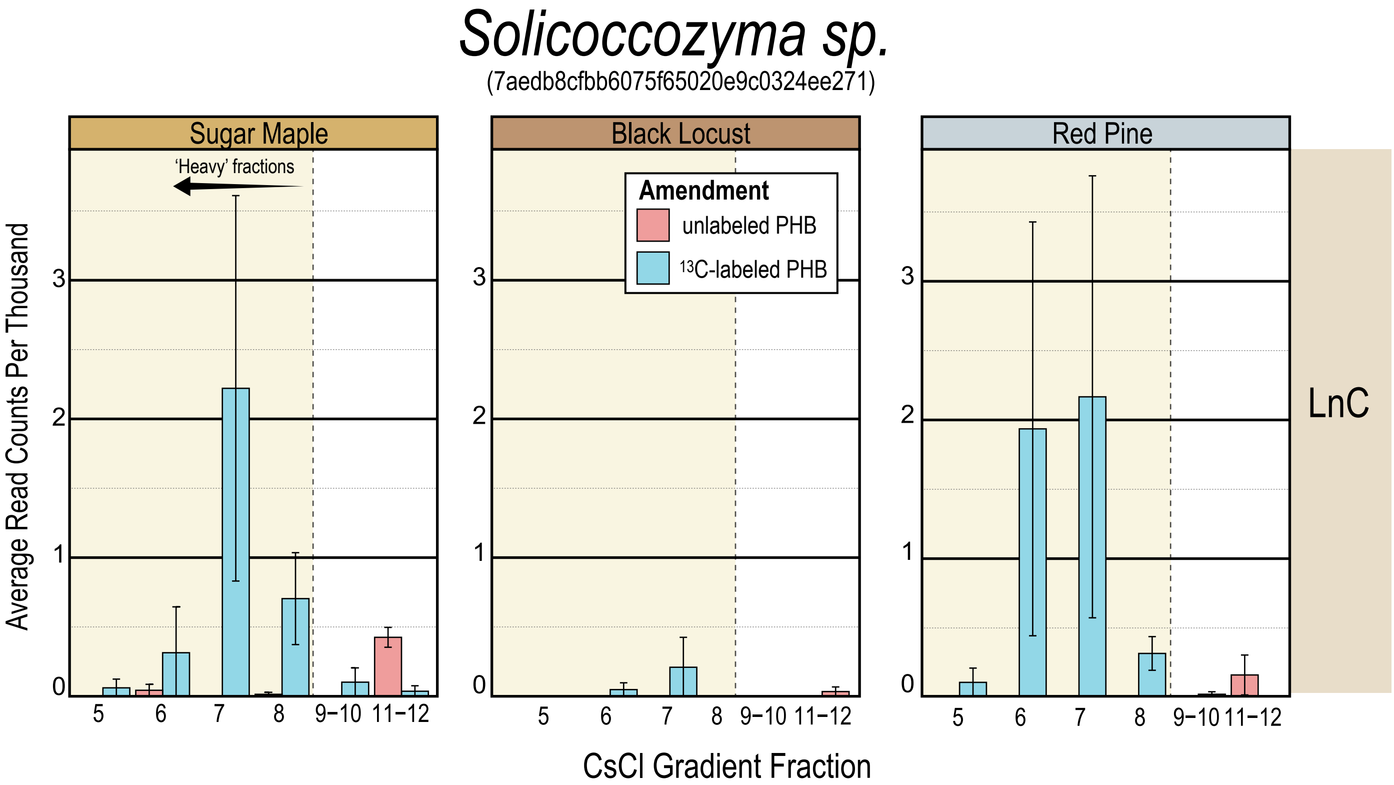

**Figure S7**. The gene for *p*-hydroxybenzoate 3-monooxygenase (*pobA*; EC 1.14.13.2) is encoded by numerous *Burkholderiaceae* genomes (n = 3, 991), and duplicated *pobA* paralogs occur in many of these genomes (n = 410), as revealed by the maximum-likelihood phylogeny constructed from 24 single-copy genes shown in (A). *Burkholderia* were subdivided into sub-genera based on phylogenetic relatedness and presence (clades A, B and C) or absence (clade D) of *pobA* paralogs. Pie charts representing each clade indicate the isolation source of genomes that have one (indicated by a single black dot) or two (indicated by two black dots) copies of *pobA*, shown inset in (A). The genera *Caballeronia* and *Paraburkholderia* are paraphyletic suggesting that phylogenetic relationships within the *Burkholderiaceae* require further refinement, as previously reported [127]. The maximum likelihood phylogeny of *pobA* encoded by *Burkholderiaceae* is comprised of seven major clades, shown in (B). The leaves of the tree are colored with respect to genus, and the proportion of genera that comprise each *pobA* clade is inset. In clade 5, the category ‘others’ includes unclassified *Burkholderiaceae*, *Lautropia*, *Polynucleobacter*, *Robbsia*, and *Paucimonas* (in order of relative abundance) and, in clade 7 the category ‘others’ includes *Trinickia*. Phylogenetic trees in Newick format and HMM models for all *pobA* clades are available in the Supplementary Data package.

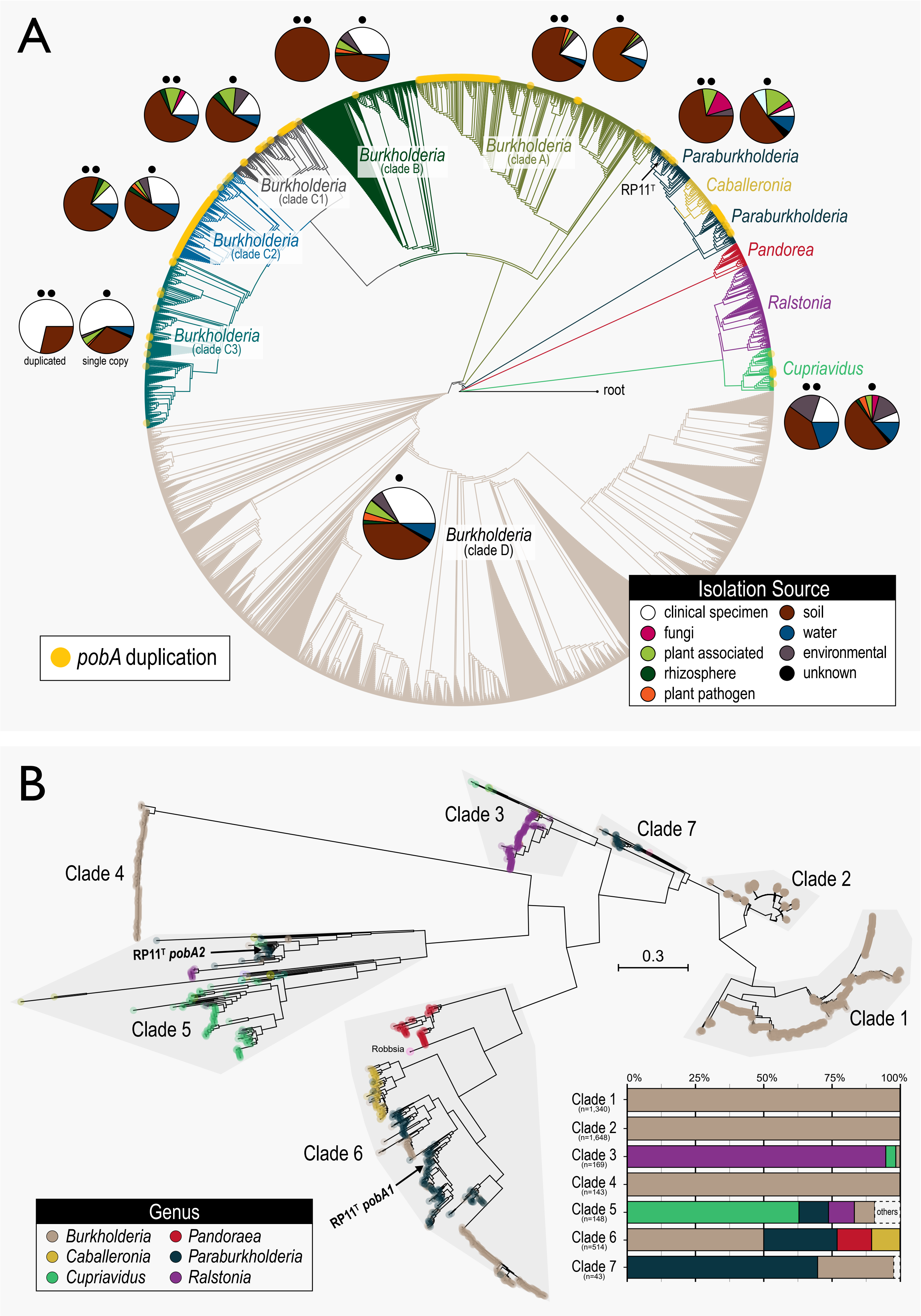
