## Supplementary material for "Phenolic acid-degrading *Paraburkholderia* prime decomposition in forest soil": Wilhelm and DeRito - Supplementary Methods.docx

From Wilhelm and DeRito *et al.*, 2020 – Phenolic acid-degrading populations of *Paraburkholderia* prime decomposition in forest soils.

*Description of field site and sampling campaigns*

The Turkey Hill Forest Plantation (THFP) is an experimental forest designed to capture variation in patterns of vegetation and soil types typical of northeastern USA. The THFP was established between 1939 and 1941 by Cornell forest ecologist Robert F. Chandler. Former agriculture land on the University of Cornell’s Mt. Pleasant property (Dryden, NY) was divided into approximately thirty 0.4 ha plots [1, 2]. These plots were individually planted in duplicate with monospecific plantations of black locust (*Robinia pseudoacacia* L.)*,* sugar maple (*Acer saccharum* Marsh), white ash (*Fraxinus americana* L.), Norway spruce (*Picea abies* L. Karst), northern red oak (*Quercus rubra* L.) and red pine (*Pinus resinosa* Ait.). Non-replicated plots were planted with six additional tree species: black cherry (*Prunus serotina* Ehrh.), red maple (*Acer rubrum* L.), tulip poplar (*Liriodendron tulipifera* L.), American basswood (*Tilia americana* L.), quaking aspen (*Populus tremuloides* L.) and white pine (*Pinus strobus* L.). Trees were planted in 1.5 m x 1.5 m spacing and received little management after planting. Soils at the THFP developed from silt-enriched glacial (Wisconsin) till derived primarily from local Devonian sandstones and siltstones. The soils are primarily a coarse-loamy, mixed, mesic Typic Fragiochrept (Inceptisols) on ~ 5% slopes [2]. Soils at the THFP belong to several different soil series (Bath, Mardin, Valois, Lordstown, Erie). The organic matter content of the upper 4 cm of soils planted to the different tree species at THFP varies approximately 4-fold (highest for Norway spruce and lowest for black cherry). The site was cultivated for approximately 100 years prior to tree planting with a plow pan occurring at 20 cm [3]. Plots at the THFP were found to differ in the abundance of earthworms, with high abundances occurring black locust (196 worms · m^-2^) and substantially lower in sugar maple (25 worms · m^-2^) and red pine (4 worms · m^-2^) [4]. The *in situ* stable isotope probing field work (i.e. measuring ^13^CO_2_ and soil sampling) was conducted in two separate campaigns. The first campaign was conducted on 7/15/2014 when ecoplots SM_LnC_ and RP_LnC_ were sampled. The second campaign was conducted on 10/19/2016 when BL_LnC_, BL_MaB_, SM_MaB_ were sampled.

*Measuring ^13^CO_2_ in situ (GC/MS instrumentation)*

At each sampling time point, 2.5 mL of headspace was sampled from *in situ* chambers and stored in evacuated 2-mL vials. Shortly after, the vials were depressurized and a subsample of 100-µl gas was injected into a Hewlett Packard 5890 gas chromatograph (Wilmington, DE) equipped with a 5971A mass selective detector. High purity helium was used as the carrier gas and an Agilent Technologies GS-GASPRO column (60m length by 0.32 mm I.D.) was used to separate CO_2_ from other atmospheric gases. Concentrations of CO_2_ were quantified using calibration curves prepared from external standards spanning 3.7 to 75 nmols CO_2_/µL (Airgas, Inc.).

*Extraction of DNA and RNA from soil*

DNA and RNA were extracted simultaneously from soils using a modified phenol-chloroform/bead-beating method from [5]. All vessels and reagents used were either certified RNase free, treated with RNase ZAP (Ambion, Austin, TX), or treated with diethyl pyrocarbonate (DEPC) for inactivation of RNase activity. Briefly, ~0.5g soil was transferred to 2-mL Lysing Matrix E tubes (MP Biomedicals) and 0.5 mL CTAB (hexadecyltrimethylammonium bromide) buffer (10% CTAB in 0.7 M NaCl, 240 mM potassium phosphate buffer, pH 8) and 0.5 mL phenol (pH>7.8) : chloroform:isoamyl alcohol (25:24:1) were added. Tubes were homogenized for 45 sec in a Mini-Bead Beater-8 instrument (Biospec Products), and then centrifuged at 16,000 g for 5 minutes at 4°C. The aqueous layer was transferred to a new 1.5-mL microcentrifuge tube containing an equal volume of chloroform:isoamylalcohol (24:1), vortexed, then centrifuged again. The aqueous layer was transferred to a new tube containing two volumes of 30% (wt/vol) polyethylene glycol 6000/1.6 M NaCl. RNA/DNA was precipitated at 4°C overnight. RNA/DNA pellets were collected by centrifugation (10 min, 16,000 g, 4°C), washed with ice-cold 70% ethanol, air-dried, and resuspended in 30 µl nuclease-free water. Total nucleic acid concentrations were determined using a Qubit® 3.0 fluorometer (ThermoFisher Scientific) and extracts were stored at -80°C. 16S rRNA gene and 16S rRNA libraries were constructed from extracts from the soil priming experiment, while only 16S rRNA gene libraries were constructed for the *in situ* SIP experiment.

*Density gradient ultracentrifugation and recovery of ^13^C-labeled DNA*

Briefly, 5 µg of total nucleic acid were diluted to 1.20 mL with sterile GB buffer (0.1 M Tris, 0.1 M KCl, 1 mM EDTA), mixed with 4.80 mL of 7.163M cesium chloride, and loaded into ultracentrifuge tubes which were then heat-sealed. Tubes were centrifuged at 140,000 g (41,900 rpm; Vti80 rotor) for 66 hours at 20°C. CsCl density was determined based on the refractive index of solution measured with an AR200 digital refractometer (Leica Microsystem). Reagent blanks (no added DNA) and a positive controls were processed identically, the latter consisted of 5 µg each of ^12^C- and ^13^C-DNA, prepared from cultures of *Pseudomonas putida* G7 grown in with either ^12^C- or ^13^C_6_-glucose as the sole carbon source (Sigma-Aldrich, St. Louis, MO).

*Characterizing PHB-degrading activity of isolates*

Nine isolates from the THFP (7 from RPL and 2 from SML soils) were tested for their ability to degrade PHB. Briefly, one colony of each isolate was transferred to test tubes containing 5 ml of either: 1) MSM or 2) MSM + 3 mM PHB. Uninoculated tubes were included as controls. Tubes were incubated for 3 days at 20 °C with gentle shaking (150 rpm) and growth was measured periodically by UV absorbance at 600 nm (Spectronic 21 spectrophotometer, Bausch & Lomb).

After the 3-day incubation, cultures were analyzed by GC/MS for the production of protocatechuate (aerobic metabolite of PHB degradation) and for the presence of residual PHB. Briefly, 5 mL of each culture was extracted twice with 5 mL of ethyl acetate. Extracts were dried over sodium sulfate and concentrated to 300 µL using a TurboVap LV evaporator (Zymark). Next, 100 µL of extract was derivitized with 25 µL of N,O-Bis(trimethylsilyl)trifluoroacetamide (BSTFA) and 1 µL of sample was injected into a Hewlett-Packard Model 6890 gas chromatograph equipped with an HP-5 fused silica capillary column (5% phenylmethyl silicone, 30 m x 0.25mm x 0.25 μm film thickness; Hewlett-Packard) connected to a Hewlett-Packard Model 5973 quadrupole mass selective detector (operated at an electron energy of 70 eV and a detector voltage of 2000-3000). A splitless injection was used with a 1-min delay before septum purge. Helium was the carrier gas (linear gas velocity of 30 cm/s). The injector and detector temperatures were 250 and 300°C, respectively. The ion source pressure was maintained at 1.0 x 10-5 Torr. The oven temperature program began at 50°C and increased to 250°C at 10 degree/min.

*Genome assembly and functional gene annotation*

Genomic DNA RP11^T^ was sequenced using the capacity of a quarter lane of Illumina MiSeq (2 x 250 bp). The sequencing output was quality preprocessed using Trimmomatic (v. 0.32) [6] and FastX Toolkit (v. 0.7) [7] and assembled using SPAdes (v. 3.10.1) [8]. Prodigal was used to predict open-reading frames (v. 2.6.2) [9]. Auxiliary activity (putatively lignin-degrading) genes were annotated using custom hidden-Markov models for laccases, aryl alcohol oxidases and dye-decoloring peroxidases developed by [10] using hmmscan from HMMER (v. 3.2.0) [11]. Dioxygenases were annotated using Prokka [12] in KBase [13]. All annotations were verified against the non-redundant NCBI database using BLASTp. The completeness of aromatic degradation pathways was assessed using the KofamKOALA annotation pipeline [14]. The average nucleotide identity of RP11^T^ compared to all *Burkholderiaceae* genomes in the NCBI refseq_genomic database (n = 3,991; on June 20^th^, 2019). Average nucleotide identity for all genomes was calculated using FastANI [15].

*Quantifying expression of pobA*

Prior to reverse transcription, DNA was eliminated from RNA extracts by treatment with DNAse I (Invitrogen). Resulting cDNA was PCR reactions contained 1X Power SYBR qPCR Master Mix (Applied Biosystems), 0.5 µM of forward and reverse primer, and 1 µL template. Thermal cycling conditions were as follows: 95°C for 10 minutes, 45 cycles of 95°C for 15 seconds and 58°C for 1 minute, and a 7-min extension at 72°C. A standard curve was generated by diluting PCR amplified pobA from RP11^T^ genomic DNA, spanning 2.76 x 10^2^ to 2.76 to 10^7^ copies per uL. Our *pobA1* primer set covered approximately 40% of the genes present in clade 6 with a maximum of a single base mismatch, while our custom HMMs models (available in Supplementary Data) were 100% accurate and identified six additional homologs missed using BLAST homology searches.

*Discrepancies in ASV Naming*

Due to differences in trimming parameters used in the processing of 16S rRNA gene libraries from the soil priming experiment versus *in situ* DNA-SIP experiment, the ASV identifiers differ between the two datasets. For example, the ASV for P. madseniana RP11^T^ is ‘5b18237cf385326c2cda6c3441ba8d40’ and ‘6b64ff37fff389a42eccf7387ffac8b9’, respectively. The first 5 nt from 515F region of V4 were truncated during the processing of the soil priming amplicon data. The corresponding ASV names for all PHB-degraders is available in Table S3, based on 100% identity across the full amplicon sequences.

*Evolutionary reconstruction of pobA paralogs*

The phylogeny of all *Burkholderiaceae* was determined based on a multi-locus sequence alignment (MLSA). All *Burkholderiaceae* genomes were downloaded from NCBI refseq_genomic on June 20^th^, 2019 (NCBI taxid 119060; n = 3,991). Open reading frames were predicted with Prodigal (v. 2.6.4) [9] and the translated peptide sequences for a common set of 24 single-copy genes were identified using HMMs from BUSCO [16]. Peptide sequences were independently aligned with MAFFT (algorithm: FFT-NS-2 & progressive method) [17] and then concatenated into the MLSA containing 13,366 positions and a total of 3,559 genomes. A maximum likelihood tree was built from the MLSA using RAxML (n_bootstraps_ = 100) with GTR substitution and gamma heterogeneity models [18]. The ‘isolation source’ for each genome was recovered from the NCBI BioSample database.

The phylogeny of all *pobA* encoded by *Burkholderiaceae* was determined using similar methods. *pobA* were recovered from genome according to a DIAMOND BLASTp [19] homology of >= 40% identity across 95% of full-length of the *pobA* identified in *Paraburkholderia madseniana* RP11^T^ (*pobA1* and *pobA2*). An alignment and maximum likelihood tree were built using the previously described software and methods. The tree was divided into seven clades for which HMMs were built using hmmbuild [11] from HMMER with the input MAFFT alignments for each clade. The hmm profiles were concatenated and are provided in the Supplementary Data package. The accuracy and sensitivity of each hmm was validated against all *Burkholderiaceae* genomes. The phylogenetic conservation of *pobA* paralogs was assessed using consenTRAIT based on mapping genomes encoding paralogs onto the MLSA tree and contrasting against the distribution of other dioxygenase genes involved in degrading aromatics: gentisate 1,2 dioxygenase, benzoate 1, 2-dioxygenase, catechol 1, 2-dioxygenase and protocatechuate 4,5-dioxygenase [20]. The phylogenetic dispersion for the same genomic traits was assessed using Purvis and Fritz’s D [21] based on the same MLSA using the ‘phylo.d’ function in the R-package caper [22]. These analysis of phylogenetic conservation and dispersion results are presented in the Supplementary Extended Results section.

*Assessing function of pobA paralogs*

A structural analysis of the *pobA* paralogs in the RP11^T^ genome was performed based on a TCoffee ‘Expresso’ alignment [23] using homologs whose structure and function have been elucidated [24–26]. The analysis was performed to determine whether both contained the correct active and substrate-binding site residues required for p-hydroxybenzoate 3-monooxygenase activity.

*Comparing pobA genes to SIP-lignin study*

A targeted assembly of *pobA* in metagenomes from a related SIP-lignin [10] was performed using megaGTA [27] and our custom *pobA* HMMs (accessions in Table S6). The taxonomy of assembled *pobA* was determined using the Lowest Common Ancestor algorithm implemented by MEGAN [28] based on DIAMOND BLASTp searches against the NCBI ‘nr’ database (downloaded Feb. 3rd, 2017).

*In vitro Soil Priming Experiment*

A soil microcosm experiment was performed to link the activity of phenolic-acid degrading populations with soil priming. Soil treatments were aimed at identifying differences according to substrate: (i) water-only (baseline), (ii) ^13^C-glucose and (iii) ^13^C-PHB, or the role of *pobA* and *Paraburkholderia* sp. RP11^T^ activity: (iv) ^13^C-PHB + *pobA* inhibitor, (v) *pobA* inhibitor-only control, (vi) ^13^C-labeled RP11^T^ cells + dilute ^12^C-PHB and (vii) unlabeled RP11^T^ cells + dilute ^13^C-PHB (overview in Figure S2).

For each treatment, the sum total of mg C per gram dry weight soil required for all replicates was weighed in bulk on an analytical scale (0.1 mg accuracy) and dissolved in the correct volume of water needed to aliquot four replicates. *p*-hydroxybenzoic acid (PHB) and methyl 4-hydroxy-3-iodobenzoate (‘mPHB’, the pobA inhibitor) were dissolved with the assistance of sonication at 60 °C for approximately 30 minutes. A saturated solution of mPHB (0.486 mM) was added to soils, resulting in slight differences in the total mg C of mPHB amended. This was done to maximize inhibitory strength, which was accounted for by supplying the same amount to controls.

The individual amendments of glucose (cat #: G-8270; Sigma-Aldrich) and PHB (cat #: H-3766; Sigma-Aldrich) were diluted to 17.5 atom % ^13^C using corresponding compounds with natural abundance in order to provision enough C for an amendment rate of 0.5 mg C per g dry wt soil. The final isotopic concentration of amended solutions were verified using EA-IRMS at the Cornell Stable Isotope Lab. Prior to EA-IRMA, solutions were diluted with 100 mg/mL of ^12^C-glucose to prevent saturation of the detector. Final atom % ^13^C values for glucose (17.46 % ^13^C) and PHB (17.23 % ^13^C), matching desired labeling concentrations, were obtained.

Soils were amended with *Paraburkholderia madseniana* RP11^T^ cells to isolate its role in priming. Cells were added in solution with dilute ^13^C-labeled PHB (25 µg C per g dry wt. soil) to stimulate PHB-degrading activity. Two treatments were necessary to isolate the mineralization of native SOC (i.e. priming). To account for CO_2_ derived from RP11^T^ biomass, cells were grown in MSM with ^13^C-glucose (99% atom ^13^C: Sigma-Aldrich) as the sole carbon source to label cells. The ^13^C-labeled cells were amended to soil with dilute, unlabeled PHB. Conversely, to account for the respiration of the dilute PHB amendment, unlabeled RP11^T^ cells were grown in MSM with unlabeled glucose and amended with the same ^13^C-PHB amended in the other PHB treated soils, except diluted 1:20. RP11^T^ cultures were grown for 30 hrs on 0.2% glucose (^13^C and ^12^C). Cells were washed by two cycles of pelleting (3,000 rcf for 20 min) and resuspension in saline solution (0.85%). Plate counts were performed on the washed cell prior to equally dividing cell biomass into three portions to treat soil from each ecoplot. Cells were pelleted and resuspended in dilute PHB in a volume corresponding to the desired water holding capacity of each soil. The resuspended cell slurry was incubated for 20 minutes to stimulate activity prior to amendment. A total of 1.8 x 10^7^ cells per g dry wt soil were added for each treatment.
