## Supplementary material for "Phenolic acid-degrading *Paraburkholderia* prime decomposition in forest soil": Wilhelm and DeRito - Supplementary Results.docx

From Wilhelm and DeRito *et al.*, 2020 – Phenolic acid-degrading populations of *Paraburkholderia* prime decomposition in forest soils.

*PobA inhibition of RP11^T^ growth on PHB*

The effective dose of methyl 4-hydroxy-3-iodobenzoate (mPHB), an inhibitor of PobA, was determined by measuring the growth of *P. madseniana* RP11^T^ on PHB in the presence of a dilution gradient of mPHB. The MSM+PHB medium was prepared with varying concentrations of mPHB: 0.486 mM (saturated solution at RT), 0.486 mM x 10^-1^, 0.486 mM x 10^-2^, 0.486 mM x 10^-3^ and 0.486 mM x 10^-4^. Growth was measured by OD. All concentrations of mPHB initially resulted in decreased growth, but only the saturated concentration inhibited growth below detection, relative to uninoculated controls. In all cases, growth was eventually observed with the saturated concentration suppressing growth completely for 4 days.

*Re-analyzing published 16S rRNA gene amplicon libraries*

16S rRNA gene amplicon data from several studies reporting the involvement of ‘*Burkholderia’* in white-rot [1], lignin-degradation [2], soil priming [3–5] or plant roots [6–9] were re-analyzed and determined to consist primarily of *Paraburkholderia* with dominant populations matching to *P. madseniana* RP11^T^ and *P. xenovorans* LB400^T^ in all but [7]. The majority of this information was published in Supplementary Table 5 in Wilhelm *et al.* [10].

*Structure-function relationships of pobA paralogs*

The paralogs of *pobA* in RP11^T^ were distinct and homologous to structurally characterized enzymes from *Pseudomonas CBS3* [11] (67% identity to *pobA1*) and *P. fluorescens* [12] (76% to *pobA2*). Key residues involved in PHB-binding (Ser_212_, Arg_214_ and Tyr_222_) were conserved in both paralogs, but *pobA1* and *pobA2* differed in a residue involved in flavin-binding that can alter affinity for NADPH (Arg_42_) versus NADH (Thr_42_), respectively [13]. We propose several hypotheses about the role of the putative NADPH-consuming PobA, given that NADPH is primarily used in anabolism. One hypothesis is that this NADPH dependent PobA paralog modifies *p*-hydroxybenzoate as part of an uncharacterized biosynthetic pathway. Alternatively, the *pobA* paralog might serve in a *luxR* dependent sense and response system as described for *p*-coumaric acid, which is hypothesized to enable bacterial-plant signalling [14]. RP11encodes *luxI* and *luxR* homologs and endophytic *Paraburkholderia* from poplar roots (isolate BT03) exhibit quorum sensing activity [15]. It is also possible that the NADPH dependent *pobA* paralog might function in detoxification or O_2_ scavenging in the cytoplasm. For example, *Paraburkholderia* are obligate aerobes that are often capable of nitrogen fixation [16], and O_2_ scavenging enzymes could serve to protect nitrogenase from inactivation by O_2_. The use of dioxygenases in O_2_ scavenging has been shown to occur in phenolic acid-rich termite guts [17].

*Evolutionary reconstruction of pobA paralogs*

The existence of *pobA* paralogs in RP11^T^ prompted an investigation into the identity, relatedness and environmental associations of *Burkholderiaceae* encoding paralogs. *Paraburkholderia* were significantly likelier to encode paralogs than all other *pobA*-encoding *Burkholderiaceae* (Fisher’s exact, O.R. = 3.1, p < 0.001). Twenty-five percent of *Paraburkholderia* encoded paralogs compared to the next highest frequency in *Burkholderia* (9%). Paralogs of *pobA* were most common in soil isolates compared to other environmental sources (Fisher’s exact, O.R. = 2.7, p < 0.001; Figure 7A). The phylogenetic depth at which paralogs occurred within related groups of *Burkholderiacaea* was relatively shallow (τ_D_ = 0.005 out of a total depth of 2.5 units) compared to other aromatic degradation genes, such as gentistate 1,2-dioxygenase (τ_D_ = 0.009), benzoate 1,2-dioxygenase (τ_D_ = 0.01) and protocatechuate 3,4-dioxygenase (τ_D_ = 1.0; Table S8). Genomes encoding paralogs were the most phylogenetically clustered of all aromatic degrading genes tested, exhibiting a Brownian mode of trait inheritance (Purvis and Fritz’s D = 0.01) in contrast to the more random dispersion pattern of the protocatechuate dioxygenase gene (D = 0.249; Table S8). Notably, 32% of *pobA*-encoding actinobacterial genomes encoded multiple copies: commonly two (21%) or three (8%), but also up to five in *Streptomyces* and *Amycolatopsis* genomes (~1%). No *Burkholderiaceae* genome encoded more than two copies.

Here is useful context for interpreting values of D. A value of D < 0 suggests a highly clustered trait, D ~ 0 indicates a Brownian motion mode of evolution, D = 1 suggests a random mode of evolution and D > 1 suggests phylogenetic overdispersion. Martiny et al. [18] write “Complex traits encompassing many genes like oxygenic photosynthesis and methanogenesis were extremely clumped (D << 0). Other traits including nitrogen and CO_2_ fixation, anoxygenic photosynthesis and sulfate reduction displayed a clumped distribution consistent with a Brownian motion model of evolution (D ~ 0). Finally, traits like carbon substrate utilization were dispersed in a mode between a Brownian motion and a random model (0 < D > 1), suggesting that the ability to grow on different carbon compounds is quite dispersed”

*Targeted assembly of pobA from published metagenomes*

A targeted assembly of *pobA* was performed on metagenomes constructed from ^13^C-enriched or ^12^C-control DNA derived from forest soil incubations with either ^13^C-lignin or natural abundance ^13^C lignin [2] (methods in Supplementary methods). The assembly yielded 1,204 *pobA* homologs occurring exclusively in metagenomes derived from ^13^C-enriched DNA pools (Table S7). The majority of *pobA* recovered belonged to clade 5 (82%), followed by clade 4 (9.4%) and clade 6 (6.7%). The *pobA* sequences were most related to *Rhizobiales* (44%), *Burkholderiales* (41%), *Sphingomonadales* (10%) and *Pseudomonadales* (4%) according to Lowest Common Ancestry. Six of the most abundant amplicon phylotypes from [2] were identical to the 16S rRNA gene of *P. madseniana* RP11^T^. However, none of the recovered *pobA* were identical to paralogs in RP11^T^ (top % identity according to BLASTp = 71% for *pobA.1* and 75% for *pobA.2*).

*Mechanism of negative priming*

The addition of stimulated RP11^T^ cells to soils was designed to test whether priming would be induced by an increased presence of PHB-degrading bacterium. Contrary to expectations, the addition of RP11^T^ biomass produced a net decrease in respiration of existing SOC of between 5 - 14% relative to controls (i.e. negative priming), which showed no signs of abating by the end on day 26. Negative priming was associated with steep increases in the relative abundance of *Streptomyces* and *Streptacidiphilus* phylotypes (Figure S3C). These populations were below detection in field soil (min_reads_ = 7,318) and increased to upwards of 35% of the total amplicon pool in RP11^T^-amended soil. The trend in their relative abundance was more pronounced in cDNA than DNA pools, indicating their increased representation in active populations at day 7 (Figure S3D).

We attribute the negative priming to the preferential use of RP11^T^ biomass by decomposers in place of soil organic matter. Soils were amended with RP11^T^ cells at a relatively low abundance to mirror a rare taxon in soil (~ 0.001% of total bacteria). Our experiment was the first attempt to use cell biomass in a priming study and the surprising strength of negative priming was unanticipated. We believe the negative priming overshadowed any positive priming which may have occurred due to the addition of stimulated RP11^T^ cells. *Streptomyces* may produce antibiotics or ammonia [19] that lowers net decomposer activity or may increase the net efficiency of organic matter degradation. Their growth in glucose-amended soils was consistent with the expected greater proportion of glucose being converted to biomass. The effect of biomass in stimulating populations that actively diminish the mineralization of SOC suggests the physical ‘entombing’ of biomass [20] is not the only mechanism by which biomass influences SOC cycling.
