## Supplementary figures and images for "Phenolic acid-degrading *Paraburkholderia* prime decomposition in forest soil"

### 3. Rarefaction curves - SIP amplicon libraries.pdf

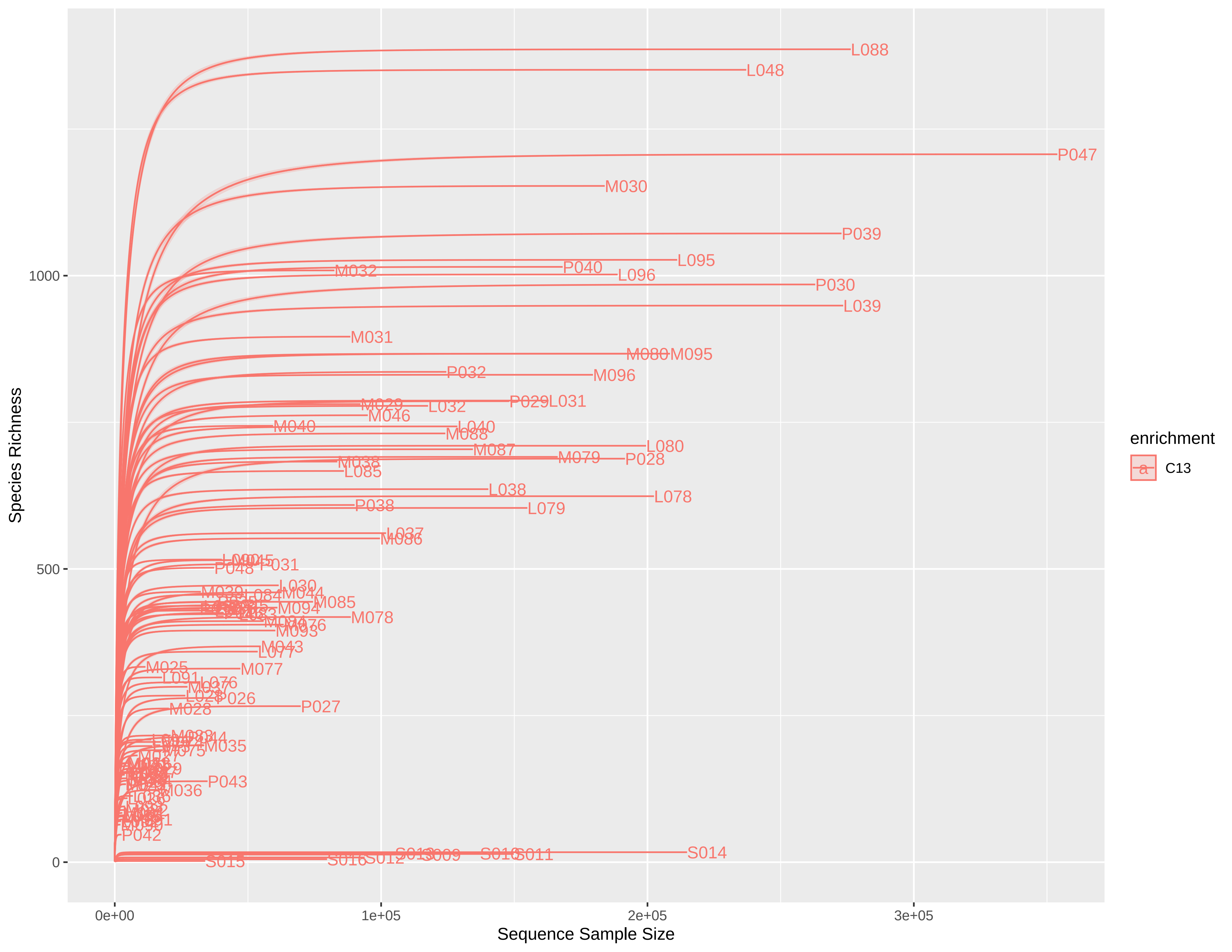
